# Persistent legacy effects on soil microbiota facilitate plant adaptive responses to drought

**DOI:** 10.1101/2024.08.26.609769

**Authors:** Nichole A. Ginnan, Valéria Custódio, David Gopaulchan, Natalie Ford, Isai Salas-González, Dylan H. Jones, Darren M. Wells, Ângela Moreno, Gabriel Castrillo, Maggie R. Wagner

## Abstract

Both chronic and acute drought alter the composition and physiology of soil microbiota by selecting for functional traits that preserve fitness in dry conditions. Currently, little is known about how the resulting precipitation legacy effects manifest at the molecular and physiological levels and how they influence neighboring plants, especially in the context of subsequent drought. We characterized metagenomes of six prairie soils spanning a steep precipitation gradient in Kansas, USA. By statistically controlling for variation in soil porosity and elemental profiles, we identified bacterial taxa and functional gene categories associated with precipitation. This microbial precipitation legacy persisted through a 5-month-long experimental drought and mitigated the negative physiological effects of acute drought for a wild grass species that is native to the precipitation gradient, but not for the domesticated crop species maize. In particular, microbiota with a low-precipitation legacy altered transcription of a subset of host genes that mediate transpiration and intrinsic water use efficiency during drought. Our results show how long-term exposure to water stress alters soil microbial communities with consequences for the drought responses of neighboring plants.

## Introduction

The increasing frequency and intensity of droughts associated with global climate change are threatening plant health and survival in both natural and agricultural ecosystems. However, the ability of soil microbial communities to quickly adapt to environmental shifts^1^ may bolster the resilience of plants and ecosystems to drought stress^2^. Additionally, the cumulative effects of past stress exposure can influence microbial communities’ responses to future environmental challenges, a phenomenon referred to as legacy effects or ecological memory^3^. Despite the growing recognition of microbial legacy effects, little is known about the mechanisms driving them, their long-term persistence, and whether their effects extend uniformly across different plant species.

To investigate microbial drought legacy effects and isolate the drivers and impacts of microbial adaptations to water limitation, we (1) evaluate natural soil metagenome variation across a steep regional precipitation gradient, (2) test the ability of legacy effects to persist through experimental perturbation, and (3) evaluate the impacts of microbiome precipitation history on plant responses to acute drought at the molecular and physiological levels. Finally, we assess the extent to which microbiome legacy effects are transferable across plant species. Our results demonstrate that legacy effects of historical exposure to dry conditions are more salient at the metatranscriptomic level than at the taxonomic or metagenomic levels, and that they trigger transcriptional changes in plant roots that improve resistance to subsequent acute droughts, at least in some plant species.

## Results

### Mineral nutrient accumulation and precipitation impact soil microbiota taxonomic composition

To identify microbial markers of precipitation legacy effects, we sequenced the metagenomes of soils from six never-irrigated remnant prairies spanning ∼568 km of a steep precipitation gradient in Kansas, USA (Fig. 1a,b; Supplementary Table S1). Although bacterial alpha diversity was similar among soils (Extended Data Fig. 1a), community composition showed a strong biogeographic signature (PERMANOVA, R^2=^0.11, *p=*0.001) and precipitation explained 5.3% of the variation. The first principal coordinate axis, which explained 10.6% of the total variation, separated the bacterial communities of the two highest-precipitation sites from the other soils (Fig. 1c). In line with previous findings^4^, Actinomycetota and Bacillota were enriched in low-precipitation soils, whereas Pseudomonadota and Acidobacteriota were enriched in high-precipitation soils (Extended Data Fig. 1b).

**Fig. 1.**
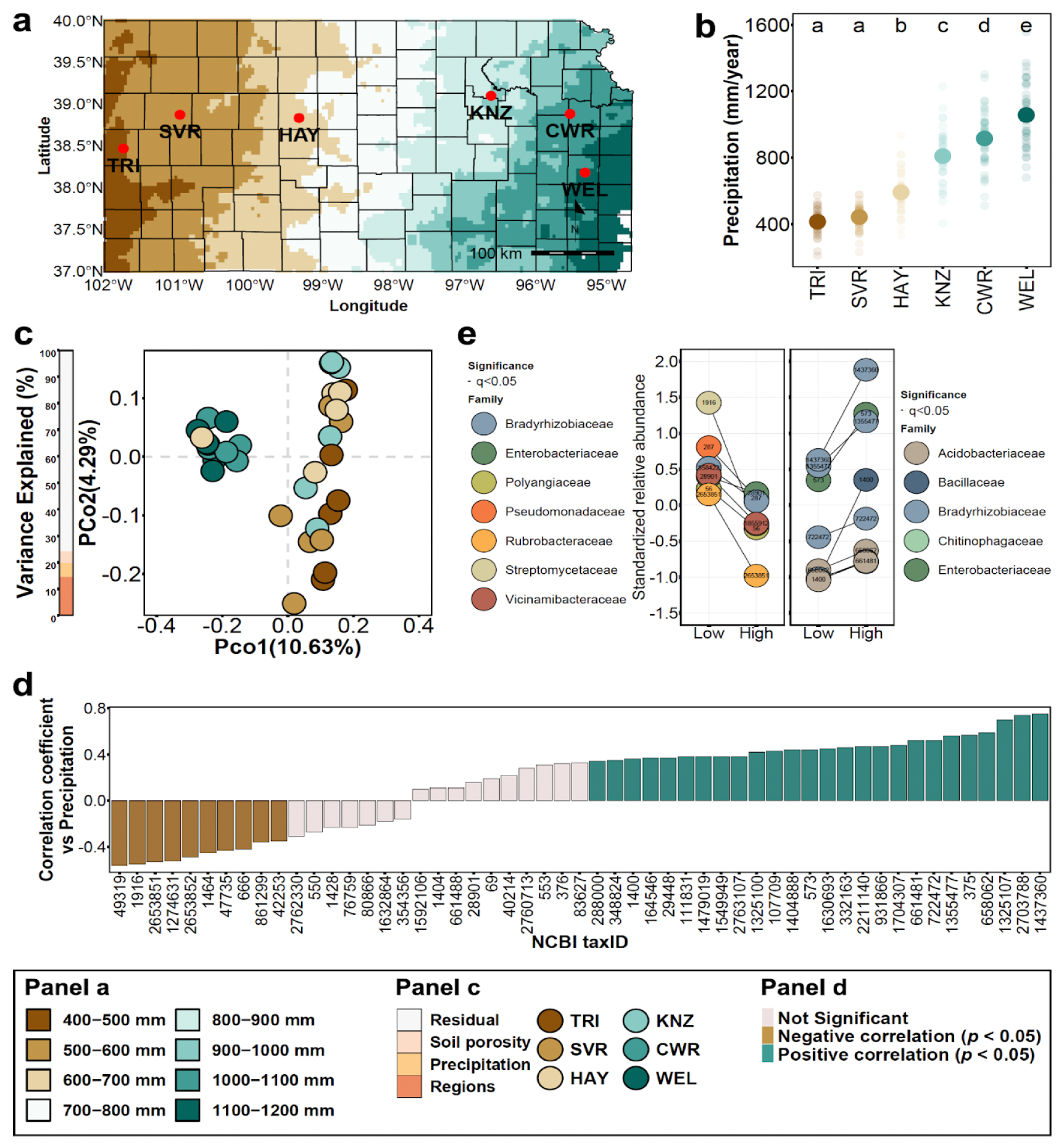
Bacterial markers of precipitation after statistically controlling for other soil properties. a. Map of Kansas, USA, showing the collection locations of the soils used in this work. b. Precipitation (mm/year) at each collection site from 1981 to 2021. Large points represent the mean. Statistical difference between soils was determined via ANOVA followed by Tukey *post-hoc* test (*p*<0.05). c. Principal coordinate analysis of the soil microbiota across the precipitation gradient. The bar on the left denotes the percentage of the overall variance explained by the independent variables. d. Pearson correlation coefficients between each bacterial taxon (NCBI taxID) and mean annual precipitation. e. Relative abundances of bacterial taxa identified in soils exposed to low or high precipitation levels after statistically controlling for soil porosity and mineral nutrient content. Coloured points represent the mean standardized relative abundance for each bacterial taxa. The line connecting both points is the difference between low and high precipitation soils. A black line indicates statistical significance (q < 0.05).

To disentangle the influence of precipitation from co-varying edaphic properties^5^, we examined 24 trace element profiles in each soil using ICP-MS. Nutrients are known drivers of taxonomic composition and functional capacity in soil microbiomes, particularly bacterial communities^6,7^. Mineral nutrient content differed among the six soils, with precipitation explaining 28.6% of the variation. The first principal coordinate axis of the soil mineral nutrient profiles, which explained 39.6% of the total variation, separated the three lower-precipitation sites, from the three higher-precipitation sites (Extended Data Fig. 1c). Concentrations of K, Mg, Ca, Li, and P were negatively correlated with mean annual precipitation, while Cd, Mn, Se, As, Zn, Co, Pb, Rb, Fe, and Cr were positively correlated (Extended Data Fig. 1d-e). The mineral nutrient dissimilarities among soils were correlated with the corresponding microbiota composition dissimilarities (Mantel, r=0.384; *p*-value=1e-04; Extended Data Fig. 1f). This suggests that precipitation patterns might influence the accumulation of mineral nutrients in these soils, and both precipitation and nutrients may impact microbial communities. For example, precipitation can drive mineral weathering and solute production in soils^8^, although this process also depends on many other geochemical and biological factors^9^.

Next, we used X-ray computed tomography to quantify soil porosity of undisturbed soil cores. Soil porosity is directly related to soil hydraulic properties; in general, lower porosity results in lower water retention and infiltration^10^. Therefore, it is a good indicator of how precipitation affects the actual water content in the soil. Consistently, soil porosity decreased with depth to about 3.5 cm before stabilizing, and increased with precipitation (Extended Data Fig. 2a-b). Notably, precipitation might affect soil porosity in surface layers of soils^11^, potentially affecting soil niche properties and, consequently, microbial communities. Nevertheless, we found no correlation between the porosity dissimilarities and the microbiota composition dissimilarities of these soils (Extended Data Fig. 2c), suggesting either that soil porosity is not a key element controlling the overall microbial community composition in these soils, or that the influence of porosity is masked by precipitation legacy.

Finally, to identify taxonomic biomarkers of precipitation legacy, we modelled the relative abundances of bacterial taxa in relation to precipitation levels while controlling for soil porosity and soil elemental composition. This analysis revealed distinct clusters of bacterial taxa whose relative abundances varied significantly along the precipitation gradient (Extended Data Fig. 2d-f). Across the six soils, 19 taxa (NCBI taxIDs) were positively correlated and nine were negatively correlated with precipitation (Fig. 1d). Additionally, 15 of the most abundant (>0.1%) and prevalent (>20%) taxa were enriched or depleted in the three lower-precipitation soils relative to the higher-precipitation soils (Fig. 1e). Together, these results indicate that water availability shapes soil bacterial communities, possibly by selecting for functions necessary to adapt to dry conditions and/or subsequent re-wetting^12^.

### Functional category enrichments and strain-level genetic analysis suggest molecular mechanisms of precipitation legacy effects

We used assembled metagenomic contigs to explore the functional potential of the soil communities spanning the precipitation gradient. To focus on functions that are associated with water availability rather than site-specific variation, we collapsed our six soils into two groups representing our sites with low-precipitation *vs.* high-precipitation histories. These groupings preserved the observed similarities in taxonomic composition, mineral nutrient content, and porosity (Fig. 1c; Extended Data Fig. 1c, 2a). In total, 62 Gene Ontology (GO) categories and 3396 KEGG reactions were differentially abundant between groups (Extended Data Fig. 2g; Supplementary Table S2). Biological processes enriched in low-precipitation soils included nitrogen cycling, fatty acid biosynthesis, DNA repair, and glucan metabolism, all of which have been linked to drought or stress tolerance^13–15^. Additional processes linked to stress responses were depleted in low-precipitation soils, including ion transport (involved in osmotic adjustment), lipid catabolism (relevant for membrane integrity), and metabolism of cellular aldehydes and ketones (involved in oxidative stress)^16,17^ (Extended Data Fig. 2g). The observed differences in functional potential between low-precipitation and high-precipitation sites suggest that these microbiomes are functionally adapted to local precipitation levels, making them excellent candidates for exploring how microbial precipitation legacy affects plant drought tolerance.

Next, we investigated precipitation-associated genetic variation within 33 focal bacterial species, including the previously-identified bacterial biomarkers and other highly abundant and prevalent taxa. Shotgun metagenomic reads were mapped to reference genomes from the NCBI database, and sequence variants were identified. Analysis of genetic distances showed variations in strain-level microbiome structure across the precipitation gradient (Extended Data Fig. 3a) and between precipitation levels (Extended Data Fig. 3b).

Subsequently, we conducted a genotype-environment association analysis and identified genetic variants associated with mean annual precipitation in several bacterial lineages, including *Streptomyces*, *Luteitalea*, *Rubrobacter*, *Lacibacter*, and *Rhizobium*, and three *Bradyrhizobium* lineages (Extended Data Fig. 3c, Supplementary Table S3). Most of the associated variants were located within or near protein-coding regions. Notably, some of the corresponding genes have known adaptive functions such as the phenolic acid stress response (PadR family transcriptional regulator)^18,19^, maintenance of cellular functions under iron starvation and oxidative stress (Fe-S cluster assembly protein SufD and SufB)^20^, and fatty acid synthesis (acetyl-CoA carboxylase biotin carboxylase subunit)^21^, which impacts membrane composition and stress tolerance^17^ (Extended Data Fig. 3c; Supplementary Table S3). These results indicate that precipitation legacy effects manifest through genetic differentiation within bacterial species, not just variation in community composition. Further study of these variants could reveal mechanisms by which precipitation shapes soil microbiota over ecological and evolutionary time.

### Metagenomic and especially metatranscriptomic precipitation legacies are resilient to short-term perturbations

To assess the effects of short-term perturbations on soil microbiome legacy, we exposed the six focal soils to a five-month-long conditioning phase experiment. Replicate pots of each soil were either left unplanted or planted with a seedling of *Tripsacum dactyloides* (eastern gamagrass, which is native to Kansas), and were either drought-challenged or well-watered, in a factorial design (Extended Data Fig. 4a). Compared to well-watered plants, the droughted plants were shorter and had more root aerenchyma (Fig. 2a; Supplementary Fig. 4a,b), confirming that the conditioning phase drought treatment was severe enough to induce stress responses in the plants, and presumably in the microbes^22^.

**Fig. 2.**
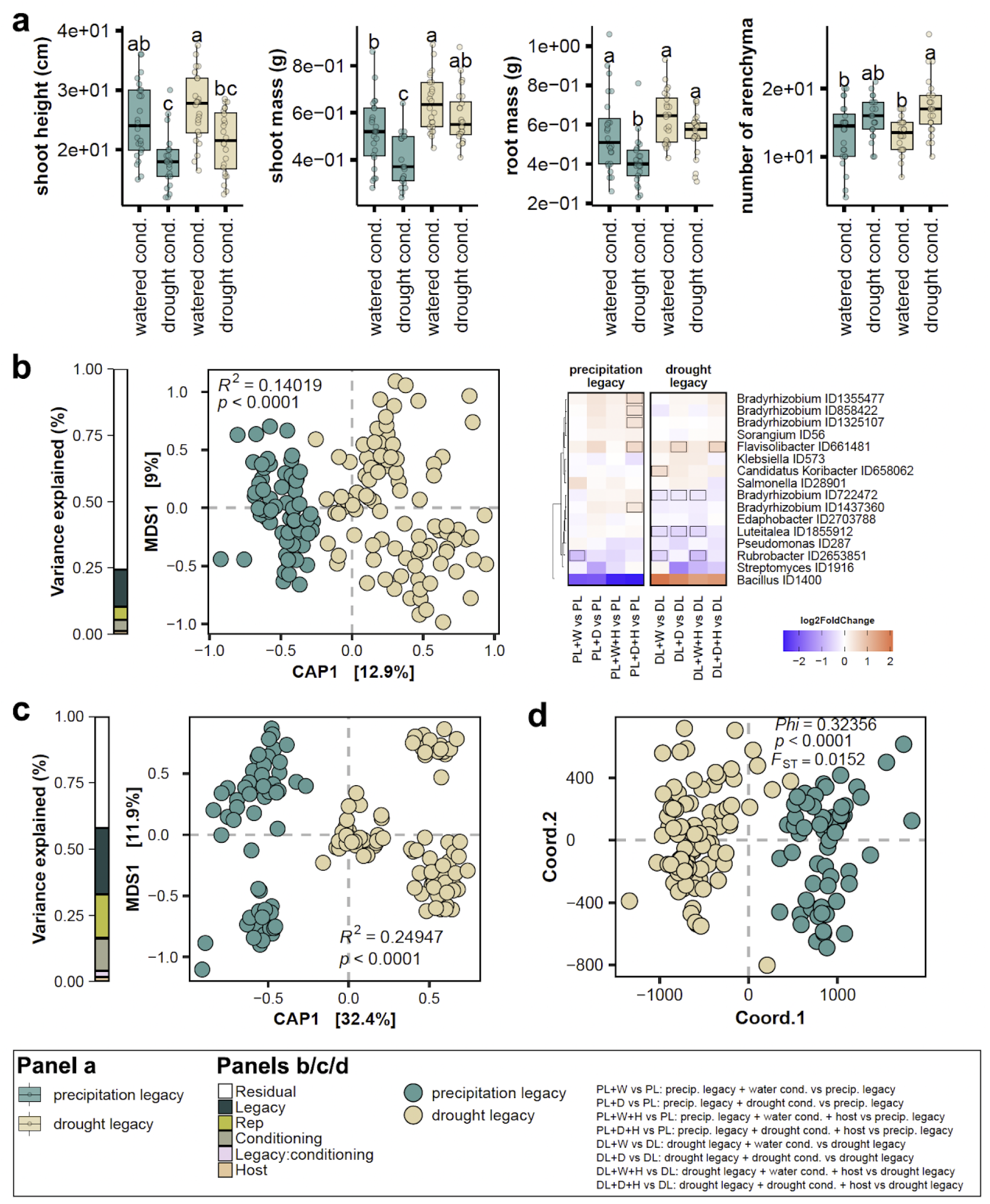
Precipitation legacy effects on the soil microbiota are resilient to short-term perturbations. **a.** Box plots showing phenotype distributions of *Tripsacum dactyloides* plants grown during the conditioning phase, in which they experienced five months of drought (+D) or well-watered (+W) conditions and were grown in soils with either low-precipitation legacies (“DL”, brown) or high-precipitation legacies (“PL”, blue) (for additional phenotypes, see Extended Data Fig. 4b,d). PL and DL indicate the baseline soils prior to the initiation of conditioning phase treatments. Box edges represent the first and third quartiles; whiskers indicate the range of data points that fall within 1.5 times the interquartile range of the first and third quartiles; the center lines indicate the medians. **b.** Left: Constrained ordination of soil metagenome taxonomic composition after the conditioning phase treatments. Right: enrichment patterns of precipitation biomarker taxa in response to the different treatments, relative to the pre-conditioning baseline. Rectangles outlined in black indicate bacterial markers that were significantly enriched (red) or depleted (blue) (q < 0.1). **c.** Constrained ordination of soil metatranscriptome content after the conditioning phase treatments. **d.** Principal coordinates analysis of standardized pairwise genetic distances calculated from SNPs in the genomes of the precipitation biomarker taxa. For panels b-d, note that even after five months of experimental perturbation, there is a clear separation of the samples on the first axis based on the precipitation legacy.

Next, we explored how precipitation legacy affected the microbial communities’ responses to intermittent drought and watering events. Congruent with observations from the field-collected soils, water availability did not affect bacterial community alpha diversity regardless of whether a plant was present (Extended Data Fig. 4b). Although phylum-level taxonomic profiles were also similar across treatments, constrained ordination of metagenomic sequences indicated that the conditioning phase watering and host treatments explained 4.3% and 1.1% of the variation in metagenome content, respectively (Extended Data Fig. 4c-d). In contrast, precipitation legacy explained 14.1% of the metagenome variation, confirming that the conditioning phase treatments did not erase the ecological memory of these soils (Fig. 2b). Additionally, we still detected significant taxonomic differences between high- and low-precipitation soils (Supplementary Fig. 2a,b) and the previously-identified taxonomic markers of drought legacy (Supplementary Table S4) generally followed the expected enrichment and depletion patterns, regardless of conditioning phase treatment. This resilience was particularly evident in soils with low-precipitation legacies (Fig. 2b, Supplementary Fig. 2b).

To assess whether precipitation legacy effects on strain-level genetic variation were also robust to experimental perturbation, we investigated genetic variants within the identified bacterial markers and other prevalent species across the samples, as described previously. A principal coordinate analysis of standardized pairwise genetic distances based on those variants revealed that samples clearly separated along the first axis based on precipitation legacy rather than the experimental treatments (Fig. 2c). Furthermore, our experimental perturbations did not affect allele frequencies (Supplementary Fig. 3a,b) whereas a genome-environment association analysis identified precipitation-associated genes in *Luteitalea* and two *Bradyrhizobium* lineages, recapitulating results from the pre-conditioning phase soils (Supplementary Fig. 3c, Supplementary Table S5).

Next, we tested whether functional potential still differed between low- and high-precipitation soils after 5 months of experimental perturbation. GO terms related to the nitrogen cycle metabolic process, including purine−containing compound metabolic process and pyrimidine−containing compound metabolic process, were enriched in dry-legacy soils, while GO terms related to ion transport and amino acid catabolic process were depleted in the dry-legacy soils, regardless of conditioning treatment (Supplementary Fig. 4a, Supplementary Table S6). These enrichment patterns mirrored the original field soil observations: of the 62 GO categories that were associated with precipitation legacy in the pre-conditioning soils, 49 retained the same pattern after five months of experimental drought, and 50 did so after five months of ample watering (Extended Data Fig. 2g, Supplementary Table S7). These results show that the legacy effect of precipitation on the functional capacity of soil metagenomes was robust to the conditioning phase perturbations.

Metagenome data often includes unexpressed genes and sequences from dormant or dead organisms, which could exaggerate the robustness of soil legacy effects. Therefore, we also quantified metatranscriptomes from the same samples to focus on biologically active processes across the treatments and soils. Constrained ordination showed that precipitation history explained 24.9% of the variation in microbial gene expression while conditioning phase drought and host treatments explained only 12.3% and 1.8% of the variation, respectively (Fig. 2d; Extended Data Fig. 5a). Furthermore, metatranscriptome-based taxonomic enrichment patterns confirmed the metagenome-based results: even after five months of experimental perturbation, transcriptionally active Actinomycetota and Bacillota remained enriched in low-precipitation soils while Acidobacteriota, Planctomycetota, and Pseudomonadota remained enriched in high-precipitation soils (Extended Data Fig. 5b). In general, the differences in transcriptionally active bacterial taxa between low- and high-precipitation soils mirrored our previous taxonomic-level observations, regardless of the conditioning treatments (Supplementary Fig. 2a; Extended Data Fig. 5c, 6a).

Notably, the metatranscriptome analysis revealed GO categories and KEGG reactions that remained enriched in low-precipitation soils after five months of experimental perturbation, such as tetrapyrrole metabolic process, response to osmotic stress, liposaccharide metabolic process, heme metabolic process, and trehalose catabolic process (Extended Data Fig. 6b; Supplementary Table S8-S9). These results confirm that precipitation legacy strongly shapes both functional potential and gene expression in soil microbiota, and remains robust to perturbation (*e.g.,* five-month-long acute drought). The functional resilience of the soil microbiota creates the potential for microbial legacy effects to influence host responses to environmental changes in natural ecosystems, conceivably enhancing plant resilience to future droughts.

### Microbiome precipitation legacy alters host plant responses to subsequent drought despite taxonomic convergence of root microbiomes

During the conditioning phase, we observed that droughted *T. dactyloides* plants grown in low-precipitation-legacy soils were larger and produced more aerenchyma compared to plants grown in high-precipitation-legacy soils (Fig. 2a). To confirm that these effects of soil precipitation legacy were conferred by the microbiome rather than by co-varying soil properties, we extracted the microbial communities from the conditioning phase pots and used them to inoculate a new generation of *T. dactyloides* seedlings (Extended Data Fig. 4a). These “test phase” plants were divided between well-watered control conditions (N=100) and a drought treatment (N=200; Extended Data Fig. 4a), which we confirmed was severe enough to impair plant growth (Supplementary Fig. 5a). We uprooted five-week-old gamagrass plants for phenotyping and sampled crown roots, which are highly active in water acquisition^23,24^, for 16S rRNA gene microbiome profiling and RNA-seq analysis. We focused on bacterial microbiomes because of evidence that fungi from these soils were insensitive to drought^25^, and because bacterial sequences accounted for 93.63% of metagenomic reads in field-collected soils and 89.99% in conditioning-phase soils (Supplementary Fig. 6a,b). To best capture the precipitation gradient extremes, we measured gene expression only in plants inoculated with microbiota derived from the two lowest-precipitation and two highest-precipitation soils.

Precipitation legacy affected neither alpha diversity nor taxonomic composition of the gamagrass root microbiome (Fig. 3a). Bacterial taxonomic composition was impacted by the test phase drought treatment (ANOVA.CCA, F_1,139_=2.58, p=0.003) but not inoculum precipitation legacy (ANOVA.CCA, F_1,139_=0.89, p=0.55) (Fig. 3b); together these factors explained 2.2% of root microbiome variation (ANOVA.CCA, F_2,139_=1.74, R^2^=0.022, p=0.01). To assess root microbiome stability, we calculated beta-dispersion, a measure of within-group variability, and found it to be equal among treatment groups (Fig. 3c; ANOVA, F_1,144_=0.57, p=0.45). Together, these results suggest that gamagrass exerts a strong homogenizing influence during root microbiome formation, resulting in a stable and drought-resistant microbiota. Because eastern gamagrass shares a long evolutionary history with Kansas soil microbes^26^, this result supports previous findings that co-evolution promotes stable community assembly^27,28^.

**Fig. 3.**
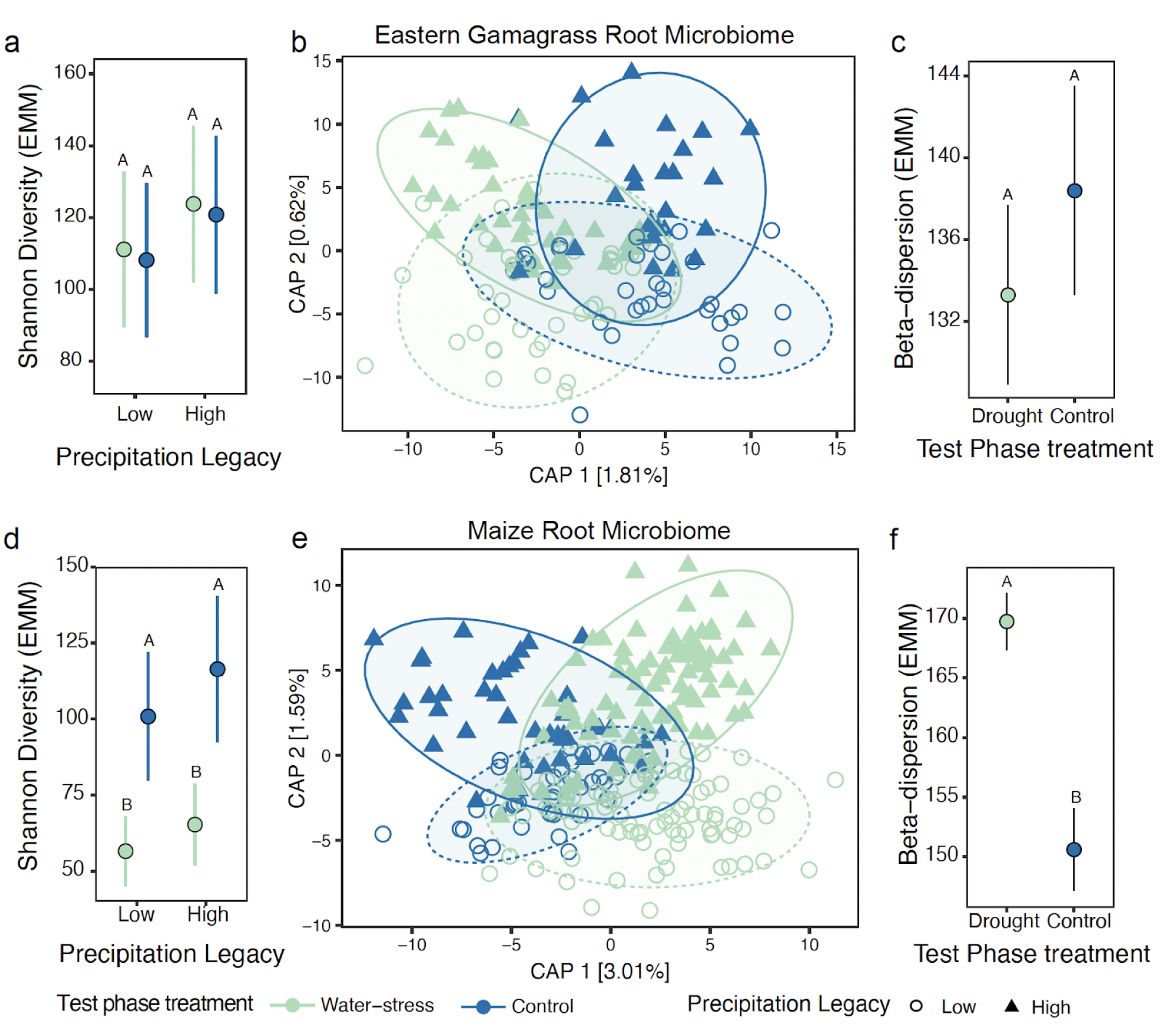
Root bacterial microbiome composition is more stable in *T. dactyloides* (eastern gamagrass) than *Z. mays* (maize) plants exposed to varying inocula and water treatments. For panels a,c,d,f the points are estimated marginal means (EMMs); error bars represent standard errors; letters indicate statistical contrasts (ANOVA with Tukey’s post hoc test *p*<0.05). **a.** Shannon diversity (*e*^Shannon index) was not impacted by precipitation legacy or test phase water treatment. **b**. Ordination constrained by test phase water treatment and inoculum precipitation legacy (ANOVA.CCA, full model F_2,139_=1.74, R^2^=0.022, p=0.01) indicates that test phase water treatment (ANOVA.CCA by term, F_1,139_=2.58, p=0.003), but not legacy (ANOVA.CCA by term, F_1,139_=0.89, p=0.55) significantly impacted the *T. dactyloides* root bacterial microbiome composition. Ellipses represent the 95% confidence intervals. **c.** Drought treatment did not impact the within-group variation (β-dispersion) of the *T. dactyloides* root microbiome. **d**. *Zea mays* (maize) root microbiome Shannon diversity was significantly impacted by test phase water treatment, but not the precipitation legacy of the inoculum. **e**. Ordination constrained by test phase water treatment and inoculum precipitation legacy (ANOVA.CCA, F_2,255_=6.156, R^2^=0.041, p=0.001) indicates that legacy (ANOVA.CCA by term, F_1,255_=4.2538, p=0.001) and test phase water treatment (ANOVA.CCA by term, F_1,255_=8.058, p=0.001) significantly impacted the *Z. mays* root bacterial microbiome composition. Ellipses represent the 95% confidence intervals. **f.** Acute drought treatment increased *Z. mays* root microbiome within-group variation.

Although precipitation legacy did not shape *16S rRNA* gene diversity within the *T. dactyloides* root microbiome, its clear effect on the metagenome and metatranscriptome (Fig. 2c,d) suggested high potential to influence plant phenotype. Indeed, the gene expression profiles in crown roots of plants inoculated with dry-legacy microbiota were distinct from those of plants inoculated with wet-legacy microbiota. Fifteen *T. dactyloides* genes were differentially expressed in plants receiving a high-precipitation *vs.* low-precipitation inoculum (Fig. 4a; Supplementary Table S10). Furthermore, inoculum precipitation legacy influenced the responses of 183 *T. dactyloides* genes to acute drought (Fig. 4b). Of these, 55% were unresponsive to drought in plants grown with a dry-legacy microbiota but were down-regulated or up-regulated in plants grown with a wet-legacy microbiota (Fig. 4b, gene sets I and II, respectively; Supplementary Table S10). This strongly suggests that soil microbiota from low-precipitation sites tend to dampen the transcriptional response of gamagrass to acute drought. For instance, 50 *T. dactyloides* genes were downregulated due to the drought treatment, but only in plants that had been inoculated with high-precipitation-legacy microbiota (Fig. 4b gene set I). These included five orthologs of maize genes predicted to be involved in ethylene- or ABA-mediated signalling of water stress (*Td00002ba004498*, *Td00002ba024351*, *Td00002ba011993*, *Td00002ba005402*, *Td00002ba000033*), and a heat shock protein linked to temperature stress (*Td00002ba042486*) (Supplementary Table S10). Notably, the latter three genes were not identified as drought-sensitive when averaging across inocula, demonstrating that the microbial context is necessary for a complete understanding of molecular plant drought responses. We note that these signatures of microbiota precipitation legacy on host gene expression are averaged across all treatments applied during the conditioning phase (Extended data Fig. 4a), again confirming that microbiome legacy effects are robust to short-term perturbations.

**Fig. 4.**
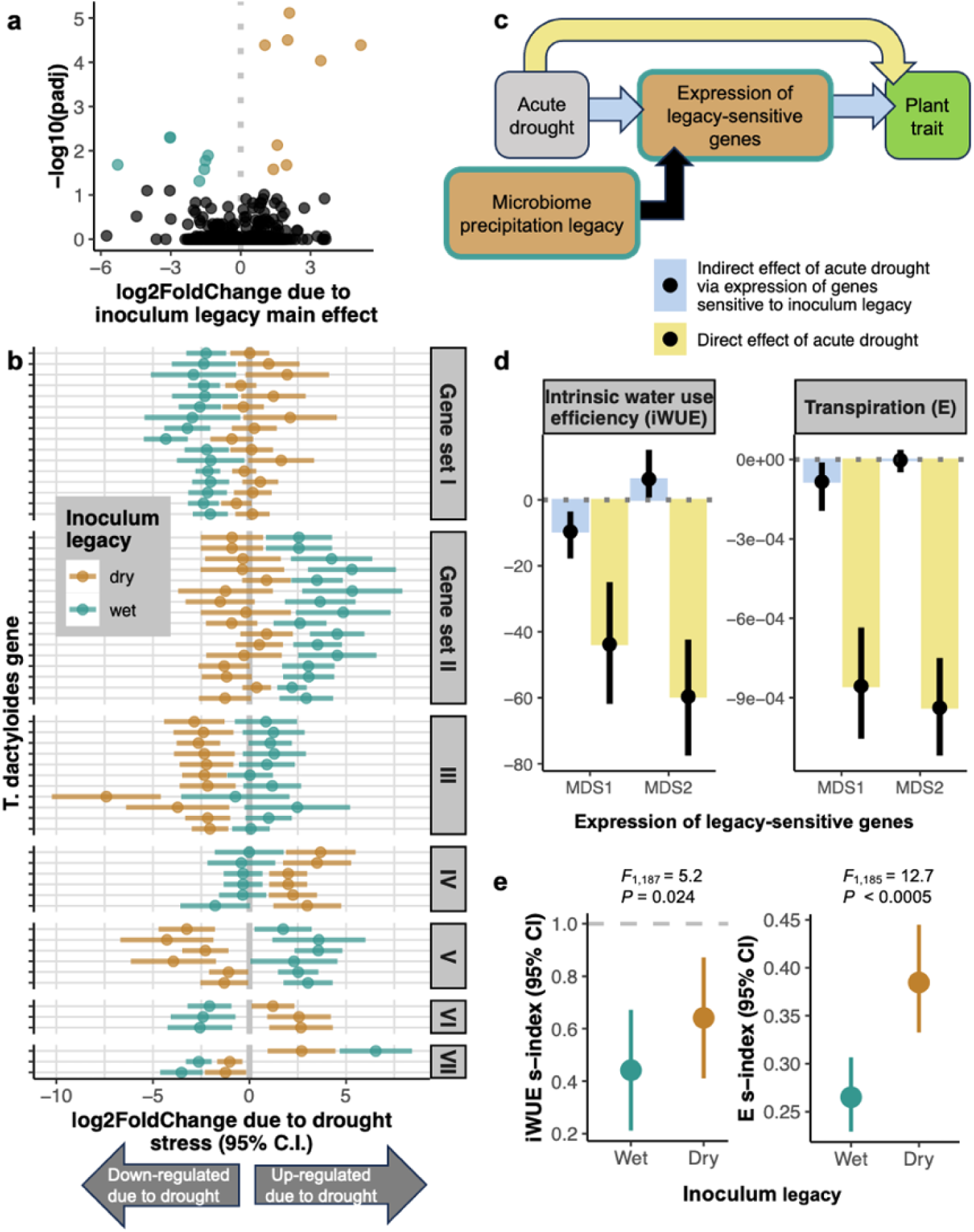
The precipitation legacy of the microbial inoculum mediates transcriptional and physiological responses of *T. dactyloides* (eastern gamagrass) to acute drought. a. Fifteen genes were differentially expressed between plants inoculated with low-precipitation-legacy microbiota *vs.* high-precipitation-legacy microbiota. **b.** In total, 183 *T. dactyloides* genes responded to drought in a manner that was dependent on the drought legacy of the soil microbiota (the inoculum legacy*drought treatment interaction; Supplementary Table 10). Only genes with a drought response of ≥4-fold and with annotated maize ortholog(s) are shown for illustration purposes. Each pair of points represents one gene; the position of each point illustrates how the gene’s transcription level responded to drought treatment in plants inoculated with low-precipitation-legacy (brown) or high-precipitation-legacy microbiota (turquoise). Genes sets correspond to patterns of how inoculum legacy altered their drought responses. **c.** The model used for mediation analysis to test whether the expression of the 198 legacy-sensitive genes (a-b) contributed to the overall effect of acute drought on plant phenotype. **d.** Mediation analysis confirmed that the expression of legacy-sensitive genes (summarized in two dimensions, MDS1 and MDS2) is involved in drought-induced decreases in intrinsic water-use efficiency (iWUE, units: μmol/mol) and transpiration (E, units: mol m^-2^ s^-1^). Yellow bars indicate the “direct” effect of the drought treatment on trait values, i.e., the portion of the trait response that is independent of the transcription levels of microbiota legacy-sensitive genes. Blue bars show the portion of the trait response that is mediated by microbiota legacy-sensitive genes. Error bars: 95% confidence intervals. **e.** Low-precipitation-legacy microbiota (brown) stabilized iWUE and E during acute drought. The s-index describes trait values scaled by the mean value of well-watered control plants, such that an s-index of 1 indicates that droughted plants were phenotypically identical to non-droughted plants. Points are EMMs and error bars indicate 95% confidence intervals.

Next, we assessed whether microbial precipitation legacy altered host physiological and morphological drought responses. For each of 63 traits, we calculated a drought susceptibility index (S-index) that measures the stability of a trait under drought relative to control plants that received the same inoculum type (Supplementary Table S11). We used a random forest model to identify the ten most important traits for describing the phenotypic effects of microbiome precipitation legacy (Supplementary Fig. 7a,b). Redundancy analysis revealed that microbiome precipitation legacy explained 5% of the total variation in host phenotypic response to acute drought (ANOVA.CCA, F_1,187_=10.1, R^2^=0.05, *p*=0.001; Extended Data Fig. 7a). Microbiota precipitation legacy impacted six of the top eight non-collinear traits: transpiration, leaf chlorophyll content, maximum root width, metaxylem area, median root diameter, Cu, K, and intrinsic water use efficiency (iWUE)–an important plant drought response trait^29,30^ (Extended Data Fig. 7b-j).

To solidify the mechanisms linking microbiome legacy to plant phenotype, we used mediation analysis to determine whether phenotypic drought responses were mediated by expression patterns of the 198 genes that were responsive to inoculum precipitation legacy (Fig. 4a-c), summarized in two dimensions (MDS1 and MDS2). MDS1 mediated 18% of the total drought-induced decrease in iWUE (*p* < 0.001), and 8.9% of the total decrease in transpiration (*p*=0.018; Fig. 4d). In contrast, MDS2 mediated 11% of the iWUE response to drought (*p=*0.03), but in the opposite direction, meaning that the activity of these genes counteracted the negative effects of acute drought on iWUE. Four of the top eight genes with the strongest positive loadings on MDS2 were orthologs of the maize gene *ZmNAS7* (nicotianamine synthase); two of these were also in the top 5% of genes with the strongest negative loadings on MDS1 (Extended Data Fig. 8), indicating that they stabilize iWUE during drought. All four *ZmNAS7* orthologs were up-regulated in response to drought, but only in plants that were inoculated with dry-legacy microbiota (Fig. 4b, gene set IV), suggesting a possible mechanism by which low-precipitation-legacy microbiota confer drought tolerance. Consistent with this, salicylic acid–a key regulator of root microbiome assembly^31^–is reported to induce expression of *ZmNAS7*^32^. Although nicotianamine is best known for its role in metal transport, the overexpression of *NAS* genes has conferred drought tolerance, including maintenance of photochemical efficiency at pre-drought levels, in rice^33^ and in the grass *Lolium perenne*^34^. Together, these results show that microbiomes with a low-precipitation legacy improve the drought response of eastern gamagrass.

### Beneficial microbiome legacy effects do not extend to the crop species *Zea mays*

To explore whether the observed microbial legacy effects were transferable across hosts, we simultaneously applied the same inocula to seedlings of maize (*Zea mays* ssp. *mays*), a relative of eastern gamagrass that is non-native to Kansas (Extended Data Fig. 4a). Unlike eastern gamagrass, the maize root microbiome retained taxonomic signatures of both the inoculum precipitation legacy (Fig. 3e., ANOVA.CCA, F_1,255_=4.25, p=0.001) and the drought treatment (ANOVA.CCA, F_1,255_=8.06, p=0.001) (Fig. 3e), which together explained 4.1% of the variation in the data (ANOVA.CCA, F_2,255_=6.16, R^2^=0.041, p=0.001). Also unlike in gamagrass, drought treatment significantly increased beta-dispersion and decreased alpha diversity (Fig. 3d,f; ANOVA F_1,262=_18.17, *p*<0.001), indicating lower microbiome stability.

Next, we identified amplicon sequence variants (ASVs) that were differentially abundant between droughted *vs.* control plants in a manner that depended on the precipitation legacy of the inoculum. These included only two ASVs in gamagrass roots (*Azospirillum* sp. and *Enterobacteriaceae*), but 100 ASVs in maize roots (Supplementary Fig. 8a-b, Supplementary Table S12). Again, this result indicates that gamagrass root microbiomes are more stable–*i.e*., they experience less change in microbiome taxonomic composition and assembly–than those of maize, in response to varying water availability and inoculation with different starting communities.

RNA-seq analysis identified four sets of orthologous genes that were sensitive to microbiota legacy in both maize and gamagrass, but none showed congruent drought responses (Supplementary Note 1). In maize, 23 genes were up-regulated plants inoculated with dry-legacy microbiota, relative to those that received wet-legacy inocula (Extended Data Fig. 9a). These included six defense-related genes, particularly related to jasmonic acid signalling, suggesting that low-precipitation-legacy microbiota were perceived as pathogens by maize but not gamagrass. Other notable functions that were up-regulated by the dry-legacy inocula include root development and iron acquisition (Supplementary Table S10). Additionally, 109 maize genes responded to drought treatment in a manner that was dependent on inoculum precipitation legacy. Only 30 of these were identified as drought-responsive when averaging across inocula (Extended Data Fig. 9b), reinforcing the importance of microbial context for understanding plant drought responses. Like in gamagrass, most of these genes were drought-responsive only in wet-legacy-inoculated or only in dry-legacy-inoculated plants; relatively few reversed the direction of their drought response (Extended Data Fig. 9c). For example, 30 genes were up-regulated in response to drought only in plants grown with wet-legacy microbiota (Extended Data Fig. 9c, gene set I). These genes included several related to pathogen defense (*Zm00001eb116230*, *Zm00001eb150050*, *Zm00001eb222540*) and response to symbiotic fungi (*Zm00001eb033580*), suggesting that soils in high-precipitation regions contain both harmful and beneficial microorganisms and that interactions with both groups are activated under water deprivation in maize (Supplementary Table S10). Notably, one of the genes that reversed its drought response depending on the inoculum’s precipitation legacy was *TIP3*, which encodes an aquaporin that regulates water storage in the vacuole^35^. In plants inoculated with wet-legacy microbiota, *TIP3* expression increased 7-fold due to drought, but decreased nearly 3-fold in plants inoculated with dry-legacy microbiota (Extended Data Fig. 9c, gene set VI). Averaged across the two inocula, however, *TIP3* was not differentially expressed between droughted and well-watered plants; thus, its role in drought response was apparent only when accounting for microbiota precipitation legacy.

Finally, we investigated whether the beneficial effects of microbiome legacy on gamagrass phenotype (Fig. 4d-e) were also conferred on maize. Overall, microbiome precipitation legacy explained only 2.0% of the phenotypic variation in maize (Extended Data Fig. 15a., ANOVA.CCA, F_1,175_=3.72, R^2^=0.02, *p*=0.001), a weaker effect than observed in eastern gamagrass. In general, we observed no significant phenotypic differences attributable to microbiome legacy (Extended Data Fig. 10b,c). Thus, the capacity to integrate the beneficial effect of soil drought legacy is host-specific and may be related to the ability of the host to maintain a stable microbiota in response to drought stress.

## Discussion

Microbially-mediated legacy effects of water limitation can affect plant performance during subsequent droughts^36,37^. Our results reveal how these legacy effects manifest in terms of microbiota taxonomic composition, functional potential, gene expression, and strain-level genetic variation, and how they affect host responses to water-scarce conditions. We identified bacterial taxa and numerous microbial genes and functional pathways that are associated with mean annual precipitation, including those suggesting an increased nitrogen-metabolizing and DNA repair capacity in dry-legacy microbiota. Thus, we provide evidence for the ecological and molecular mechanisms involved in the formation of soil drought memory and demonstrate its robustness to a five-month perturbation, which could mimic a particularly dry or wet season.

We have shown that these metagenomic precipitation legacies have significant implications for plant response to drought. Inoculation with dry-legacy microbiota altered the transcription of key genes controlling plant drought responses. In the prairie grass *Tripsacum dactyloides*, dry- and wet-legacy soil microbiomes give rise to taxonomically similar root bacterial communities yet have strikingly different effects on plant gene expression and phenotypes during a subsequent acute drought. These benefits largely did not extend to maize, which also had a relatively unstable root microbiome and reduced physiological and morphological sensitivity to soil microbiome legacy during drought. However, further research is needed to confirm whether root microbiome stability is a mechanism of adaptive plant responses to drought. Importantly, differences between gamagrass and maize responses to microbiome history indicate that crops may not reap the same benefits as native plant species from potentially beneficial microbial communities. Therefore, our discoveries significantly contribute to our mechanistic understanding of microbiome drought legacy effects, their resistance to perturbation, and their role in plant drought responses, with implications for agricultural and ecosystem management.

## Online Methods

No statistical methods were applied to predetermine sample size. The experiments were randomized, and investigators were blinded to allocation during experiments and outcome assessment.

### 1. Legacy phase: Characterization of soils across precipitation gradients

#### 1.1 Region selection and soil collection

Soils were sampled from six never-irrigated native prairies across Kansas, USA in October 2020. This selection includes eastern Kansas tallgrass prairies: Welda Prairie (WEL), Clinton Wildlife Reserve (CWR), and Konza Wildlife Reserve (KNZ), as well as western Kansas shortgrass prairies: Hays Prairie (HAY), Smoky Valley Ranch (SVR), and Kansas State University’s Tribune Southwest Research Center (TRI). The GPS coordinates of each collection site are available in the Supplemental Table S1. For soil collection, each site was split into three subplots. In each subplot, the surface soil (≈10 cm) containing thick plant root masses was removed with a bleach-sterilized metal shovel. Then, approximately 2.5 L of soil was collected from each site and pooled into a clean plastic bag, for a total of ≈7.5 L of soil collected per geographical location. Soil was held at room temperature for transport back to the laboratory where it was then stored at 4°C until use in the growth chamber experiments (∼ 1 month).

Sub-samples for downstream metagenomic sequencing and nutrient mineral content analyses were air-dried in sterile plastic trays at room temperature for 1 week and then sieved using a 2 mm sieve to remove rocks, big soil particles, and vegetable debris. The sub-samples were shipped to the University of Nottingham for further processing, and all sub-samples were stored at 4 °C until use (∼3 days).

#### 1.2 Precipitation data collection

Daily precipitation from 1981 to 2021 was extracted from the NASA POWER database based on the latitude and longitude of each site^38^.

#### 1.3 Soil elemental content analysis

The soil mineral nutrients and trace elements profiles were determined using inductively coupled plasma mass spectrometry (ICP-MS). The soil samples were dried using plastic weighing boats in the fume hood for approximately 72 h at ambient temperature. Five grams of soil were weighed in 50 mL conical tubes with a four-decimal balance, and treated with 20 mL of 1 M NH_4_HCO_3_, 5 mM diamine-triamine-penta-acetic acid (DTPA), and 5 mL 18.2 MΩcm Milli-Q Direct water (Merck Millipore), for 1 h at 150 r.p.m. in a rotary shaker (adapted from^39^) to extract available elements. Each treated sample was gravity-filtered through a quantitative filter paper (Whatman 42-WHA1442070) to obtain approximately 5 mL of filtrate. Prior to the digestion, 20 µg/L of Indium (In) was added to the nitric acid Primar Plus (Fisher Chemicals) as an internal standard for assessing error in dilution, variations in sample introduction and plasma stability in the ICP-MS instrument. Next, 0.5 mL of the soil filtrates were open-air digested in glass Pyrex tubes using 1 mL of concentrated trace metal grade nitric acid spiked indium internal standard for 2 h at 115 °C in dry block heater (DigiPREP MS, SCP Science; QMX Laboratories, Essex, UK). After cooling, the digests were diluted to 10 mL with 18.2 MΩcm Milli-Q Direct water and elemental analysis was performed using an ICP-MS, PerkinElmer NexION 2000 equipped with Elemental Scientific Inc 4DXX FAST Dual Rinse autosampler, FAST valve and peristaltic pump. The instrument was fitted with PFA-ST3 MicroFlow nebulizer, baffled cyclonic C3 high sensitivity glass spray chamber cooled to 2°C with PC3X Peltier heated/cooled inlet system, 2.0 mm i.d. quartz injector torch and a set of nickel cones. Twenty-three elements were monitored including following stable isotopes: ^7^Li, ^11^B, ^23^Na, ^24^Mg, ^31^P, ^34^S, ^39^K, ^43^Ca, ^52^Cr, ^55^Mn, ^56^Fe, ^59^Co, ^60^Ni, ^63^Cu, ^66^Zn, ^75^As, ^82^Se, ^85^Rb, ^88^Sr, ^98^Mo, ^111^Cd, ^208^Pb and ^115^In. Helium was used as a collision gas in Kinetic Energy Discrimination mode (KED) at a flow rate of 4.5 mL/min while measuring Na, Mg, P, S, K, Ca, Cr, Mn, Fe, Ni, Cu, Zn, As, Se and Pb to exclude possible polyatomic interferences. The remaining elements were measured in the standard mode. Any isobaric interferences were automatically corrected by the instrument Syngistix™ software for ICP-MS v.2.3 (Perkin Elmer).

The ICP-MS measurements were performed in peak hopping scan mode with dwell times ranging from 25 to 50 ms depending on the element, 20 sweeps per reading and three replicates. The ICP-MS conditions were as follows: RF power – 1600 Watts, auxiliary gas flow rate 1.20 L/min. Torch alignment, nebulizer gas flow and quadrupole ion deflector (QID) voltages (in standard and KED mode) were optimized before analysis for highest intensities and lowest interferences (oxides and doubly charged ions levels lower than 2.5%) with NexION Setup Solution containing 1 µg/L of Be, Ce, Fe, ln, Li, Mg, Pb and U in 1% nitric acid using a standard built-in software procedure. To correct for variation between and within ICP-MS analysis runs, liquid reference material was prepared using pooled digested samples, and run after the instrument calibration, and then after every nine samples in all ICP-MS sample sets. Equipment calibration was performed at the beginning of each analytical run using seven multi-element calibration standards (containing 2 µg/L In internal standard) prepared by diluting 1000 mg/L single element standards solutions (Inorganic Ventures; Essex Scientific Laboratory Supplies Ltd) with 10% nitric acid. As a calibration blank, 10% nitric acid containing 2 µg/L In internal standard was used and it was run throughout the course of the analysis.

Sample concentrations were calculated using the external calibration method within the instrument software. Further data processing, including the calculation of final elements concentrations was performed in Microsoft Excel. First, sample sets run at different times were connected as an extension of the single-run drift correction. Linear interpolation between each pair of liquid reference material standards was used to generate a theoretical standard for each sample that was then used to correct the drift by simple proportion to the first liquid reference material standard analysed in the first run. Liquid reference material composed of pooled samples was used instead of the CRM to match the chemical matrix of the samples as closely as possible, thereby emulating the sample drift. Second, the blank concentrations were subtracted from the sample concentrations and each final element concentration was obtained by multiplying by the dilution factor and normalizing the element concentrations to the sample’s dry weight.

#### 1.4 Soil porosity analysis

To quantify soil porosity, we used X-ray computed tomography. First, six soil samples were collected in October 2020 from each of the selected locations in Kansas (see materials and methods 1.1). The top layer of soil was removed (10 cm), and the soil samples were collected using polyvinylchloride (PVC) columns of 5 cm internal diameter and 7 cm length. After the soil collection, the bottom of each column was sealed with tape to retain the soil in the column.

The undisturbed soil columns were non-destructively imaged using Phoenix v|tome|x MDT (Waygate Technologies GmbH, Wunstorf, Germany) at The Hounsfield Facility, University of Nottingham. Scans were acquired by collecting 2695 projection images at 180 kV X-ray energy, 200 µA current and 334 ms detector exposure time in fast mode (15 min total scan time per column). Scan resolution was 55 µm.

#### 1.5 Metagenomic analysis of free-living soil microbiota - DNA extraction

For metagenomic analysis, approximately 5 g of each soil sample was transferred into 50-mL conical tubes containing 20 mL of sterile distilled water. To remove large plant debris and soil particles, the samples were shaken thoroughly and filtered into new sterile 50-mL tubes using 100-μm nylon mesh cell strainers. The filtered soil solutions were centrifuged at high speed for 20 min in a centrifuge Eppendorf 5810R and most of the supernatants were discarded. The remaining 1-2 mL of supernatants were used to dissolve the soil pellets. The resulting suspensions were transferred to sterile 1.5-mL Eppendorf tubes. Samples were centrifuged again at high speed in a benchtop centrifuge and the supernatants were discarded. The remaining pellets were stored at - 80°C for DNA extraction. For DNA isolation we used 96-well-format MoBio PowerSoil Kit (MOBIO Laboratories; QIAGEN) following the manufacturer’s instructions. Before starting the extraction, all samples were manually randomized by placing them in a plastic bag that was shaken several times. Samples were then taken individually from the bag and loaded in the DNA extraction plates. This random distribution was maintained throughout library preparation and sequencing.

#### 1.6 Metagenomic library preparation and sequencing

DNA sequencing libraries were prepared using the Rapid PCR Barcoding Kit (SQK-RPB004) from Oxford Nanopore Technologies, UK. In brief, 1 µL Fragmentation Mix (FRM) was added to 3 µL DNA (2-10 ng/µL), and the reaction was mixed by gently finger-flicking. The DNA was fragmented using the following conditions: 30°C for 1 min, then 80°C for 1 min in an Applied Biosystems Veriti 96-Well Thermal Cycler (Applied Biosystems, CA, USA). The fragmented DNA was cooled, then barcoded and amplified in a PCR reaction containing 20 μL nuclease-free water, 25 μL LongAmp Taq 2X master mix (New England Biolabs, MA, USA), 4 μL fragmented DNA, and 1 µL barcode adaptor. The reaction was gently mixed and amplified using the following conditions: 95°C for 3 min, 20 cycles of denaturation at 95°C for 15 s, annealing at 56°C for 15 s and extension at 65°C for 6 min, and a final extension of 65°C for 6 min. The resulting DNA library was purified using 0.6X Agencourt AMPure XP beads (Beckman Coulter, CA, USA) and eluted in 10 μL 10 mM Tris-HCl pH 8.0, 50 mM NaCl. The library concentration was determined using a Qubit 4 Fluorometer with the Qubit dsDNA HS Assay Kit (Thermo Fisher Scientific, USA). Equimolar quantities of individual barcoded sample libraries were pooled and the volume was adjusted to 10 µL using 10 mM Tris-HCl pH 8.0, 50 mM NaCl. Subsequently, 1 µL rapid sequencing adapter (RAP) was added to the pooled library and the tube was incubated at room temperature for five minutes. Then, 34 μL Sequencing Buffer, 25.5 μL Loading Beads, and 4.5 μL nuclease-free water were added to the tube, and the contents were mixed gently. The prepared pooled library was added to a verified and primed FLO-MIN106 R9.4.1 flow cell (Oxford Nanopore Technologies, UK) in a MinION (Oxford Nanopore Technologies, UK) following the manufacturer’s instructions. DNA sequencing was conducted with default parameters using a MinIT (Oxford Nanopore Technologies, UK) with MinKNOW v2.1.12 (Oxford Nanopore Technologies, UK). Fast5 files were base-called with Guppy v4.0.15 using the template_r9.4.1_450bps_hac.jsn high accuracy model (Oxford Nanopore Technologies, UK).

#### 1.7 Metagenomic sequences processing

The initial dataset underwent demultiplexing, and primer and barcode sequences were trimmed using qcat v1.1.0 (Oxford Nanopore Technologies Ltd., UK). Reads with ambiguous barcode assignments were excluded from further analysis. The reads were filtered with NanoFilt v2.8.0^40^ to discard low-quality sequences (Q-score < 9) and sequences < 100 bp. We used the Kraken v2.1.2 pipeline^41^ to classify the whole metagenome shotgun sequencing reads. The reads were classified using the Kraken 2 archaea, bacteria, viral, plasmid, human, UniVec_Core, protozoa, and fungi reference database (k2_pluspf_20220607). To estimate relative abundances, the Bracken v2.7 pipeline^42^ was applied to the classification results. Subsequently, Pavian v1.0^43^ was used to extract abundance and taxonomic tables.

#### 1.8 Precipitation Gradient Data Analysis

##### 1.8.1 Precipitation data analysis

To determine the differences in precipitation levels across regions, we compared the mean annual precipitation in each region by fitting a linear model with the following formula:

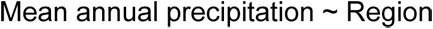

Differences between regions were indicated using the confidence letter display derived from Tukey’s *post hoc* test implemented in the package multcomp v.1.4.25^44^. We inspected the normality and variance homogeneity (here and elsewhere) using Q-Q plots and the Levene test, respectively. We visualized the results via a point range plot using the ggplot2 v3.5.1 R package^45^.

##### 1.8.2 Soil elemental content analysis

For the soil elemental profile, we created a matrix in which each cell contained the calculated element concentration in one sample. Then, we applied a z-score transformation to each ion across the samples in the matrix. Afterward, we applied a principal component analysis (PCA) using the Euclidean distance between samples and the z-score matrix as input to compare the elemental profiles of soils. Additionally, we estimated the variance explained by porosity, precipitation, region, and the interaction between them by performing a PERMANOVA via the function *adonis2* from the vegan v.2.6-4 R package^46^. The significant variables were visualized via a stacked bar plot using the *ggplot2* v3.5.1 R package.

To visualize the mineral content in each region, the z-score matrix created above was hierarchically clustered (method *ward.D*, function *hclust*), and we visualized the results using a heatmap. The rows in the heatmap were ordered according to the dendrogram order obtained from the clustered ions, and the regions were ordered according to the precipitation gradient (low precipitation to high). The heatmap was coloured based on the z-score.

To explore the relationship between ion concentrations and the precipitation gradient, we performed a correlation test using *cor* function from *stats* v.4.3.1 package in R^47^ of the average z-score measurement of each ion against the precipitation gradient. Afterward, we plotted the correlation coefficient for each ion in a barplot.

##### 1.8.3 Soil porosity analysis

The soil core images taken using x-ray computed tomography were analyzed using ImageJ v. 1.54b^48^. First, XY slice projection images were filtered using a median function to remove any noise from the raw data then an automatic threshold (Li method) was applied to produce binary images. In the binary images pores and solid particles were represented by black and white pixels, respectively. Afterwards, a region of interest (ROI) was defined in the central part of each projection to remove any potential border effect, cropping to a 600×600 pixel area. From the ROI defined in each image we extracted soil features, including particle area, perimeter, circularity, roundness, solidity, compactness, percentage of pores and pore size using the *measurement* function.

To remove the variability in the topsoil due to transportation and handling, we used two strategies. Firstly, we plotted the pore average size (mm) and soil porosity (%) for each sample. Then, we excluded from the analysis all projections having a value of soil porosity > 40% and considered the topsoil of those samples as the first projections after the exclusion. The second strategy was applied to samples with an irregular shape (e.g., mountain-like shape) at the topsoil level. We discarded all the projections with an irregular shape until we found projections with a regular distribution of soil layers. We created a data frame with the projection number (or slice) and the soil depth (projection number multiplied by the resolution). The soil porosity for each soil type was visualized via a point plot using ggplot2 v3.5.1 R package^45^.

To compare the soil elemental profiles against soil porosity, first, we applied a z-score transformation of each ion and soil porosity across the samples. Then, we estimated the distance between samples using Euclidean distance. Afterwards, we contrasted the dissimilarity matrices of each pair of datasets (soil elemental profile vs. soil porosity) using the *mantel* test implemented in the vegan v.2.6-4 R package^46^. Finally, we computed the significance of the correlation between matrices by permuting the matrices 10,000 times^44^.

##### 1.8.4 Taxonomic data analysis

To compare the alpha diversity across regions, we calculated the Shannon diversity index using the diversity function from the vegan v.2.6-4 package in R^46^. We used an ANOVA to test the alpha diversity differences between regions. Differences between regions were indicated using the confidence letter display derived from Tukey’s *post hoc* test implemented in the R package multcomp v.1.4.25^44^.

The beta diversity analysis (Principal Coordinates Analysis, PCo) was based on Bray-Curtis dissimilarity matrices calculated using the rarefied relative abundance tables. Additionally, we estimated the variance explained by soil porosity, precipitation, regions, and the interaction between them by performing a PERMANOVA via the function adonis2 from the vegan v.2.6-4 R package^46^. The significant variables were visualized via a stacked bar plot using ggplot2 v3.5.1 R package^45^.

The relative abundance of bacterial phyla was depicted using a stacked bar plot using ggplot2 v3.5.1 package.

To compare the microbiome composition against the elemental profiles and soil porosity, we contrasted the dissimilarity matrices of each pair of datasets (soil microbiome vs. soil elemental profile and soil microbiome vs. soil porosity) using the *mantel* test implemented in the vegan v.2.6-4 R package^46^. Briefly, we calculate the microbiome dissimilarity matrix using Bray-Curtis distance. Next, we applied a z-score transformation of each ion and soil porosity across the samples. Afterward, we calculated the soil elemental dissimilarity matrix and soil porosity dissimilarity matrix using Euclidean distance. Finally, we used the *mantel* test to compare and test the significance of the correlation between matrices by permuting the matrices 10,000 times^44^.

##### 1.8.5 Heatmap and enrichment analysis

We used the R package DESeq2 v.1.40.2^49^ to compute the bacterial enrichment profiles in the soils across the precipitation gradient. For each taxID (NCBI’s taxonomic identifier assigned to the taxa) in the rarefied table, as well as at the species, and family levels, we estimated difference in abundance compared to the wettest collection site (Welda Prairie) using a generalized linear model (GLM) with the following design:

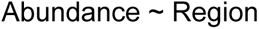

We extracted the following comparisons from the fitted model: CWR vs WEL, HAY vs WEL, KNZ vs WEL, SVR vs WEL, TRI vs WEL. A taxID, species, or family was considered statistically significant if it had a *p*-value < 0.05. We visualized the results using a heatmap. The rows in the heatmap were ordered according to the dendrogram obtained from the tax ID, species and family analysis. The relative abundance matrix was standardized across the significant tax ID, species, and family by using the z-score and the heatmap was coloured based on this value.

##### 1.8.6 Identification of marker taxa associated with precipitation gradients

To identify the corresponding bacterial isolates considered as “biomarker” taxa associated with precipitation gradients, we identified the principal components (PCs) that explain more than 80% of the variance in the data. These identified PCs were used to control the effects of soil elemental profile on the taxa abundances. Then, we fit five models: Poisson, negative binomial, two zero-inflated, and a multiple regression model.

For Poisson and negative binomial models, we fitted the following design:

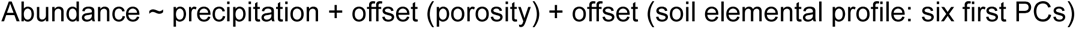

In parallel, we fitted the zero-inflated models using the following design:

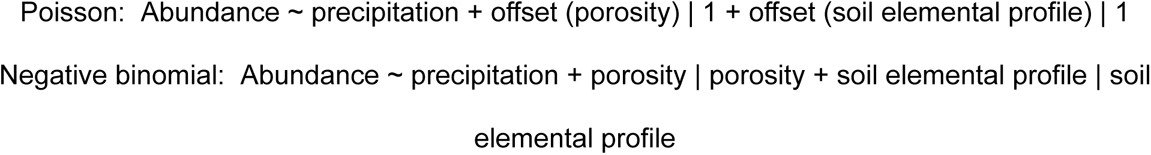

Next, to assess the statistical significance, we applied ANOVA to the best-performing model for each taxon according to the Akaike Information Criterion (AIC). Additionally, we applied a multiple regression model with the following design:

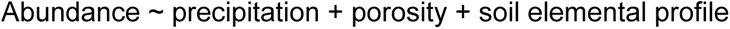

Then, we applied ANOVA to find taxIDs with a significant partial regression coefficient for precipitation.

A taxa ID was considered a “marker” if it had a relative abundance > 0.01 and a prevalence > 20%. We visualized the average standardized relative abundance (z-score) of the significant taxa ID in a point plot using ggplot2 v3.5.1 package^45^.

##### 1.8.7 Enrichment of bacteria biological functions associated with precipitation gradients

To identify biological processes enriched within the microbial communities, the sequence reads were assembled into contigs for each sample using metaFlye from the Flye v2.9 package^50^ with default mode. The contigs generated were then grouped and deduplicated using the dedupe.sh tool in BBTools v38.76^51^ to eliminate redundancies. Next, we determined the relative abundance of the contigs by mapping the reads from the samples to the contigs using minimap2 v2.17^52^, and extracting the relative abundance counts using CoverM v0.6.1^53^ in the ‘contig’ mode and reads_per_base coverage method. Taxonomic classification of the contigs was performed using the CAT v8.22 taxonomic classification pipeline^54^. Subsequently, the contigs were filtered to retain only bacteria sequences. DESeq2 v1.40.0^49^ was used to determine the contig enrichment profiles by fitting a GLM with the following design:

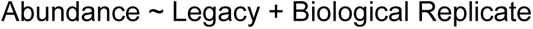

The low-precipitation soil versus the high-precipitation soil contrast was extracted from the fitted model. Contigs meeting the criteria of an FDR-adjusted *p*-value (*q*-value) <0.05 and a log_2_-transformed fold change >2 were selected for further analysis. Open reading frames encoded within the contigs were predicted using FragGeneScanRs v1.1.0^55^ with default settings. This was followed by functional annotation of the predicted proteins using the eggNOG-mapper v2.1.9^56^ pipeline with the eggNOG v5.0.2 database^57^ with Diamond v2.0.11^58^ and MMseqs2^59^.The genes annotated with Gene Ontology (GO) classifications were subsequently extracted, and a GO enrichment analysis focusing on biological processes was conducted. This involved employing adaptive GO clustering in conjunction with Mann–Whitney U testing, using the GO_MWU tool^60^, which evaluates the enrichment of each GO category based on whether genes linked to the GO category are significantly clustered at either the top or bottom of a globally ranked gene list. First, genes were ranked based on the signed log_2_-transformed fold change values. For each gene, any missing parental terms for specific GO categories were then automatically added. Next, fully redundant categories (those containing identical sets of genes), were collapsed into the more specific GO term. To further streamline the analysis, highly similar categories were grouped using complete linkage clustering based on the fraction of shared genes. We used default settings, where GO categories were merged if the most dissimilar pair within a group shared more than 75% of the genes in the smaller category. The merged group was named after the largest category. Significantly enriched and depleted GO categories were then determined by an adjusted *p*-value of <0.05. This approach simplified the GO hierarchy and addressed multiple testing which improved the statistical power of the GO enrichment analysis. The most prominent enriched and depleted GO categories shared across comparisons were visualized in ggplot2 v3.4.2 and coloured based on the square root transformed delta rank values (enrichment score) of the GO categories.

##### 1.8.8 Analysis of genetic variation among bacterial lineages along the precipitation gradient

To assess genetic differences between bacteria lineages along the precipitation gradient, we focused on 15 of the identified bacterial markers (Pseudomonas ID287, Salmonella ID28901, Sorangium ID56, Bradyrhizobium ID722472, Luteitalea ID1855912, Bradyrhizobium ID1355477, Flavisolibacter ID661481, Bradyrhizobium ID858422, Rubrobacter ID2653851, Bradyrhizobium ID1437360, Candidatus Koribacter ID658062, Streptomyces ID1916, Klebsiella ID573, Bradyrhizobium ID1325107, Edaphobacter ID2703788), as well as 18 additional abundant and prevalent species (Rubrobacter ID49319, Bacillus ID1428, Bradyrhizobium ID1274631, Priestia ID1404, Lacibacter ID2760713, Bradyrhizobium ID1325100, Candidatus Solibacter ID332163, Burkholderia ID28450, Flavisolibacter ID1492898, Bacillus ID1396, Escherichia ID562, Rhizobium ID384, Rubrobacter ID2653852, Microvirga ID2807101, Archangium ID83451, Pseudomonas ID303, Paenibacillus ID1464, Nitrosospira ID1231) across the samples, as proxies for the broader bacterial communities. These taxa were selected for their high genome coverage across samples, enabling more precise allele frequency estimates. Reference genomes for each species were retrieved from the NCBI Genome database, and the filtered shotgun metagenomic reads were aligned to these genomes using minimap2 v2.17-r941^52^. Alignments were sorted and indexed with SAMtools v1.18^61^, followed by variant calling using BCFtools v1.18^62^. This process identified 23,197,278 sequence variants, which were then filtered using VCFtools v0.1.16^63^, to retain only biallelic single nucleotide polymorphisms (SNPs) for further analysis. SNP filtering criteria included a variant quality score >20, a minor allele frequency (MAF) >0.01, <50% missing data, and a minimum sequencing depth of 10x in each sample. After filtering, 23,061 high-quality biallelic SNPs were retained. Genetic distances were computed using PLINK v1.90p^64^, and reduced to two dimensions through classical multidimensional scaling with the stats v4.3.0 package. PCoA plots were created with ggplot2 v3.4.2^45^, colored by soil type, precipitation levels and geographical region.

To identify genes potentially under selection across the precipitation gradient, we conducted a genetic-environment association (GEA) analysis. For this, SNPs were re-filtered using VCFtools v0.1.16^63^ with the same criteria, but allowing <50% missing data and a minimum sequencing depth of 5x per sample. After filtering, 93,013 biallelic SNPs were retained. Subsequently, GEA analysis was performed using a general linear model in the rMVP v1.1.1 package^65^, with native SNP data imputation, and average precipitation at each sampling location as the environmental variable. The genetic structure in the data was corrected using the first 10 principal components (PCs). Manhattan plots were generated using CMplot v4.5.1^66^ and significant associations were identified using the permutation method within the rMVP package.

### 2. Conditioning phase: Soil drought legacy is resilient to short-term perturbations

#### 2.1 Experimental design

The six soils collected from across the Kansas precipitation gradient, as described in materials and methods 1.1, were used in this experiment. The conditioning perturbations imposed in these experiments took place over approximately 20 weeks at the University of Kansas from December 17th, 2020 to May 5th, 2021. Each of the six input soils remained independent throughout the experiment. Mesocosms consisted of a 1:1 (v/v) mixture of field-collected soil to sterile turface MVP (Turface Athletics, Buffalo Grove, IL). A total of 192 sterile 100 mL pots were filled with the six soils and which were then randomly assigned to one of four conditions in a fully-factorial design: with or without a host, and either water-stressed or well-watered. Half the pots were planted with seedlings of the native prairie grass *Tripsacum dactyloides* (Eastern gamagrass, cultivar “Pete”); the rest remained unplanted. These 24 treatment groups (6 soils X 2 water-stressed/well-watered X 2 planted/unplanted) each had *N=*8 replicates for a total of 192 experimental soils in pots. All mesocosms were allowed to adapt to their watering regimes in a growth chamber set to a 12-hour day cycle, 27°C/23°C, and ambient humidity. Well-watered control pots were watered every 1-2 days and water-stressed plants were watered every 3-5 days when plants displayed drought symptoms (e.g., leaf curling). All pots were fertilized with 35 mL of 1mL/L concentration of Bonide 10-10-10 plant food (Bonide Products LLC, Oriskany, NY) on week 8 and week 12.

#### 2.2 Sample collection

To characterize the baseline microbial communities going into the conditioning treatments, we sampled four replicates of each soil/treatment combination one week after beginning the experiment. To collect the samples, the top centimeter of soil was discarded and the remaining soil was homogenized to ensure even sampling of the top, middle, and bottom of the pot. Two grams of this homogenized soil were placed in a 15mL tube, flash-frozen in liquid nitrogen, and stored at −80°C for microbial DNA and RNA extraction. To characterize the effects of the four treatments on the microbial communities, we sampled all remaining replicates at the end of the 20-week conditioning phase. Soil samples were collected as described for the baseline communities, but an additional 6 g of homogenized soil was preserved at 4°C in 50 mL conical tubes for use as inocula in a downstream experiment. Additionally, for the planted pots, we measured *T. dactyloides* shoot height before uprooting the plants. We collected samples of a crown root from each plant (3 cm long each, beginning 2 cm from the base of the plant) and stored them in 50% ethanol at 4°C for downstream laser ablation tomography analysis. The remaining roots and shoots were dried in an oven at 225°C for 12 hours and then weighed separately.

#### 2.3 Changes in bacterial community structure associated with drought and well-watered conditioning with and without a host

##### 2.3.1 DNA extraction

Total DNA was extracted from baseline and post-conditioning soil sub-samples using the DNA Set for NucleoBond RNA Soil Kit (Macherey-Nagel, Düren, Germany) according to the manufacturer’s instructions.

##### 2.3.2 Library preparation and sequencing

DNA sequencing libraries were prepared using the Rapid PCR Barcoding Kit (SQK-RPB004) from Oxford Nanopore Technologies, UK, and sequenced on a FLO-MIN106 R9.4.1 flow cell (Oxford Nanopore Technologies, UK) in a MinION (Oxford Nanopore Technologies, UK), with a MinIT (Oxford Nanopore Technologies, UK) using MinKNOW v2.1.12^67^ (Oxford Nanopore Technologies, UK) as described in materials and methods section 1.6.

##### 2.3.3 Sequence processing

Raw sequence data were demultiplexed and primer and barcode sequences were trimmed using qcat v1.1.0 (Oxford Nanopore Technologies Ltd., UK). Reads with ambiguous barcode assignments were excluded from further analysis. The reads were filtered with NanoFilt v2.8.0^40^ to discard low-quality sequences (Q-score <9) and sequences <100 bp. We used the Kraken v2.1.2 pipeline^41^ for classifying the whole metagenome shotgun sequencing reads. The reads were classified using the Kraken 2 archaea, bacteria, viral, plasmid, human, UniVec_Core, protozoa and fungi reference database (k2_pluspf_20220607). To estimate relative abundances, the Bracken v2.7 pipeline^42^ was applied to the classification results. Subsequently, Pavian v1.0^43^ facilitated the extraction of abundance and taxonomic tables. Functions in phyloseq v1.44.0^68^ with microbiome v1.22.0 and microbiomeutilities v1.0.17^69^ were used to filter the dataset and remove samples with low read depth (<1000 reads), remove unidentified taxa and singletons, transform abundance values using rarefaction, subset and merge sample and taxonomic groups and perform other data frame manipulations.

#### 2.4 Plant biomass

Root and shoots were detached, dried in an oven at 225°C for 12 hours, and then weighed.

#### 2.5 Root Laser Ablation Tomography (LAT) analysis

We collected samples of a crown root from each plant (3 cm long each, beginning 2 cm from the base of the plant) and stored them in 50% ethanol at 4°C. The samples were shipped to the University of Nottingham for downstream LAT analysis. Briefly, root segments were dehydrated in 100% methanol for 48 hours, transferred to 100% ethanol for 48 hours, then dried with an automated critical point dryer (CPD, Leica EM CPD 300, Leica Microsystem). Root anatomical images were acquired using a laser ablation tomograph (LATScan, Lasers for Innovative Solutions LLC). This utilises a combination of precise positioning stages with a guided pulsed UV (355 nm) laser, to thermally vaporise thin sections of the root, and then to illuminate the exposed surface. The tomograph was retrofitted with a microscopic imaging system, using a machine vision camera unit (Model Grasshopper3, FLIR) and infinity-corrected long working distance magnifying objectives (Mitutoyo (UK) Ltd.)

#### 2.6 Changes in gamagrass morphological features under drought and well-water conditioning

To identify morphological features in gamagrass that changed with the conditioning (drought and well-watered) treatments, we used Image J to quantify the average area aerenchyma, number aerenchyma, number metaxylem, cortical cell layers, total area metaxylem, average area metaxylem, stele min. diameter, stele max. diameter, stele area, stele perimeter, total area aerenchyma, adjusted cortex area, root min. diameter, cortex min. diameter, root max. diameter, cortex max. diameter, total perimeter, cortex perimeter, root total area, and cortex area. Additionally, we quantified the root-shoot ratio, number of leaves, number of green leaves, root mass, shoot mass, and shoot height.

#### 2.7 Metatranscriptome analysis

##### 2.7.1 RNA isolation

Total RNA was extracted from baseline and post-conditioning soil sub-samples with the NucleoBond RNA Soil Kit (Macherey-Nagel, Düren, Germany) using the manufacturer’s instructions. Isolated RNA was treated with Turbo DNA-free (Applied Biosystems, Waltham, MA, USA) to remove contaminating DNA, following the manufacturer’s instructions.

##### 2.7.2 Library preparation and sequencing

RNA libraries were prepared using 1 µg of total RNA according to established protocols with modifications. Briefly, poly(A)-tail-containing RNA was removed from the RNA samples using Sera-mag oligo(dT) magnetic beads (GE Healthcare Life Sciences, Marlborough, MA, USA) and then the samples were subjected to ribodepletion with the NEBNext rRNA Depletion Kit (New England Biolabs, MA, USA), following the manufacturers’ instructions. The purified RNA was resuspended in a fragmentation mix consisting of 6.25 µL Milli-Q water, 5 µL 5x First strand buffer, and 1.25 µL random primers (3 µg/µL) and fragmented at 94°C for 6 min. First-strand cDNA synthesis was performed using a mixture of 0.8 µL reverse transcriptase, 2 µL 100 mM DTT, 0.4 µL 25 mM dNTP, 0.5 µL RNAseOUT (40U/µL), 10 µL RNA, and 6.3 µL Milli-Q water. The reactions were incubated at 25°C for 10 min, 42°C for 50 min, and 70°C for 15 min. Second-strand cDNA synthesis was performed by adding a master mix of 18.4 µL Milli-Q water, 5 µL 10X second strand buffer, 1.2 µL 25 mM dNTP, 0.4 µL RNAse H (5U/µL), and 5 µL DNA Pol I (10U/µL) to the sample, followed by incubation at 16°C for 1 h. The samples were then purified using Agencourt AMPure XP beads. Subsequently, the libraries were end-repaired with a mixture of 30 μL sample, 2.5 μL of 3 U/μL T4 DNA polymerase, 0.5 μL of 5 U/μL Klenow DNA polymerase, 2.5 μL of 10 U/μL T4 PNK, 5 μL of 10X T4 DNA ligase buffer with 10 mM ATP, 0.8 μL of 25 mM dNTP mix, and 8.7 μL Milli-Q water, incubated at 20°C for 30 min, and purified again using Agencourt AMPure XP beads. Following this, the RNA libraries were adenylated in a mix containing 34 μL of the end-repaired sample, 3 μL of 5 U/μL Klenow exo-, 5 μL of 10X Enzymatics Blue Buffer, 1 μL of 10 mM dATP, and 9 μL of Milli-Q water. The mixture was incubated at 37°C for 30 min, followed by 70°C for 5 min, and then purified using Agencourt AMPure XP beads. Individual samples were indexed through ligation using a mix of 10.25 µL sample, 1 μL of 600 U/μL T4 DNA ligase, 12.5 μL of 2x Rapid Ligation Buffer, and 1.25 µL of 2.5 µM indexing adapter from the KAPA Dual-Indexed Adapter Kit (Kapa Biosystems, MA, USA). The samples were incubated at 25°C for 15 min, followed by the addition of 5 μL of 0.5 M EDTA pH 8. The libraries were purified twice with Agencourt AMPure XP beads. The libraries were enriched in a reaction containing 20 μL sample, 25 μL of 2X KAPA HiFi HS Mix (Kapa Biosystems, MA, USA), 2.5 μL of 5 μM I5 primer, and 2.5 μL of 5 μM I7 primer. The reactions were initially heated to 98°C for 45 seconds, followed by 14 cycles of 98°C for 15 seconds, 60°C for 30 seconds, and 72°C for 30 seconds, with a final extension at 72°C for 1 minute. The resulting RNA libraries were purified using Agencourt AMPure XP beads, quantified on a Qubit 4 Fluorometer (Thermo Fisher Scientific, USA), and the library size was assessed using High Sensitivity D1000 ScreenTape on the Agilent 4200 TapeStation (Agilent Technologies, Santa Clara, CA). Equimolar quantities of individual barcoded RNA libraries were pooled in a randomized manner and shipped on dry ice to Beijing Genomics Institute (BGI, Shenzhen, China). Each library pool was sequenced on an MGI Tech MGISEQ-2000 sequencing platform to generate a minimum of 10 million 100 bp paired-end reads per sample.

2.7.3 *Taxonomic classification of transcripts*

Cutadapt v4.6^70^ was used to remove primer and barcode sequences and low-quality sequences from the paired-end reads of the sequenced RNA libraries. To identify taxa with enriched gene expression activity, the reads were classified using the Kraken v2.1.2 pipeline^41^ with the archaea, bacteria, viral, plasmid, human, UniVec_Core, protozoa and fungi reference database (k2_pluspf_20220607), and the Bracken v2.7 pipeline^42^ was applied to the classification results to estimate the relative abundances. The counts table was generated from Pavian v1.0^43^. Data filtering and statistical analysis were then performed as before using phyloseq v1.44.0^68^ with microbiome v1.22.0^71^ and microbiomeutilities v1.0.17^69^.

#### 2.8 Data Analysis

##### 2.8.1 Changes in bacterial community structure associated with drought and well-watered conditioning with and without a host

To assess the alpha diversity across the samples, we calculated the Shannon Diversity Index using phyloseq v1.44.0^68^. We used ANOVA to test for significant differences in Shannon Diversity indices between groups and means were separated using Tukey’s honestly significant difference (HSD) test from the agricolae v1.3.5 R package^72^. For beta diversity, Bray-Curtis dissimilarity matrices were calculated using phyloseq v1.44.0 and the variance explained by legacy, conditioning, and host were estimated by performing permutational multivariate analysis of variance (PERMANOVA) using the adonis2 function in vegan v2.6.4 R package^46^.

Constrained ordination of beta-diversity was plotted using canonical analysis of principal coordinates (CAP) based on Bray-Curtis dissimilarity matrices calculated with vegan v2.6.4. We visualized differences with the CAP analysis, using the following models:

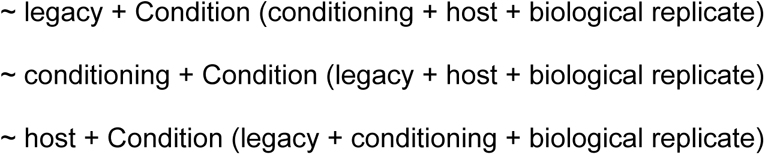

The relative abundance of taxa was plotted as a stacked bar representation using phyloseq v1.44.0. The tax_glom function in phyloseq v1.44.0 was used to agglomerate taxa, and the aggregate_rare function in microbiome v1.22.0 was used to aggregate rare groups. We used DESeq2 v1.40.0^49^ to calculate the enrichment profiles by fitting a generalized linear model (GLM) with the following design:

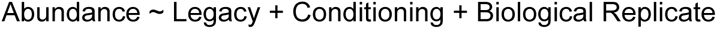

We extracted the following comparisons from the fitted model: wet soil legacy with watered conditioning vs wet soil legacy (baseline), wet soil legacy with drought conditioning vs wet soil legacy (baseline), dry soil legacy (baseline) vs wet soil legacy (baseline), dry soil legacy with watered conditioning vs wet soil legacy (baseline), and dry soil legacy with drought conditioning vs wet soil legacy (baseline). Taxa were considered significant if they had an FDR-adjusted *p*-value (*q*-value) <0.05. The results of the GLM analysis were rendered in heatmaps, coloured based on the log_2_-transformed fold change output by the GLM. Significant differences between comparisons with a *q*-value <0.05 with log_2_-transformed fold change >2 were highlighted with black squares.

Relative abundances of the taxonomic markers were extracted, and an ANOVA was performed to assess significant differences between treatment groups. Tukey’s Honest Significant Difference (HSD) test, implemented using the agricolae v1.3.5 R package^72^, was used for post-hoc pairwise comparisons. To further explore the effects of watering and drought treatments, with and without a host, on the relative abundances of the taxonomic markers, we subset the data and applied a generalized linear model (GLM) using DESeq2 v1.40.0^49^. The model was structured as follows:

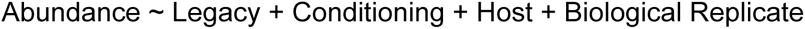

We then extracted the following comparisons from the fitted model for each soil legacy: water conditioning vs baseline, drought conditioning vs baseline, water conditioning with host vs baseline, drought conditioning with host vs baseline. Markers were considered significant if the FDR-adjusted p-value (q-value) was < 0.05. Results from the GLM analysis were visualized in a heatmap, with colours representing log2-transformed fold changes. Comparisons showing significant differences (q-value <0.05 and log2-transformed fold change >2) were highlighted with black squares.

##### 2.8.2 Gene Ontology (GO) term enrichment analysis

To identify enriched biological processes within the microbial communities, sequence reads from individual samples were assembled into contigs using metaFlye from the Flye v2.9 package^50^ with default parameters, as described in Methods Section 1.8.7. Relative abundance counts were then determined, and the resulting contigs were subjected to taxonomic classification and filtering, also as outlined in Methods Section 1.8.7. We used DESeq2 v1.40.0^49^ to determine the bacterial contig enrichment profiles by fitting a generalized linear model (GLM) with the following design:

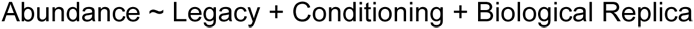

We extracted the following comparisons from the fitted model: wet soil legacy with watered conditioning vs wet soil legacy (baseline), wet soil legacy with drought conditioning vs wet soil legacy (baseline), dry soil legacy (baseline) vs wet soil legacy (baseline), dry soil legacy with watered conditioning vs wet soil legacy (baseline), and dry soil legacy with drought conditioning vs wet soil legacy (baseline). Contigs meeting the criteria of an FDR-adjusted *p*-value (*q*-value) < 0.05 and a log_2_-transformed fold change > 2 were selected. Open reading frames were predicted and functionally annotated, and genes with GO classifications were subjected to GO enrichment analysis with the GO_MWU tool^60^.

##### 2.8.3 Changes in gamagrass morphological features under drought and well-water condition

For each root feature identified, we used ANOVA to test for significant differences between groups and means were separated using Tukey’s honestly significant difference (HSD) test from the agricolae v1.3.5 R package^72^ Subsequently, the feature values were normalized using the rescale function from the scales v1.2.1 R package. The mean normalized feature values were then visually represented on a heatmap using ggplot2 v3.4.2. Subsequently, Pearson correlation coefficients between these features and corresponding *p*-values were computed using the rcorr function from the Hmisc v5.0.1 package^73^. The results of the correlation analysis were graphically presented using ggplot2 v3.4.2, where the colour of the plots reflected the correlation coefficient values. Significant correlations (*p* <0.05) were emphasized with black squares on the plots. Furthermore, the coefficient of variation for the feature values was calculated and depicted using ggplot2 v3.4.2. The three plots were integrated based on the hierarchical clustering of the Pearson correlation coefficients of the features. The clustering employed the ward.D2 method within the hclust function in R, utilizing Euclidean distances calculated using the dist function.

##### 2.8.4 Metatranscriptome sequence analysis

To assess transcriptional differences in the activity of the bacterial community, Bray-Curtis dissimilarity matrices were calculated. The variance explained by legacy, conditioning, and host was estimated using permutational multivariate analysis of variance (PERMANOVA) with the adonis2 function from the vegan v2.6.4 R package^46^. Beta-diversity patterns were visualized through constrained ordination using canonical analysis of principal coordinates (CAP). We applied CAP analysis to visualize differences using the following models:

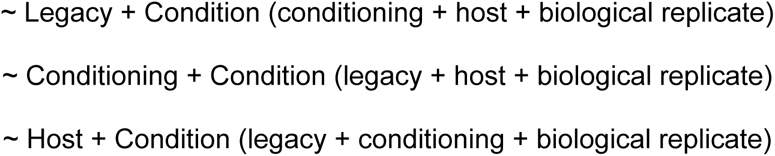

Transcriptional activity among taxa was displayed as a stacked bar plot using phyloseq v1.44.0. We employed DESeq2 v1.40.0^49^ to calculate enrichment profiles by fitting a generalized linear model (GLM) with the following design:

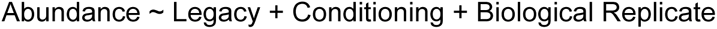

From the fitted model, we extracted the following comparisons: wet soil legacy with watered conditioning vs wet soil legacy (baseline), wet soil legacy with drought conditioning vs wet soil legacy (baseline), dry soil legacy (baseline) vs wet soil legacy (baseline), dry soil legacy with watered conditioning vs wet soil legacy (baseline), and dry soil legacy with drought conditioning vs wet soil legacy (baseline). Taxa were considered differentially abundant if the FDR-adjusted *p*-value (*q*-value) was <0.05. Results from the GLM analysis were visualized in heatmaps, where colours represent log2-transformed fold changes. Comparisons showing significant differences (*q*-value <0.05 and log2-transformed fold change >2) were highlighted with black squares.

Additionally, high-quality filtered reads of the transcriptome were de novo assembled into a reference metatranscriptome using Trinity v2.15.1^74^ with default parameters. Open reading frames in transcripts were predicted with TransDecoder v5.7.1^75^ with default settings. Functional annotation of the predicted proteins was performed using the eggNOG-mapper v2.1.9^56^ pipeline, utilising the eggNOG v5.0.2 database^57^ with Diamond v2.0.11^58^ and MMseqs2^59^. The taxonomic classification of transcripts was conducted using the CAT v8.22 taxonomic classification pipeline^54^. Sequence reads were further filtered using SortMeRNA v4.3.6^76^ with the smr_v4.3_default_db.fasta database to remove residual amplified rRNA sequences. Transcript quantification analysis was performed using Salmon v1.10.0 in the mapping-based mode with the de novo assembled reference metatranscriptome. Subsequently, the transcript-level abundance estimates from salmon were extracted for the identified transcripts using the R package tximport v1.28.044 as raw counts in default setting^77^. DESeq2 v1.40.0^49^ was utilized to determine the bacterial transcript enrichment profiles by fitting a generalised linear model (GLM) as described before. Genes meeting the criteria of an FDR-adjusted *p*-value (*q*-value) <0.05, a log_2_-transformed fold change >2, and had GO classifications, were subjected to GO enrichment analysis with the GO_MWU tool^60^.

##### 2.8.5 Analysis of genetic variation among bacterial lineages

Filtered shotgun metagenomic reads were aligned to the reference genomes of 33 selected taxa, including 22 identified bacterial markers and 11 additional abundant and prevalent species (see Supplementary Materials and Methods 1.8.8), using Minimap2 v2.17-r941^52^. The resulting alignments were sorted and indexed with SAMtools v1.18^61^. Variant calling was performed using BCFtools v1.18^62^, and variants were filtered with VCFtools v0.1.16^63^. Filtering criteria included a variant quality score >20, a minor allele frequency (MAF) >0.01, <50% missing data, and a minimum sequencing depth of 10x in each sample. After filtering, a total of 8,293 high-quality biallelic SNPs were available for further analysis. To assess genetic variation within and between groups, we analysed molecular variance (AMOVA) using poppr v2.9.6^78^. The significance of the AMOVA results was determined with a permutation test using the randtest function in ade4 v1.7.22^79^. To determine the extent of genetic differentiation between groups, we calculated the Fixation index (FST) values using the hierfstat v0.5.11^80^ package. Principal coordinates analysis (PCoA) plots were generated as previously described in Supplementary Materials and Methods 1.8.8, and coloured to reflect soil legacy, drought/ well-watered and host/ no-host treatments. To identify genes associated with soil legacy, we conducted a genome-wide association study (GWAS). SNPs were re-filtered using VCFtools v0.1.16^63^, applying the same criteria but allowing for up to 70% missing data and a minimum sequencing depth of 3x in each sample. GWAS was conducted using a general linear model in the rMVP v1.1.1 package^65^. Associations were identified by comparing bacterial lineages from dry legacy soil to those from wet legacy soil. Genetic structure was accounted for by incorporating the first 10 principal components. Significant associations were identified through permutation testing within the rMVP package, and Manhattan plots were generated using CMplot v4.5.1^66^.

### 3. Test phase: Effects of soil microbiome legacy on plant tolerance to drought

#### 3.1 Experimental design and non-destructive phenotypic measurements

At the end of the “Conditioning Phase”, homogenized soil was collected from each pot by discarding the top one centimeter of soil, mixing the soil in the pot with a clean plastic spatula, and placing six grams in a sterile 50 mL conical tube. For rhizosphere samples (i.e., planted pots), plants were gently pulled from the pots and the soil particles adhered to and within the root bundle were shaken into a sterile 50 mL conical tube. The rhizosphere particles were homogenized with a clean plastic spatula and particles were poured out of the tube until six grams remained. Soil and rhizosphere samples were stored at 4°C overnight.

Soil microbes were extracted from the 6 g soil or rhizosphere sample the following day by adding 25 mL of autoclaved 1X PBS with 0.0001% Tween-89 to the 50 mL tube containing the sample. Tubes were vigorously shaken to mix and break up large soil aggregates. Large particles were allowed to settle to the bottom of the tube; the supernatant was then filtered through autoclaved Miracloth (Sigma-Aldrich) into a new sterile 50 mL conical tube. Filtered samples were then centrifuged at 3600g for 25 min at 4°C. The supernatant was discarded, and the microbial pellet was resuspended in 6 mL of 1X PBS buffer using a vortex. The resuspended pellets were stored at 4°C until used for inoculations a few hours later.

As stated in section 2.1, the conditioning phase had 24 treatment groups with eight replicates of each treatment (192 pots total). For the test phase, microbial extracts from all eight replicates of each group (plus sterile buffer-only control inoculums) were each inoculated into a pot planted with gamagrass (N=200) and a pot planted with maize (N=200) that were then maintained under watered-stress (drought) conditions. Furthermore, four of the eight replicate extracted microbial inoculants (as well as four sterile buffer-only control inoculums) were each inoculated into an additional gamagrass planted (N=100) and maize planted (N=100) pots, which were then maintained under well watered control conditions. This makes for a total of N=600 plants at the start of the test phase. Throughout the experiment, nine water-stressed maize and five well-watered maize were lost (no gamagrass died). Therefore, phenotype measurements were completed on a total of N=586 plants. We chose this design because resource and space limitations prevented us from testing all 192 inocula under both drought and control conditions, and we were primarily interested in microbial effects on plant function under drought; we therefore opted to maximize our power to test for differences among the inocula under water limitations.

To create the inoculum, the resuspended pellet was inverted three times to mix and 1 mL of the sample was added to 100 mL of sterile 0.5X MS liquid medium, for a microbial titer equivalent to 0.01 g soil per mL. The “mock inoculation” controls were created by substituting 1 mL of sterile PBS for the resuspended microbial pellet. Finally, 25 mL of this suspension was inoculated onto the soil surface of each Test Phase pot. Thus, each microbial community extracted from one of the 192 conditioning phase pots was used to inoculate either two or four plants in the Test Phase. To maintain statistical independence of the experimental replicates from the conditioning phase, no pooling was performed.

Before inoculation, pots were planted with 3-4-day old gamagrass or maize germinants. Gamagrass seeds were soaked in 3% hydrogen peroxide for 24 hours and germinated in seed trays filled with sterile clay. Maize seeds were soaked in 70% ethanol for 3 minutes, then soaked in 5% NaClO on a rotator for 2 min, and then rinsed with sterile DI water three times. Treated maize seeds were germinated on sterile damp paper towels inside sealed plastic bags.

Pots were fully randomized and, prior to inoculation, were filled with a homogenized 5:1 (w/w) mixture of all-purpose sand (TechMix All-purpose 110241) and calcined clay (Pro’s choice rapid dry) that had been sterilized by autoclaving on a one-hour liquid cycle. Pots were autoclaved on a 30 min liquid cycle and then filled with the sand:clay mixture, leaving one inch of room at the top of the pot. To help keep the mixture from falling out of the drain holes in the bottom of the pots, a sterile filter paper was shaped into a cone, pushed to the bottom of the pot, and a sterile marble was used to weigh the paper down. This effectively blocked the substrate, but still allowed water to exit the drainage holes. Plants were grown under 12-h days, 27°C/23°C (day/night), and ambient humidity, with the light setting set to 1, which is equivalent to 312 μmol/m^2^s. Three-day old gamagrass and maize leaf photosynthetic rates and gas exchange were measured using the LI-6800 (LI-COR, Lincoln, NE, USA), across 3 days for each host.

The LI-6800 aperture was 2 cm, warmup tests were performed at the start of each measurement session, the Fluorometry was set to “on”, the APD_Leaf to set to 1.5 kpa. The newest fully emerged leaf, which was most commonly the 4th leaf, on each plant was clamped in the chamber and allowed to stabilize until all measurements were stable for at least 30 secs before the measurements were recorded, which took approximately five minutes per leaf. Similarly, the leaf chlorophyll content was also measured using the MC-100 Chlorophyll Concentration Meter (apogee instruments, Logan, UT, USA).

Maize plants were sampled four weeks after planting and gamagrass was sampled five weeks after planting. In total, we measured 300 T. dactyloides plants (200 droughted, 100 well-watered) and 286 maize plants (191 droughted, 95 well-watered).

At the end of the Test Phase, uprooted plants were gently shaken to remove the soil attached to roots, prior to the collection of phenotypic, transcriptomic, and microbiome data as described below. One crown root was cut off with a ceramic blade and placed in a 1.7 mL tube on dry ice for downstream DNA extraction for 16S rRNA gene sequencing. Another crown root (0.15 - 0.2 g) was cut off with a ceramic blade, placed in a 1.7 mL tube, and flash frozen in liquid nitrogen for downstream RNA extraction. In between plants, the ceramic blade, plastic tweezers, plastic cutting board, and gloves were cleaned with 30% bleach. All samples were held on dry ice and then transferred to a −80°C freezer for storage. Next, the root and shoot were separated with a ceramic blade. Three cm of crown root beginning ∼2 cm from the base of the shoot was cut with a ceramic blade and submerged in 50% EtOH for LAT analysis. The rest of the root system was submerged in 70% EtOH in a 50 mL centrifuge tube for downstream root architecture scanning. Shoot height and number of leaves were recorded. Shoots were placed in individual paper bags, dried in an oven at 225°C for 12 hours, and the shoot dry weight was recorded.

#### 3.2 Root system architecture and root biomass analyses

A Perfection V600 flatbed scanner (Epson, Nagano, Japan) was used to scan intact maize and gamagrass root systems collected from Test Phase plants. The scanner was set to professional mode, reflective, document, black and white mode, 600 dpi, with a threshold of 55. A clear plastic tray filled with clean water was placed on the scanning bed. Each root system was placed in the tray with water and the tangled roots were gently pulled apart using plastic tweezers until they were no longer overlapping. A small amount of fine fibrous roots that fell off during this process were pushed to the corner of the tray and not included in the root cluster scan. The scanned root images were then analyzed using Rhizovision Explorer software v. 2.0.3^81^ in “whole root” mode and converted to 600 dpi. The region of interest tool was used to outline the main root bundle before pressing play to collect feature measurements. Finally, after collection of root system architecture data, the roots were dried in an oven at 225°C for 12 hours and then weighed.

#### 3.3 Root Laser Ablation Tomography (LAT) analysis

We collected one crown root from each plant (3 cm long each, beginning 2 cm from the base of the plant) and stored them in 50% ethanol at 4°C. These samples were shipped to the University of Nottingham for LAT analysis. Briefly, root segments were dried with an automated critical point dryer (CPD, Leica EM CPD 300, Leica Microsystem). Then, samples were ablated by a laser beam (Avia 7000, 355 nm pulsed laser) to vaporize the root tissue at the camera focal plane ahead of an imaging stage and cross-sectional images were taken using a Canon T3i camera with a 5×micro-lens (MP-E 65 mm) on the laser-illuminated surface. ImageJ software^48^ was used to measure root anatomical traits captured in the high-quality LAT images.

#### 3.4 Leaf ionome and shoot biomass analyses

The elemental profiles of the shoots were measured using Inductively Coupled Plasma Mass Spectrometry (ICP-MS). The shoot biomass from all uprooted plants was dried in an oven at 225°C for 12 h and then weighed. The dried biomass samples were cut into small pieces using a clean ceramic scalpel and placed in 5 mL Eppendorf tubes with 3 zirconium oxide beads. Shoots were pulverized using a Tissue Lyzer II (Qiagen) using 2 cycles of 60 seconds at the frequency of 30 s^-1^. Next, 5-10 mg of pulverized shoot samples were weighted on a Mettler five-decimal analytical scale, and 1-3 mL (depending on the sample dry weight) of concentrated trace metal grade nitric acid Primar Plus (Fisher Chemicals) was added to each tube. Prior to the digestion, 20 µg/L of Indium (In) was added to the nitric acid as an internal standard to assess putative errors in dilution or variations in sample introduction and plasma stability in the ICP-MS instrument. The samples were then digested in DigiPREP MS dry block heaters (SCP Science; QMX Laboratories) for 4 h at 115°C. After cooling down, the digested samples were diluted to 10-30 mL (depending on the volume of the nitric acid added) with 18.2 MΩcm Milli-Q Direct water. The elemental analysis was performed using an ICP-MS, PerkinElmer NexION 2000 equipped with Elemental Scientific Inc 4DXX FAST Dual Rinse autosampler, FAST valve and peristaltic pump. The instrument was fitted with a PFA-ST3 MicroFlow nebulizer, baffled cyclonic C3 high sensitivity glass spray chamber cooled to 2 °C with PC3X Peltier heated/cooled inlet system, 2.0 mm i.d. quartz injector torch and a set of nickel cones. Twenty-four elements were monitored including the following stable isotopes: ^7^Li, ^11^B, ^23^Na, ^24^Mg, ^31^P, ^34^S, ^39^K, ^43^Ca, ^48^Ti, ^52^Cr, ^55^Mn, ^56^Fe, ^59^Co, ^60^Ni, ^63^Cu, ^66^Zn, ^75^As, ^82^Se, ^85^Rb, ^88^Sr, ^98^Mo, ^111^Cd, ^208^Pb and ^115^In. Helium was used as a collision gas in Kinetic Energy Discrimination mode (KED) at a flow rate of 4.5 mL/min while measuring Na, Mg, P, S, K, Ca, Ti, Cr, Mn, Fe, Ni, Cu, Zn, As, Se and Pb to exclude possible polyatomic interferences.

The remaining elements were measured in the standard mode. The instrument Syngistix™ software for ICP-MS v.2.3 (Perkin Elmer) automatically corrected any isobaric interferences. The ICP-MS measurements were performed in peak hopping scan mode with dwell times ranging from 25 to 50 ms depending on the element, 20 sweeps per reading and three replicates. The ICP-MS conditions were adjusted to an RF power of 1600 Watts and an auxiliary gas flow rate of 1.20 L/min. Torch alignment, nebuliser gas flow and quadrupole ion deflector (QID) voltages (in standard and KED mode) were optimized before analysis for highest intensities and lowest interferences (oxides and doubly charged ions levels lower than 2.5%) with NexION Setup Solution containing 1 µg/L of Be, Ce, Fe, ln, Li, Mg, Pb and U in 1% nitric acid using a standard built-in software procedure. To correct for variation between and within ICP-MS analysis runs, liquid reference material was prepared using pooled digested samples and run after the instrument calibration and then after every nine samples in all ICP-MS sample sets. Equipment calibration was performed at the beginning of each analytical run using seven multi-element calibration standards (containing 2 µg/L In internal standard) prepared by diluting 1000 mg/L single-element standards solutions (Inorganic Ventures; Essex Scientific Laboratory Supplies Ltd) with 10% nitric acid. As a calibration blank, 10% nitric acid containing 2 µg/L In internal standard was used, and it was run throughout the analysis. Sample concentrations were calculated using the external calibration method within the instrument software. Further data processing, including the calculation of final element concentrations, was performed in Microsoft Excel.

#### 3.5 Crown root transcriptomics

##### 3.5.1 RNA extraction and sequencing

For RNA extractions and sequencing, flash-frozen crown roots were freeze-dried for 48 hours and finely ground with pellet pestles. The RNA extraction protocol was carried out according to the NucleoSpin RNA Plant kit (Macherey-Nagel, Düren, Germany).

For the 132 maize samples, remnant DNA was removed from purified RNA using the DNA-Free kit (Invitrogen, Carlsbad, CA, USA). RNA-seq libraries were prepared using the QuantSeq 3’ mRNA-Seq V2 kit with unique dual sequences and the unique molecular identifier (UMI) module (Lexogen, Vienna, Austria) following the manufacturer’s recommendations. Libraries were pooled at equimolar concentrations and then sequenced (2×150bp, but reverse reads were not used) on a NovaSeq S4 flow cell (Illumina, San Diego, CA, USA) along with a 25% PhiX spike-in. Maize RNA-seq library preparations and sequencing were performed by the RTSF Genomics Core at Michigan State University. For the 132 *T. dactyloides* samples, RNA-seq libraries were prepared using the NEBNext Ultra II Directional Library Kit with the oligo-dT magnetic isolation module (New England Biolabs, Ipswich, MA, USA) and sequenced on the Illumina NovaSeq 6000 platform at the Genomic Sciences Laboratory at North Carolina State University to generate a minimum of 40M read pairs (2×150bp) per sample.

3.5.2 Sequence processing

For the maize sequence reads, UMIs were removed from all sequences and added to the read headers using UMI-tools^82^. Next, cutadapt version 4.2^70^ was used to remove the first four bases of each read, remove poly-A tails (if present), remove spurious poly-G runs using the --nextseq-trim=10 parameter, and remove adapter sequences. Reads that were <10 bp long or that aligned to maize rRNA gene sequences were removed; the remaining reads were aligned to the maize reference genome B73 RefGen_v5^83^ using HISAT2 version 2.2.1^84^. Aligned reads were converted to BAM format, sorted, and indexed using samtools version 1.9^61^. We then used UMI-tools^82^ to de-duplicate reads that both shared a UMI and had identical mapping coordinates. Finally, we used the FeatureCounts function of the subread package version 2.0.5^85^ with the maize genome annotation version Zm00001eb.1 and parameters -O --fraction -M --primary -g ID -t gene to generate a table of transcript counts.

For the *T. dactyloides* sequence reads, we first used cutadapt to remove NEBNext adapter sequences, poly-A tails, spurious poly-G runs, and low-quality tails using the -q 20,20 parameter and other default parameters. The cleaned reads were aligned to the *T. dactyloides* reference genome (Td-KS_B6_1-REFERENCE-PanAnd-2.0a^61^ using HISAT2. The alignments were name-sorted so that mate-pairs could be fixed using the fixmate function of samtools^61^, and then re-sorted based on coordinates, de-duplicated, and converted to indexed BAM files using the same software. Finally, a table of transcript expression estimates was generated using the FeatureCounts function of the subread package with parameters -p -O --fraction -M --primary -g ID -t gene and the *T. dactyloides* genome annotation version Td-KS_B6_1-REFERENCE-PanAnd-2.0a_Td00002ba.2.

##### 3.5.3 Statistical analyses

Because the maize and *T. dactyloides* RNA-seq datasets were generated using different approaches (3’ tag sequencing *vs.* full-length sequencing, respectively) we analyzed them in parallel rather than comparing them directly. For each species, we used DESeq2^49^ to identify genes that were differentially expressed between plants inoculated with microbiomes from a low-precipitation climate (“dry legacy”) *vs.* those inoculated with microbiomes from a high-precipitation climate (“wet legacy”). A single negative binomial model with default parameters was used to estimate log_2_-fold changes in gene expression due to inoculum legacy, while also controlling for the other experimental factors, using the model:

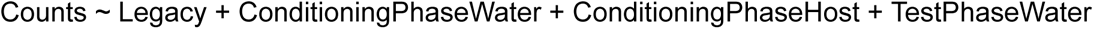

Statistical support was obtained using the Wald test with Benjamini-Hochberg FDR correction. All available samples were included in each analysis; thus, these results should be interpreted as the gene expression response to microbiome precipitation legacy, averaged across all levels of the other experimental factors.

In addition, we investigated whether plants’ gene expression responses to limited *vs.* ample water during the Test Phase were affected by inoculum precipitation legacy. To do so, we used DESeq2 to fit a model with the formula:

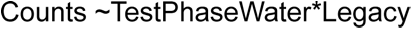

Then, we extracted the estimated log_2_-fold changes due to acute drought (relative to well-watered conditions) for both dry-legacy-inoculated and wet-legacy-inoculated plants. We inferred a meaningful interaction between these variables when the 95% confidence intervals of the two drought-induced log_2_-fold changes did not overlap at all. These interacting genes are candidates for linking real-time plant drought response to the microbiome’s historical environmental conditions.

Finally, we conducted a mediation analysis to determine whether the *T. dactyloides* genes that were sensitive to inoculum legacy were implicated in phenotypic responses to subsequent acute drought. A gene was considered legacy-sensitive if its expression was significantly affected by the main effect of inoculum legacy, or if it was affected by the interaction between inoculum legacy and test phase drought treatment, as described above. We summarized expression patterns of this subset of genes (normalized as transcripts per million, calculated using the full set of expressed genes) using non-metric multidimensional scaling of the Bray-Curtis distances among all *T. dactyloides* plants, which resulted in two axes of variation: MDS1 and MDS2. Next, we used the mediation package in R^86^ to compare the direct effects of test phase drought treatment on each focal plant trait (see section *3.7.1. Plant trait feature selection*) to the indirect effects of the drought treatment mediated through MDS1 and MDS2. Each mediation analysis used the linear models:

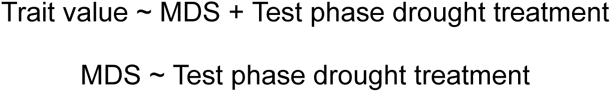

where MDS represents the “site score” of each individual plant on either MDS1 or MDS2. Separate models were fit to test the potential roles of MDS1 and MDS2 as mediator variables; however, a follow-up analysis using both MDS1 and MDS2 as simultaneous mediators, implemented in lavaan^87^, yielded equivalent conclusions.

3.5.4 Gene annotation

We downloaded maize and eastern gamagrass genome assemblies and annotations from MaizeGDB^83,88^. Functional information for maize genes was taken from the *Zea mays* genome annotation version Zm-B73-REFERENCE-NAM-5.0_Zm00001eb.1 and accessed using MaizeGDB’s MaizeMine tool^83,88–90^. For *Tripsacum dactyloides* genes, we relied on DNA sequence homology with annotated maize genes to infer function. We used BEDtools^91^ to extract gene coordinates and protein-coding sequences from the *T. dactyloides* reference genome Td-KS_B6_1-REFERENCE-PanAnd-2.0a^92^, and then used OrthoFinder^93^ to compare coding sequences from the two species. OrthoFinder identified 32,785 *T. dactyloides* genes (71.5% of the total) as homologs of maize genes, grouping them into 21,658 distinct orthogroups. All differential gene expression analyses, however, considered the entire set of expressed genes; those without maize orthologs, or with unannotated maize orthologs, were considered to be of unknown function.

#### 3.6 16S rRNA amplicon sequencing

##### 3.6.1 DNA extractions and library preparation

One crown root from each plant was cut off with a ceramic blade and placed in a 1.7 mL tube and flash frozen in liquid nitrogen for downstream amplicon library preparation. After collection, root sub-samples were kept on dry ice and ceramic tweezers were used to transfer the whole root to 1.1 mL cluster tubes (USA Scientific, Ocala, FL, USA). Tweezers were sterilized with 80% EtOH between samples. Roots were selected at random for placement in cluster tubes and were stored at −80°C until DNA extraction, at which time roots were freeze-dried for 48 hours in a FreeZone lyophilizer (Labconco, Kansas City, MO, USA). After freeze-drying, roots were flash frozen in liquid nitrogen. To break apart thick roots, sterile forceps and a dissecting needle were used before placing the rack of cluster tubes in an HT Lysing Homogenizer (OHAUS, Parsippany, NJ, USA) with two clean 5/32” steel balls in each tube. Samples were homogenized at 25 Hz for 1 min. Root material was then transferred to 2 mL bead-beating 96-well plates containing sterile 1 mm garnet beads with 850 µL of lysis buffer (1M Tris, pH = 8.0; 100 mM NaCl; 10 mM EDTA).

A positive (ZymoBiomics Microbial Community Standard) and negative control, (800 µL of lysis buffer) were included on each plate. Bead-beating plates were stored at −20°C until extraction. After thawing, 10 µL of 20% sodium dodecyl sulfate (SDS) was added to each well before homogenizing for a total of 20 min at 20 Hz. Plates were incubated in a water bath (55°C for 90 min) and centrifuged (6 min at 4500 x g). 400 µL of the resulting supernatant was transferred to new 1 mL 96-well plates containing 120 µL of 5 M potassium acetate in each well and incubated overnight at −20°C. After thawing the plates, they were centrifuged (6 min at 4500 x g) and 400 µL of the supernatant was transferred to a new 1 mL 96-well plate containing 600 µL of diluted SPRI-bead solution (protocol derived from^94^). These plates were mixed thoroughly for 5 min at 1000 r.p.m. on an orbital plate shaker. Samples were allowed to incubate for 10 min so DNA could bind to beads, after which the plate was centrifuged (6 min at 4500 x g) and placed on a magnet rack for 10 min. The supernatant was removed, and the beads were washed twice with 900 µL of 80% EtOH. After washing, the supernatant was decanted and the beads were air-dried. DNA was eluted in 75 µL of pre-heated 1x TE (pH = 7.5; 37°C) and transferred to clean 0.45 mL plates and stored at −20°C.

16S-v4 rRNA gene amplification was performed using paired 515f/806r primers^95,96^. PCR reactions contained DreamTaq Master Mix (Thermo Fisher Scientific, Waltham, MA, USA), 10 mg/µL bovine serum albumin (BSA), 100 µM peptide nucleic acid (PNA), and PCR-grade water. BSA was used to enhance PCR amplification while PNA was used to suppress primer binding and subsequent amplification of mitochondrial and chloroplast 16S regions^97^. The PCR included an initial denaturation step at 95°C for 2 min, followed by 27 cycles of an additional denaturation at 95°C for 20 s, PNA annealing at 78°C for 5 s, primer annealing at 52°C for 20 s, and extension at 72°C for 50 s. This was followed by a final extension step for 10 min at 72°C. PCR products were purified by incubating at 37°C for 20 min and then 15 min at 80°C after mixing with 0.78 µL of PCR-grade water, 0.02 µL of 10 U/µL exonuclease I (Applied Biosystems), and 0.2 µL of 1U/µL shrimp alkaline phosphatase (Applied Biosystems) per 10 µL PCR product. Two µL of the purified amplicons were used as template DNA in an indexing PCR to attach barcoded P5 and P7 Illumina adaptors. The 10 µL reaction included 5 µM each of P5 and P7 adaptors, 1x DreamTaq Master Mix (Thermo Fisher Scientific), 10 mg/mL BSA, 100 µM PNA, and PCR-grade water. An initial denaturation step was performed at 95°C for 2 min. For 8 cycles, an additional denaturation was carried out at 95°C for 20 s, PNA annealing at 78°C for 5 s, primer annealing at 52°C for 20 s, and extension at 72°C for 50 s. A final extension step was performed at 72°C for 10 min. PCR products were verified via 2% agarose gel electrophoresis and then pooled by 96-well plate using the “Just-a-Plate” PCR cleanup and normalization kit (Charm Biotech, St. Louis, MO, USA). Pools were size selected and combined in equimolar concentrations before being sequenced on an Illumina SP flow cell on the NovaSeq 6000 platform (2×250bp reads).

##### 3.6.2 Amplicon data processing

We removed primer sequences from raw 16S-v4 Illumina reads using cutadapt^70^, requiring at least 5 nucleotides of overlap. Additional quality control and processing was performed with the DADA2 software^98^. Forward reads were discarded if they had more than 6 expected errors, otherwise they were truncated at 200 nucleotides; for reverse reads the parameters were 7 expected errors and 170 nucleotides. Error rates were estimated separately for forward and reverse reads based on a sample of 1×10^8^ bases, then used to denoise and dereplicate reads using the standard DADA2 functions. Chimeric sequences were detected and removed using the “consensus” procedure in DADA2. Each individual sample was processed in parallel, after which all of the resulting amplicon sequence variant (ASV) tables were merged. Finally, taxonomy of each ASV was assigned by comparison to the RDP database^99^.

##### 3.6.3 Root bacterial community diversity analysis

All statistical analyses were performed using R (v4.4.0). Unfortunately, some samples were lost due to a 96-well plate being dropped during the sample DNA extraction and due to filtering of samples based on sequence quality. Ultimately, 156 gamagrass root microbiome and 276 maize root microbiome samples were included in these analyses. The cleaned and prepared microbiome data, as phyloseq objects, were loaded into R^68^. The phyloseq object sample data table was replaced with an updated metadata file. The “mock” samples were removed from the dataset and the data was subset by the test phase host, maize and gamagrass using phyloseq::subset_samples (v1.48.0). Maize and gamagrass root bacterial microbiome Shannon Diversity was fit to a mixed-effects model using the lme4 (v1.1-35.4) package^100^ and lmer() function, using the following model:

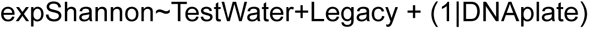

A type III ANOVA was performed using stats::anova()^101^ to test for significance, followed by pairwise comparisons and significance tests using emmeans (v1.10.2) package^102^ with false discovery rate adjustments. Data visualizations were generated using ggplot2 (v3.5.1)^45^, the error bars represent standard error and the points are estimated marginal means. All plots were saved in PDF format. Composite figures were created in Adobe Illustrator.

Bacterial community beta diversity was accessed with a Constrained Analysis of Principal Coordinates, using phyloseq::ordinate, method=”CAP”, and distance=”Euclidean” with the following formula:

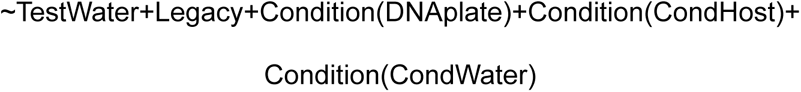

An ANOVA-like permutation test was used to test for model significance using the vegan::anova.cca() function^46^. The anova.cca by=term option was used to determine the significance of the TestWater and Legacy variables separately. CAP1 and CAP2 axes were plotted using ggplot2, including 95% confidence interval ellipse (stat_ellipse()). To evaluate treatment group beta-dispersion, Euclidean distances were calculated for the centered-log ratio transformed counts using phyloseq::distance(). Both “Legacy” and “TestWater” terms were tested for beta-dispersion differences using the Euclidean distances matrix and vegan::betadisper() and vegan::permutest() to test for significance^46^. Distances were then extracted from the results output and fit to a linear model. Estimated marginal means were calculated and plotted using the same methods as above. The error bars represent standard errors.

Then, an ASV differential abundance analysis was performed on gamagrass- and maize-associated ASVs separately. Centered-log ratio transformed counts of ASVs were each fit to a linear model using an iterative for-loop. Each ASV was subset using phyloseq:subset_taxa and the phyloseq::psmelt() function was used to reformat the phyloseq object into a data frame. Then the ASV abundance was fit to the following model:

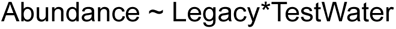

Then, stats::anova() was used to test for significance, followed by an FDR p-value adjustment using stats::p.adjust(). ASVs with significant (≤0.05) adjusted p-values for the Legacy*TestWater interaction terms were extracted and z-scores of the center-log ration transformed counts were plotted in a heatmap using pheatmap::pheatmap()^103^ with clustering_method=”complete”.

#### 3.7 Plant trait data analyses

##### 3.7.1 Plant trait feature selection

All plant data was loaded into RStudio. The intrinsic water use efficiency (iWUE) was calculated using the ratio of photosynthetic rate (A) and stomatal conductance (gs) or A/gs ((μmol m^−2^ s^−1^)/(mol m^−2^ s^−1^)). Data was subset by Test Phase host treatment groups (gamagrass and maize). For the feature selection, the “mock” treatment was removed. The 67 features that were selected for testing were subset into a data frame. Three sample rows were removed from the dataset because they had missing data. The drought susceptibility index (S-index) was calculated for each of these features using the following formula:

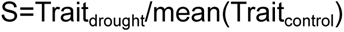

Then, for each host, a random forest model was used to select the top 10 most important features from the 67 total s-index measurements using randomForest()^104^, with “Legacy” as the predictor. Correlations (stats::cor()) were estimated for the top ten traits and if any two of the top 10 traits were highly correlated (*r* ≥ 0.7), the trait that ranked lower in the random forest model was removed. Non-correlative top features from the test water treatment groups (drought vs. watered control) were combined for each test phase host (maize and gamagrass). Vegan:rda() was used to perform a redundancy analysis on the top features using the following formula:

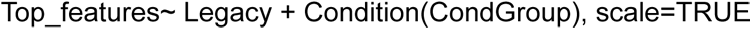

##### 3.7.2 Feature analyses

An ANOVA-like permutation test was used to test for model significance using the vegan::anova.cca() function. A biplot was created using ggplot2 plotting RDA1 and PC1 axes, as well as the species scores (represented by arrows), and stat_ellipse was used to plot 95% confidence intervals of the site scores.

Finally, s-indices for each of the top features and iWUE were fit to mixed-effects models, when possible, or a fixed-effects model if overfitting of the mixed effects model occurred. Each feature or trait was visually assessed for outliers, which were removed. Removing the outliers did not impact the interpretation of any of the results. Formulas used for the mixed effects or linear model, respectively, include:

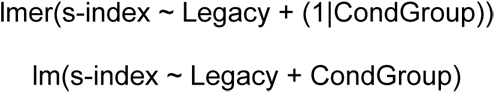

The fit of the model was accessed and if needed, the s-index was transformed using sqrt(), exp(), or log() to improve the fit. ANOVA was used to assess significance and estimated marginal means were calculated and plotted as described above.

## Data availability

The *16S rRNA* gene amplicon sequencing data, shotgun metagenomic data, and metatranscriptome data associated with this study have been deposited in the NCBI Sequence Read Archive under the BioProject IDs PRJNA1267293, PRJNA1267715, PRJNA1268489, and PRJNA1186942. The raw RNA-seq data from gamagrass and maize have been deposited in the Gene Expression Omnibus under accessions GSE282586 and GSE282587, respectively.

## Code availability

We deposited all scripts and additional data structures required to reproduce the results of this study in a Zenodo repository (http://doi.org/10.5281/zenodo.13821006). Source data are provided in the Zenodo repository and with this paper.

## Acknowledgements

We thank Dr. Arun Seetharam, Dr. Toby Kellogg, Dr. Matthew Hufford, and the PanAnd consortium for granting early access to the *Tripsacum dactyloides* genome and annotation. We thank Dr. Mitch Greer, Michaela VonLintel, Justin Roemer, Dr. Alan Schlegel, and the Nature Conservancy for granting access to soil collection sites in Kansas, and Dr. Peter Balint-Kurti for providing maize seeds. We thank Dr Craig J. Sturrock and Dr Brian S. Atkinson from the Hounsfield Facility at the University of Nottingham for their assistance with the analysis of the CT images. We are grateful to the undergraduate student researchers and technicians who helped to set up experiments and collect data: Felicity Tso, Hannah Reid, and Carmen Rodriguez. Thank you to Dr. Joel Swift, Ceyda Kural-Rendon, Dr. Ben Sikes (University of Kansas), Dr. Manuel Kleiner (North Carolina State University), Dr Omri Finkel (The Hebrew University of Jerusalem, Israel), and Dr Nicholas Girkin (University of Nottingham, UK) for critical reading of the manuscript. This work was supported by a UKRI/BBSRC - NSF/BIO Lead Agency Opportunity, National Science Foundation award #IOS-2016351 and BBSRC grant #BB/V011294/1. Maize root transcriptome library preps and sequencing were performed in the RTSF Genomics Core at Michigan State University. *T. dactyloides* library preps and RNA-seq were performed at the NC State University Genomic Sciences Laboratory (Raleigh, NC, USA).

Research reported in this publication was made possible in part by the services of the KU Genome Sequencing Core. This lab is supported by the National Institute of General Medical Sciences (NIGMS) of the National Institutes of Health under award number P30GM145499. We thank the University of Kansas Medical Center – Genomics Core for generating the 16S amplicon sequence data sets. The Genomics Core is supported by the Kansas Intellectual and Developmental Disabilities Research Center (NIH U54 HD 090216), the Molecular Regulation of Cell Development and Differentiation – COBRE (P30 GM122731), The NIH S10 High End Instrumentation Grant (NIH S10OD021743) and the Frontiers CTSA Grant (UL1TR002366).

## Author contributions

Conceptualization: N.A.G., V.C., D.G., G.C., and M.R.W. Data curation: N.A.G., V.C., D.G., and M.R.W.; Formal analysis: N.A.G., V.C., D.G., I.S.-G., and M.R.W.; Investigation: N.A.G., V.C., D.G., N.F., D.D.J., D.M.W., A.M., G.C., and M.R.W.; Methodology: G.C. and M.R.W.; Project administration: G.C. and M.R.W.; Software: N.A.G., V.C., D.G., and M.R.W.; Supervision: G.C. and M.R.W.; Visualization: N.A.G., V.C., D.G., and M.R.W.; Writing – original draft: N.A.G., G.C., and M.R.W.; Writing – review & editing: V.C. and D.G.

## Extended Data Figures

**Extended Data Fig. 1.**
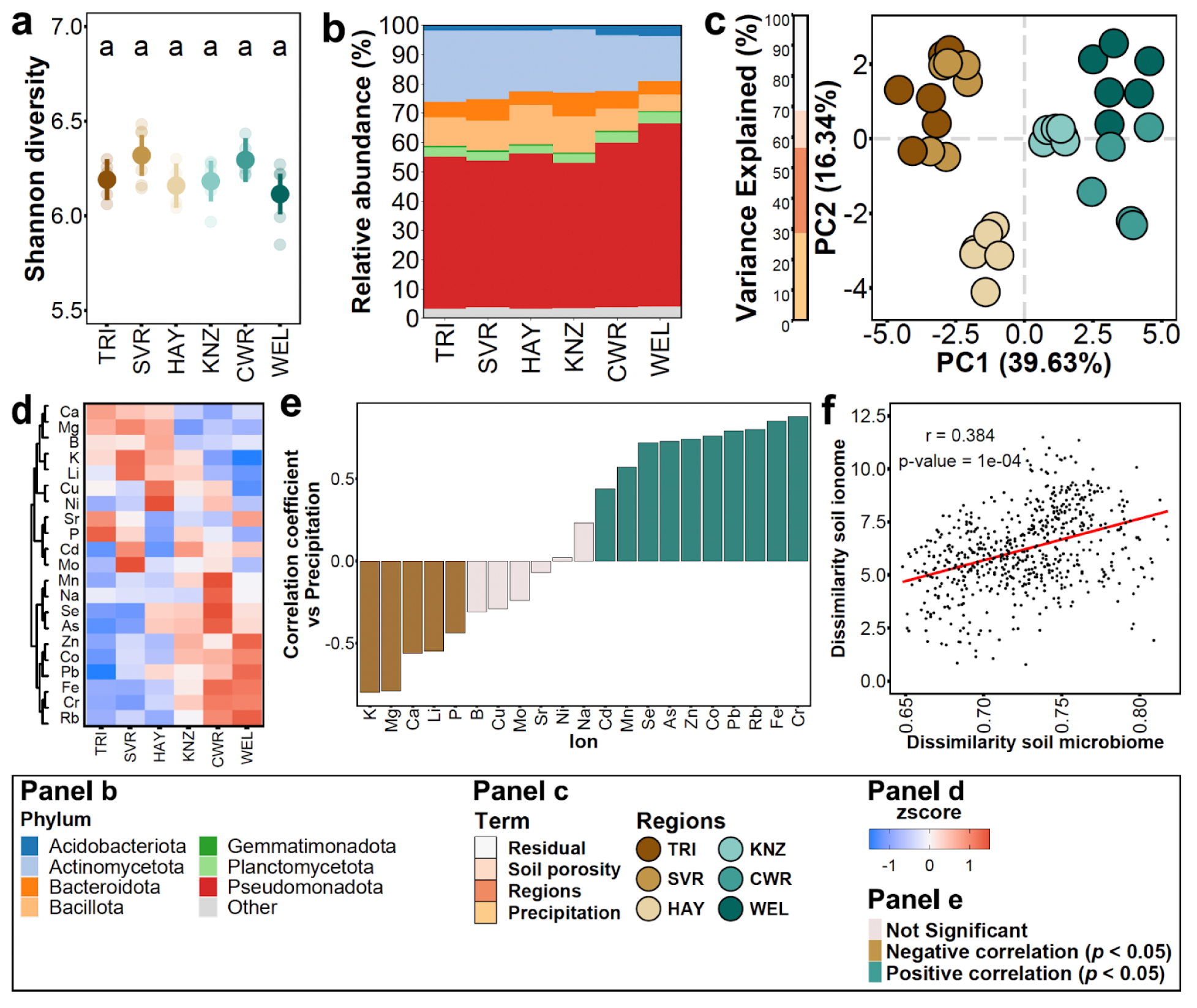
Precipitation level and mineral nutrient content in Kansas soils correlate with soil microbiota composition. **a.** Alpha diversity (estimated using the Shannon diversity index) did not differ across soil samples. Soils are ordered according to the gradient of precipitation (Fig 1a). **b**. Phylogram showing the relative abundance profiles of the main bacterial phyla across soils exposed to different precipitation levels. **c**. Principal component analysis showing the ionomic profiles of the six focal soils (N=6 per soil). The bar on the left denotes the percentage of the variance explained by the predictor variables. **d**. Heatmap showing the standardized concentration (z-score) of each mineral nutrient (rows) in the collection of soil exposed to different precipitation levels. The values were clustered according to the ion concentration and the region was ordered according to the precipitation gradient (low to high). **e**. Bar graph showing the Pearson correlation coefficient between each mineral nutrient abundance and the level of precipitation across the collection of soils used. Coloured bars indicate statistically significant correlations (*q* < 0.05). **f**. Pairwise correlation analysis between soil mineral nutrient dissimilarities and soil microbiome composition dissimilarities in soils representing a gradient of precipitation (N=6 per soil). Each point represents one pair of soil samples. Panel shows the Mantel test *r* statistic and its *p*-value.

**Extended Data Fig. 2.**
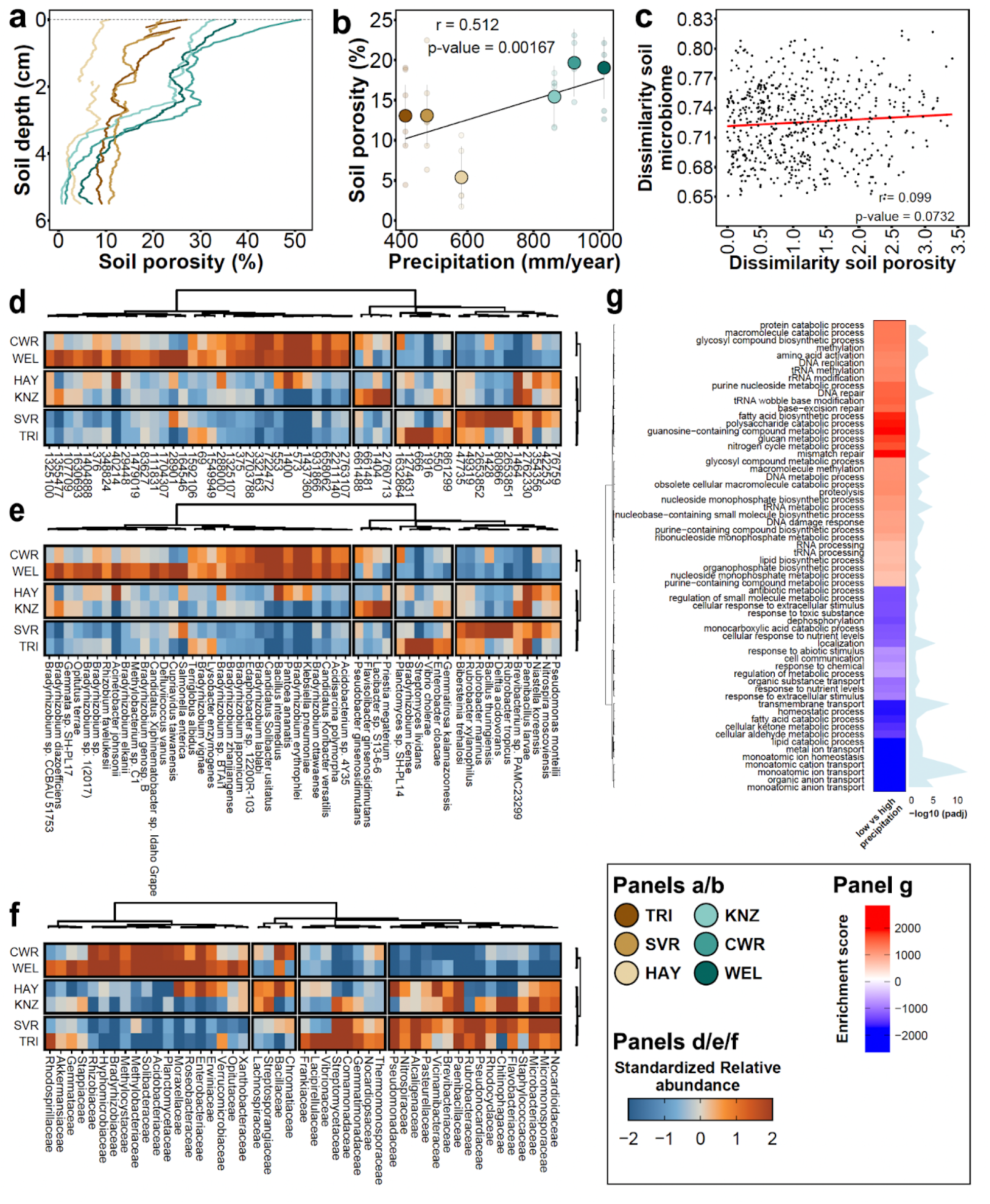
Precipitation, but not soil porosity, explains microbiota composition across the Kansas soil collection. **a.** Percent soil porosity changes with depth in soils exposed to the precipitation gradient. **b**. Pearson correlation analysis between soil porosity (averaged across depths) and mean annual precipitation. **c**. Pairwise correlation analysis between soil microbiota dissimilarities and soil porosity dissimilarities. The panel shows the Mantel r statistic and its p-value. **d-f.** Heatmaps showing changes in the relative abundances of bacterial taxa (NCBI TaxIDs, **d**), species (**e**), and family (**f**) relative to the WEL soil, from the highest-precipitation site. In all cases, the values have been clustered according to taxonomic categories and soils. **g**. Numerous biological processes were enriched (red) or depleted (blue) in soils from low-precipitation sites (TRI, SVR, and HAY) relative to high-precipitation sites (WEL, CWR, and KNZ) (q<0.05). Gene enrichment analysis was conducted using a generalized linear model, followed by Gene Ontology (GO) classification. Enrichment scores were calculated using square root-transformed delta rank values of the GO categories.

**Extended Data Fig. 3.**
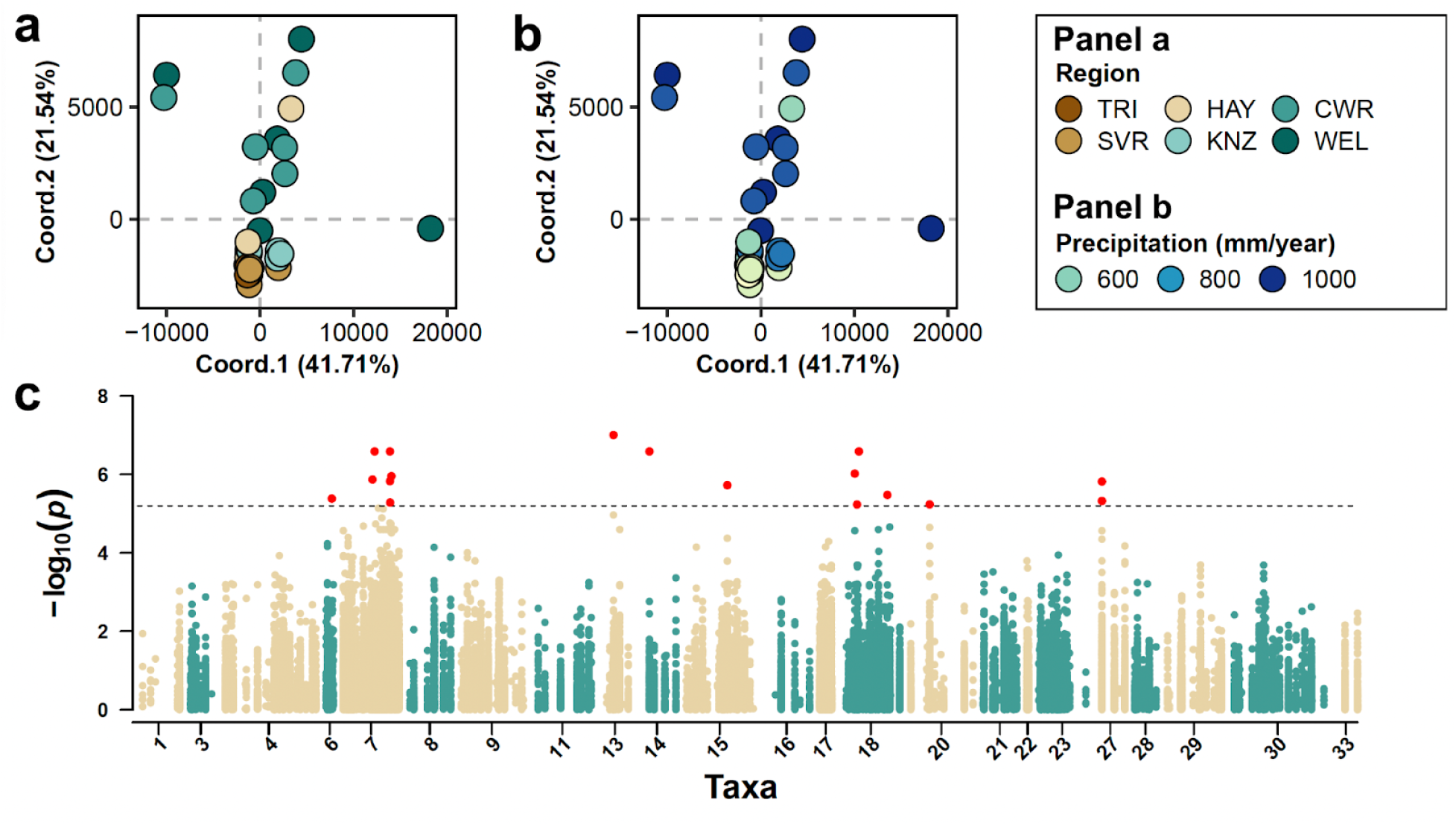
Precipitation legacy shapes the genetic differentiation of bacterial lineages within taxa. To assess genetic differences between bacterial lineages along the precipitation gradient, we selected 33 bacterial species (15 of the bacterial biomarkers of precipitation, plus 18 additional abundant and prevalent taxa) that exhibited high genome coverage in our metagenomic dataset as proxies for the broader bacterial communities. Reference genomes for each species were retrieved from the NCBI Genome database, and filtered shotgun metagenomic reads were mapped to these genomes to identify high-quality biallelic single nucleotide polymorphisms (SNPs). Genetic distances between the bacterial lineages were calculated based on the identified SNPs and PCoA plots were generated and coloured by **a**. soil collection sites and **b**. mean annual precipitation. The variance explained by each axis is indicated. **c.** The Manhattan plot illustrates significant SNPs associated with precipitation, derived from the genetic-environment association (GEA) analysis. The GEA was conducted using a general linear model, with precipitation at each sampling location as the environmental variable. Significant associations were identified using the permutation method. The x-axis of the plot represents the SNP positions along the genomes of the selected bacterial species, while the y-axis displays the -log_10_ p-values from the association model. The horizontal line indicates the statistical significance threshold, as determined by the permutation test. SNPs above this threshold, highlighted in red, were significantly associated with the precipitation gradient. Bacterial taxa with fewer than 1,000 high-quality biallelic SNPs after filtering, and with no significant SNPs detected from the GEA analysis, are not shown in the plot.

**Extended Data Fig. 4.**
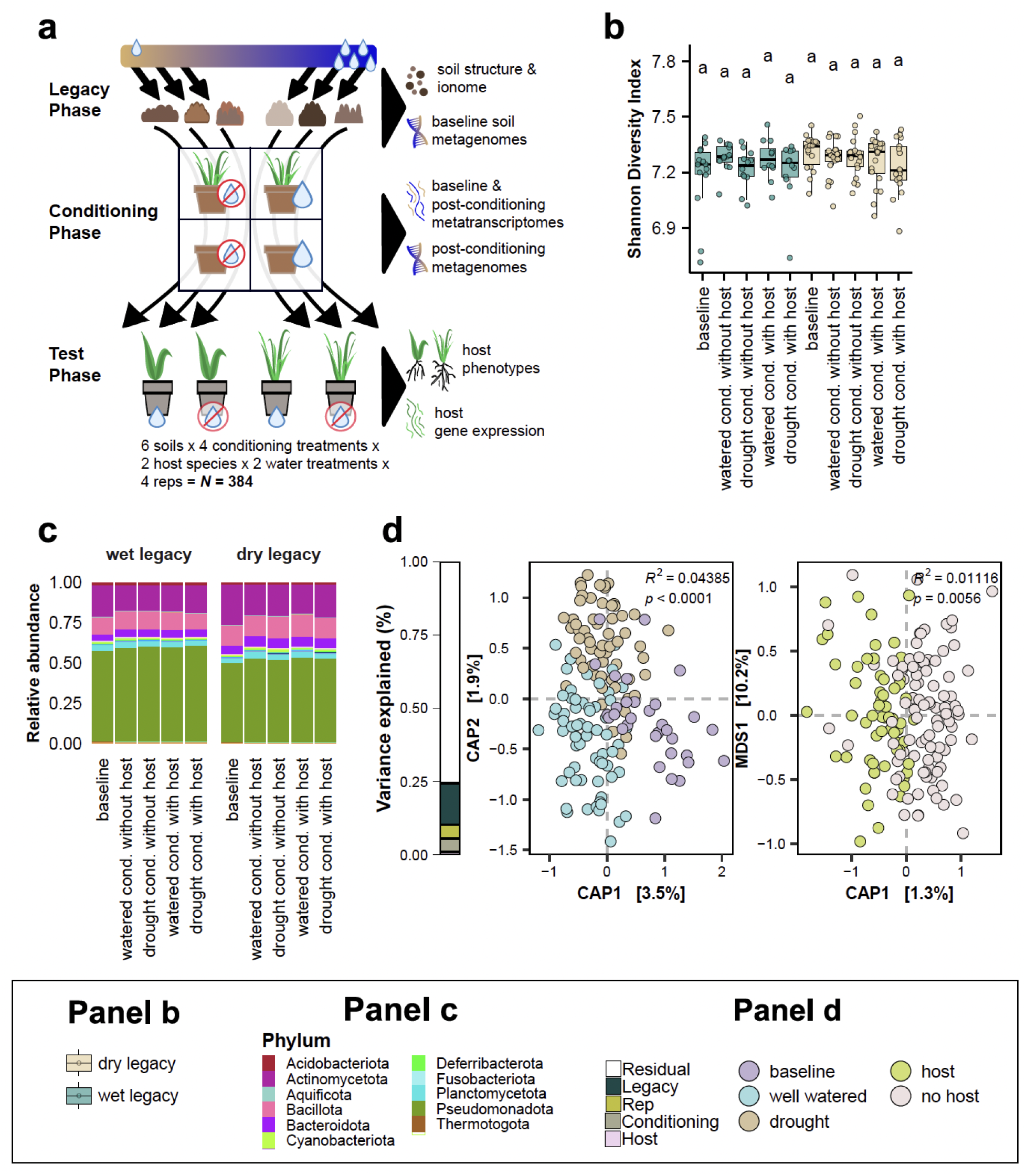
Precipitation legacy effects in the soil microbiota are resilient to short-term water- and host-related perturbations. **a**. Schematic representation of the experimental design used to evaluate the resilience of precipitation legacy effects to perturbations and their functional importance to plant drought response. Six soils spanning the Kansas precipitation gradient (“legacy phase”), were either left unplanted or planted with seedlings of the native grass species *Tripsacum dactyloides* (eastern gamagrass), and subjected to either drought conditions or regular watering in a factorial design (“conditioning phase”). To evaluate how soil precipitation legacy affects plants, and to disentangle the role of the microbiota from possible effects of co-varying abiotic soil properties, we then used the experimentally-conditioned microbial communities to inoculate a new generation of *T. dactyloides* and *Z. mays* plants. These “test phase” plants were divided between water-limited conditions and well-watered control conditions. **b**. Alpha diversity of bacterial communities was not affected by the different conditioning treatments (drought or well-watered, with or without host). **c**. Phylograms show that the different conditioning treatments (drought or well-watered, with or without host) did not impact the relative abundance profiles of main bacterial phyla. **d**. Constrained ordination of metagenome taxonomic composition in response to conditioning phase treatments. Statistics are from permutational MANOVA. The bar on the left describes the percentage of the variance explained by the experimental variables.

**Extended Data Fig. 5.**
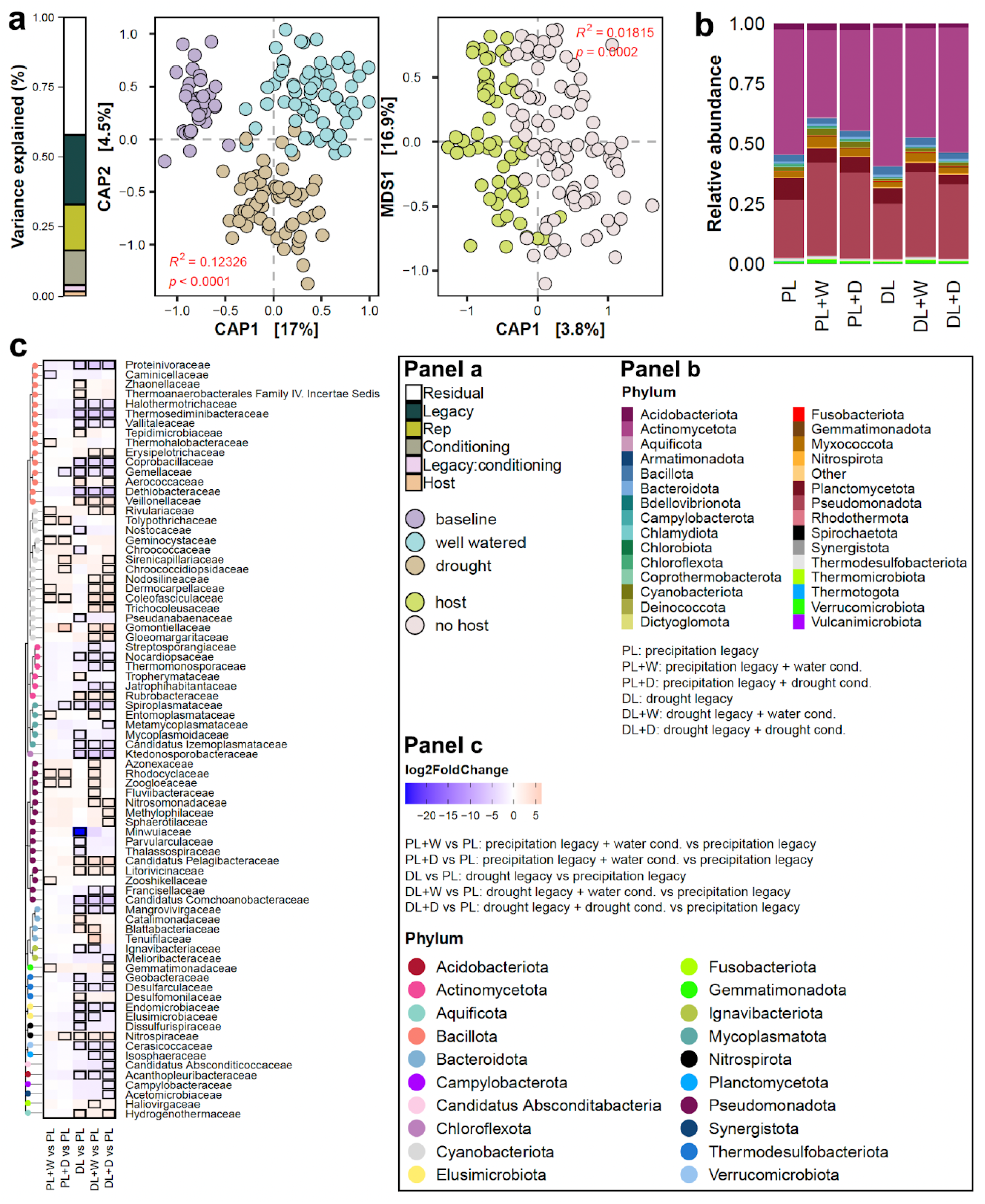
Precipitation legacy effects shape the transcriptionally-active soil microbiota even after five months of experimental perturbation. **a**. Constrained ordination of metatranscriptome content in response to conditioning phase treatments: (left) baseline soils and soils after exposure to well-watered or drought treatment, and (right) host or no-host treatment. The bar on the left describes the percentage of the variance explained by the experimental variables. **b**. Phylogram showing the main bacterial phyla that were transcriptionally active across soils with different precipitation legacies after five months of drought or well-watered conditions, compared to the baseline for each legacy group (PL and DL). **c**. Heatmap showing enrichment or depletion of transcriptionally active bacterial families relative to the baseline high-precipitation-legacy (PL) soil. DL indicates the baseline (pre-conditioning) low-precipitation-legacy soil; +W and +D indicate five months of well-watered or drought conditions, respectively. Heatmap was coloured based on log_2_ fold changes derived from a generalised linear model contrasting the abundance of each family in a given treatment against the high-precipitation-legacy baseline soil. Tiles outlined in black denote statistically significant enrichment (red) or depletion (blue) (q < 0.05) with a |log_2_ fold change| > 2. Heatmaps were clustered based on taxonomic classification (tree on the left).

**Extended Data Fig. 6.**
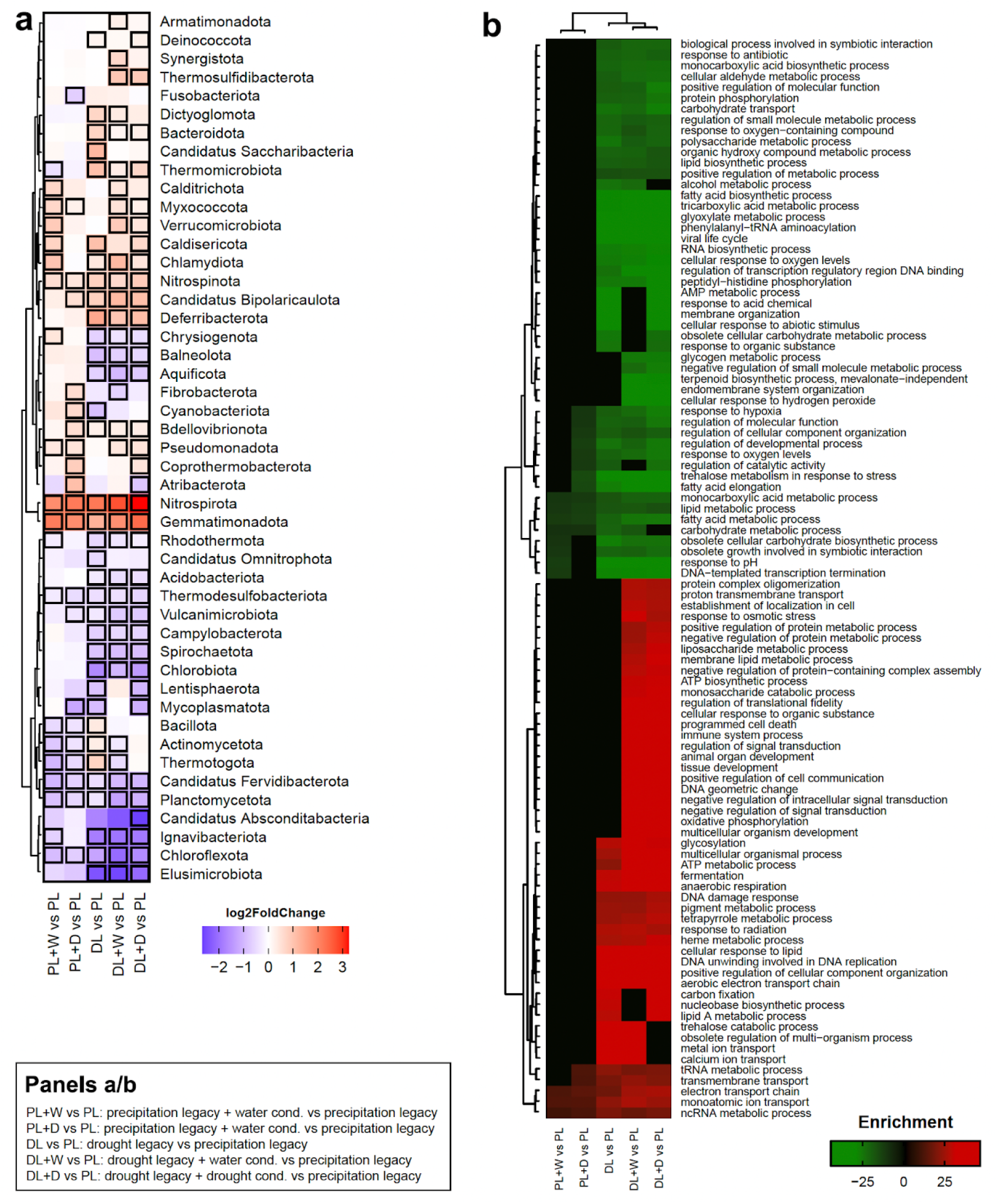
Transcriptional responses of soil microbiomes to short-term water perturbations are shaped by precipitation legacy. **a**. Heatmap showing the enrichment of transcriptionally active bacterial phyla in soils with low-precipitation (DL) or high-precipitation (PL) legacies, exposed to either drought (+D) or well-watered (+W) treatments, relative to the high-precipitation-legacy baseline (PL). DL indicates the baseline (pre-conditioning) low-precipitation-legacy soil. Colours represent log₂ fold changes derived from a generalized linear model comparing each treatment to the high-precipitation-legacy baseline (PL). Tiles outlined in black indicate statistically significant enrichment (red) or depletion (blue) (q < 0.05; |log₂ fold change| > 2). Taxa were hierarchically clustered based on taxonomic classification (dendrogram on the left). **b**. Heatmap depicting enriched or depleted biological processes identified through metatranscriptomic analysis. Soils with high-precipitation legacies (PL) or low-precipitation legacies (DL) were subjected to five-month-long drought or well-watered treatments and then compared to the high-precipitation-legacy baseline. Gene enrichment analysis was conducted using a generalized linear model followed by Gene Ontology (GO) classification. Significantly enriched or depleted GO categories (adjusted p < 0.05) are coloured according to enrichment scores, calculated from square root-transformed delta rank values (red: enrichment; green: depletion). Clustering was performed based on soil treatments and GO terms.

**Extended Data Fig. 7.**
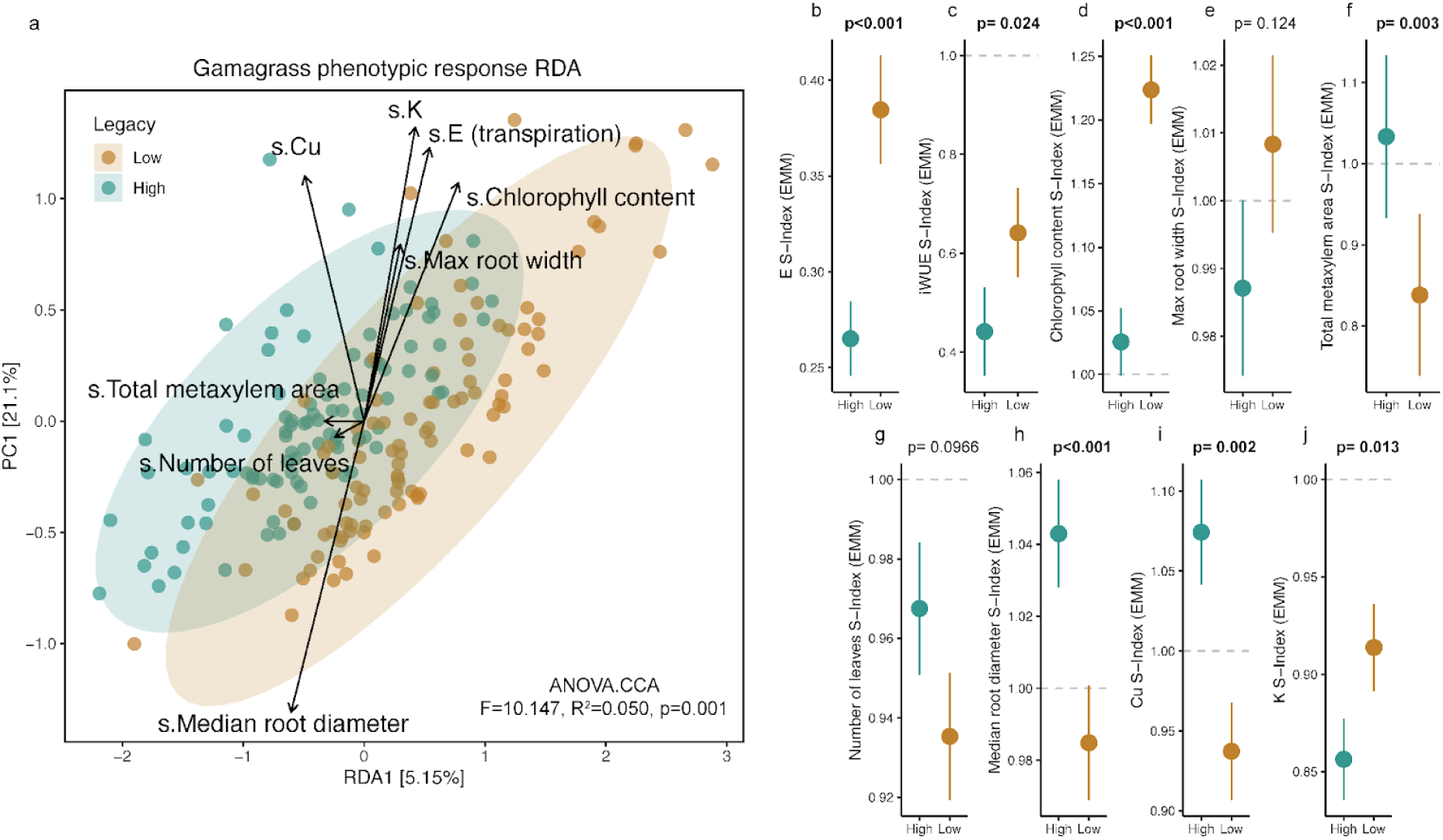
Precipitation legacy of microbial inoculum impacts gamagrass phenotypic drought response. **a.** A constrained redundancy analysis of the top non-collinear traits found that legacy explains 5.0% of the phenotypic response to acute drought in eastern gamagrass (*Tripsacum dactyloides*). Turquoise points represent plants that were inoculated with high-precipitation-legacy microbiota and brown points represent plants that were inoculated with low-precipitation-legacy microbiota. The ellipses indicate 95% confidence intervals. **b-j**. Assessment of individual traits indicates that microbiota with a low-precipitation legacy improved gamagrass performance under drought. Points are estimated marginal means and error bars represent the standard error, and significant p-values (≤0.05) are bolded.

**Extended Data Fig. 8.**
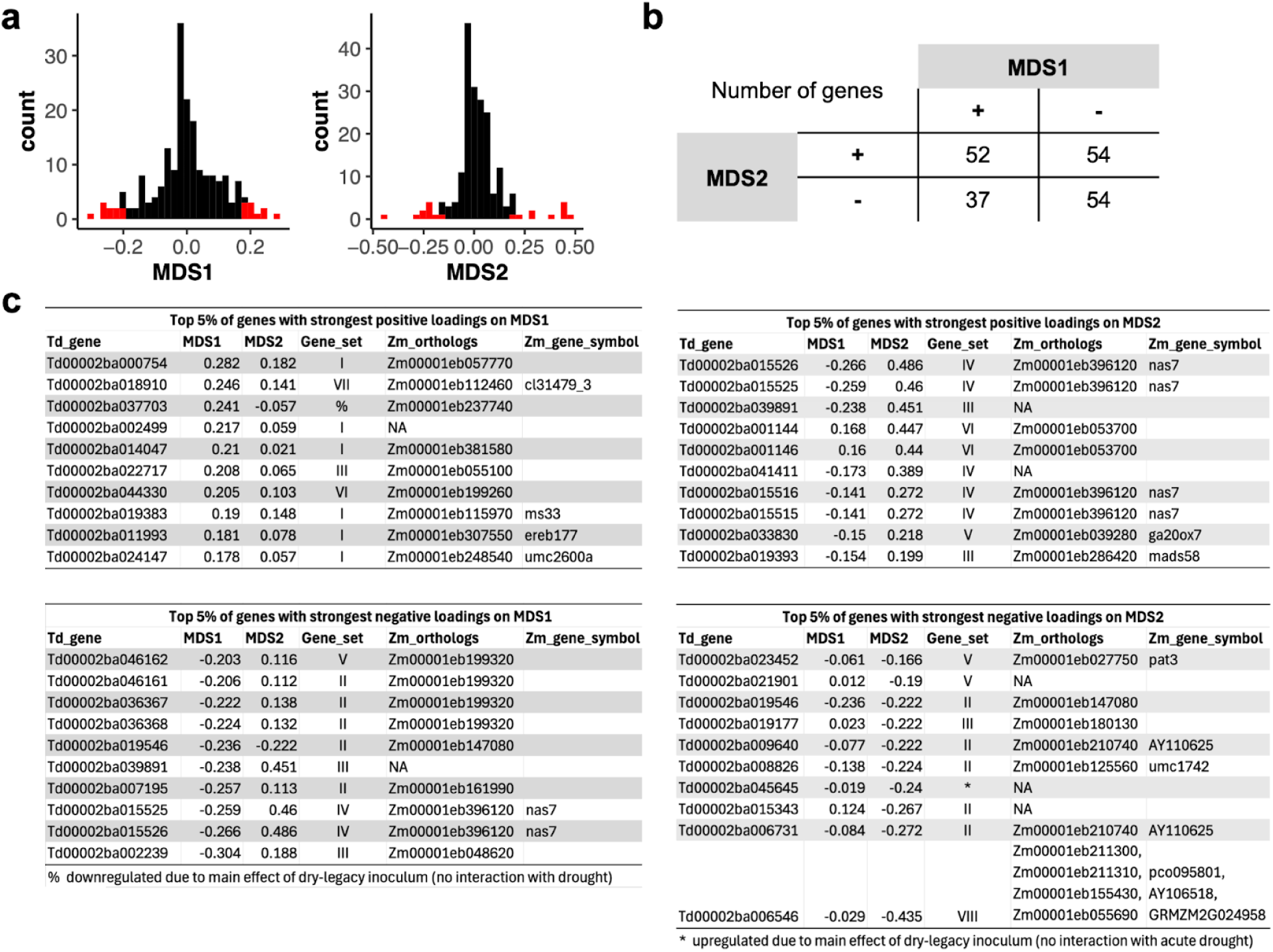
Expression patterns of *Tripsacum dactyloides* genes in relation to microbiome legacy and NMDS axes that mediate drought responses. **a**. Histograms showing the distributions of gene loadings onto both axes of an ordination based on non-metric multidimensional scaling of RNA-seq data (see Fig. 4). The top 5% of genes with the strongest positive and negative loadings onto each axis are in red. **b**. Breakdown of the number of genes with positive/negative loadings onto each axis. **c**. Lists of genes comprising the 5% tails of the distributions in panel (a). Detailed information about the expression responses of these genes is available in Supplementary Table S10. ‘Gene_set’ refers to the patterns of response to microbiome legacy presented in Fig. 4a-b.

**Extended Data Fig. 9.**
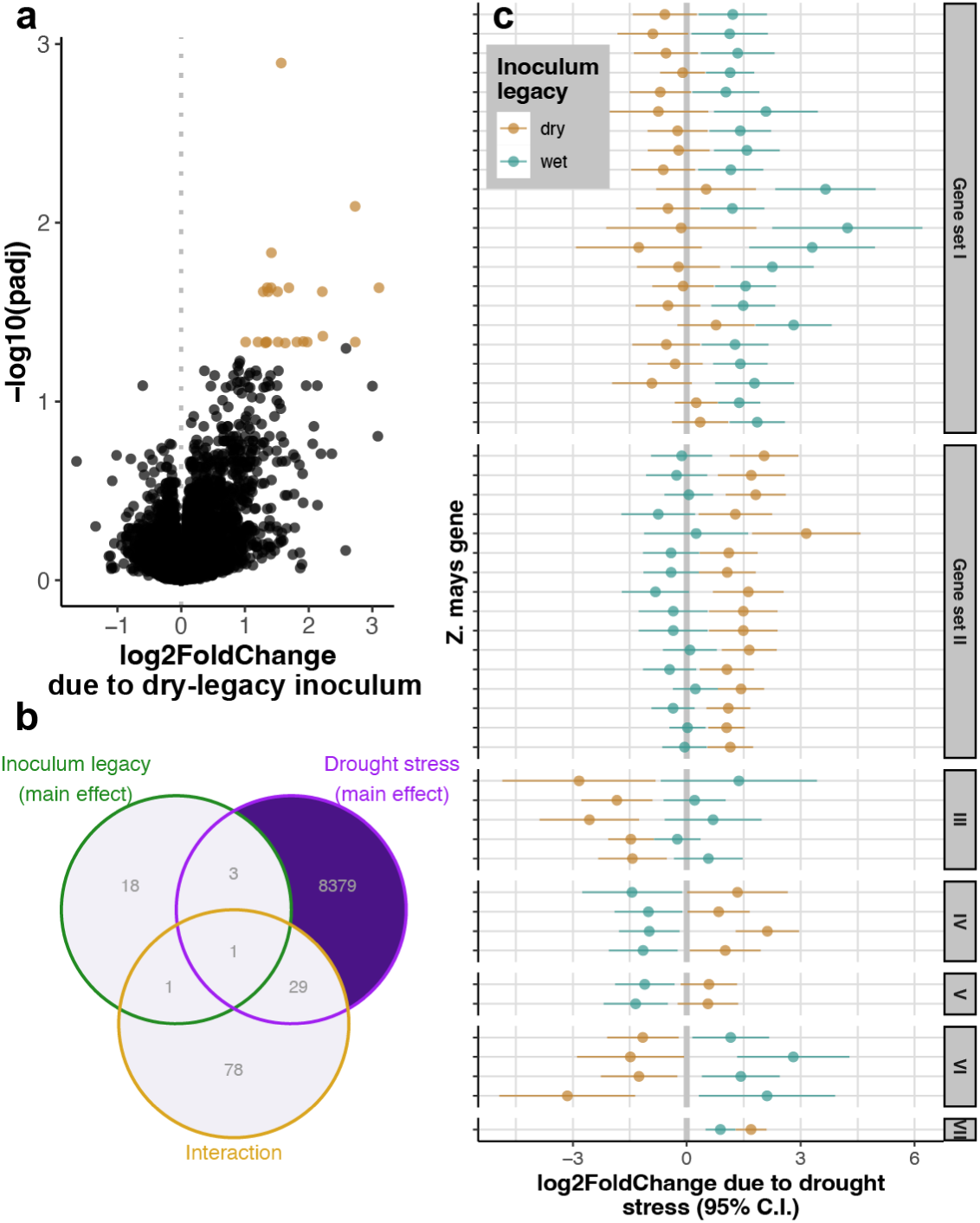
Precipitation legacy of microbial inocula alters the transcriptional response to drought in the maize crown root. **a.** 23 genes were up-regulated in plants inoculated with soil microbiota from a low-precipitation region, relative to plants inoculated with microbiota from a high-precipitation region. **b.** The sets of genes that responded to the main effects of inoculum legacy and test phase drought treatment had little overlap with each other or with the set of genes that were sensitive to the interaction between the two. **c.** In total, 109 maize genes responded to drought in a manner that was dependent on the drought legacy of the soil microbiota (the inoculum legacy * drought treatment interaction term), regardless of the inoculum’s treatment during the conditioning phase. For illustration purposes, only annotated genes with |log_2_FoldChange| > 1 in at least one microbial context are shown here; the full list is available in Supplementary Table S10. Each pair of points shows one gene; the position of each point illustrates how the gene’s expression changed in response to drought stress during the Test Phase, depending on whether the plant had been inoculated with microbiota derived from a low-precipitation (brown) or dry-precipitation (turquoise) environment. Genes are grouped into sets according to the pattern of how inoculum legacy altered their drought responses. Note: the names of these gene sets are not meant to correspond to the names of the *T. dactyloides* gene sets shown in Fig. 4b; in each species, Gene set I contains the most genes, Gene set II contains the next most, and so on.

**Extended Data Fig. 10.**
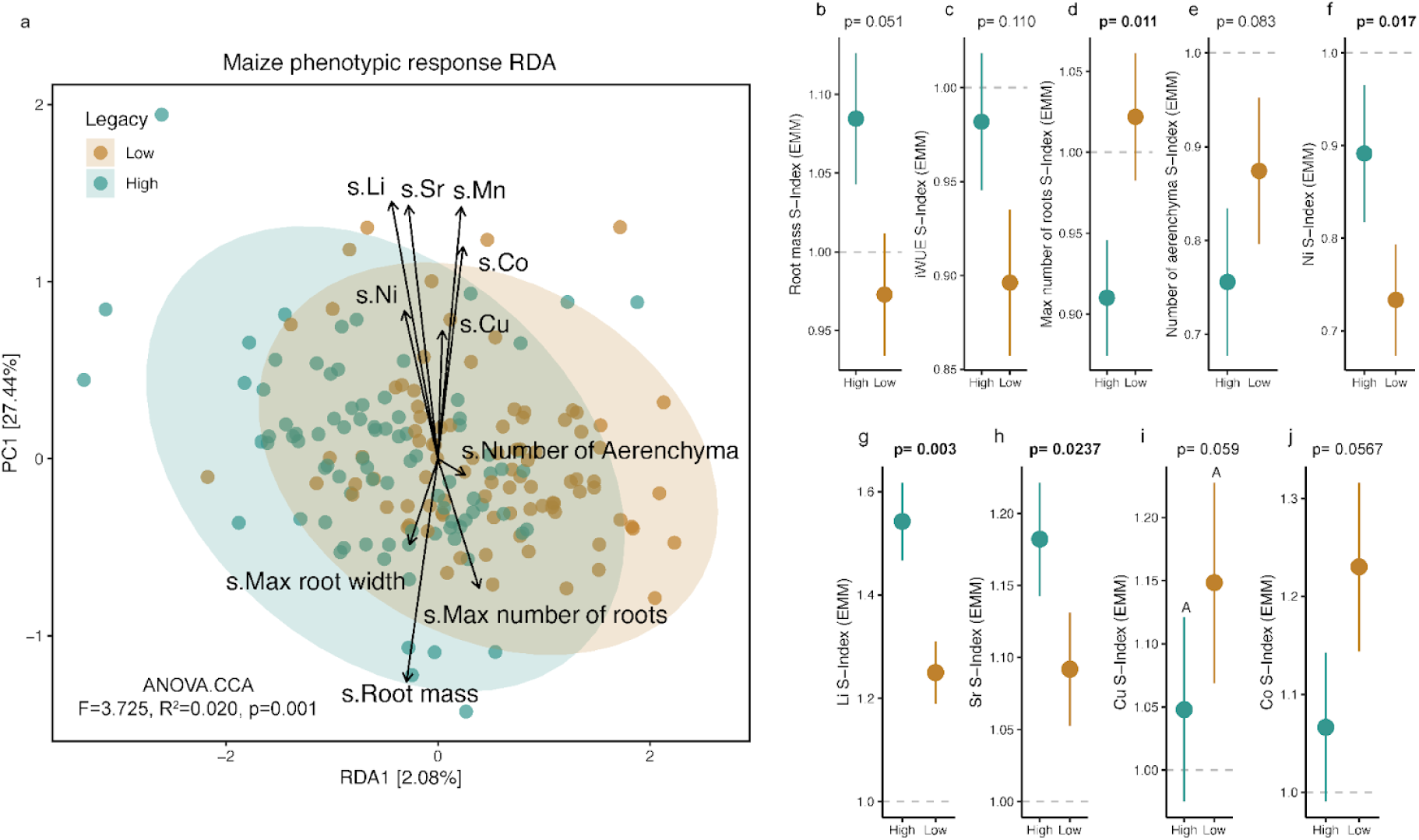
Precipitation legacy of the microbiota had only weak impacts on the phenotypic drought response of maize, compared to that of eastern gamagrass. For comparison, the effects of microbiota precipitation legacy on eastern gamagrass drought responses are shown in Fig. 4 and Extended Data Fig. 7. **a.** A constrained redundancy analysis of the top non-collinear traits found that the precipitation legacy of the microbial inoculum only explains 2.0% of the maize phenotypic response to acute drought. Turquoise points represent root microbiomes with high precipitation legacy and brown points represent those with low precipitation legacy. The ellipses indicate 95% confidence intervals. **b-j.** Assessment of individual traits indicates that microbiota with a low-precipitation legacy did not significantly improve maize performance under drought, but did impact several mineral nutrient concentrations. Points are estimated marginal means and error bars represent the standard error, and significant *p*-values (≤ 0.05) are bolded.

