## Supplementary Information for "Persistent legacy effects on soil microbiota facilitate plant adaptive responses to drought"

**This PDF file includes:**

Supplementary Fig. 1-8

Supplementary Tables 1-12 (Legends)

Supplementary Note 1

### Supplementary Figures

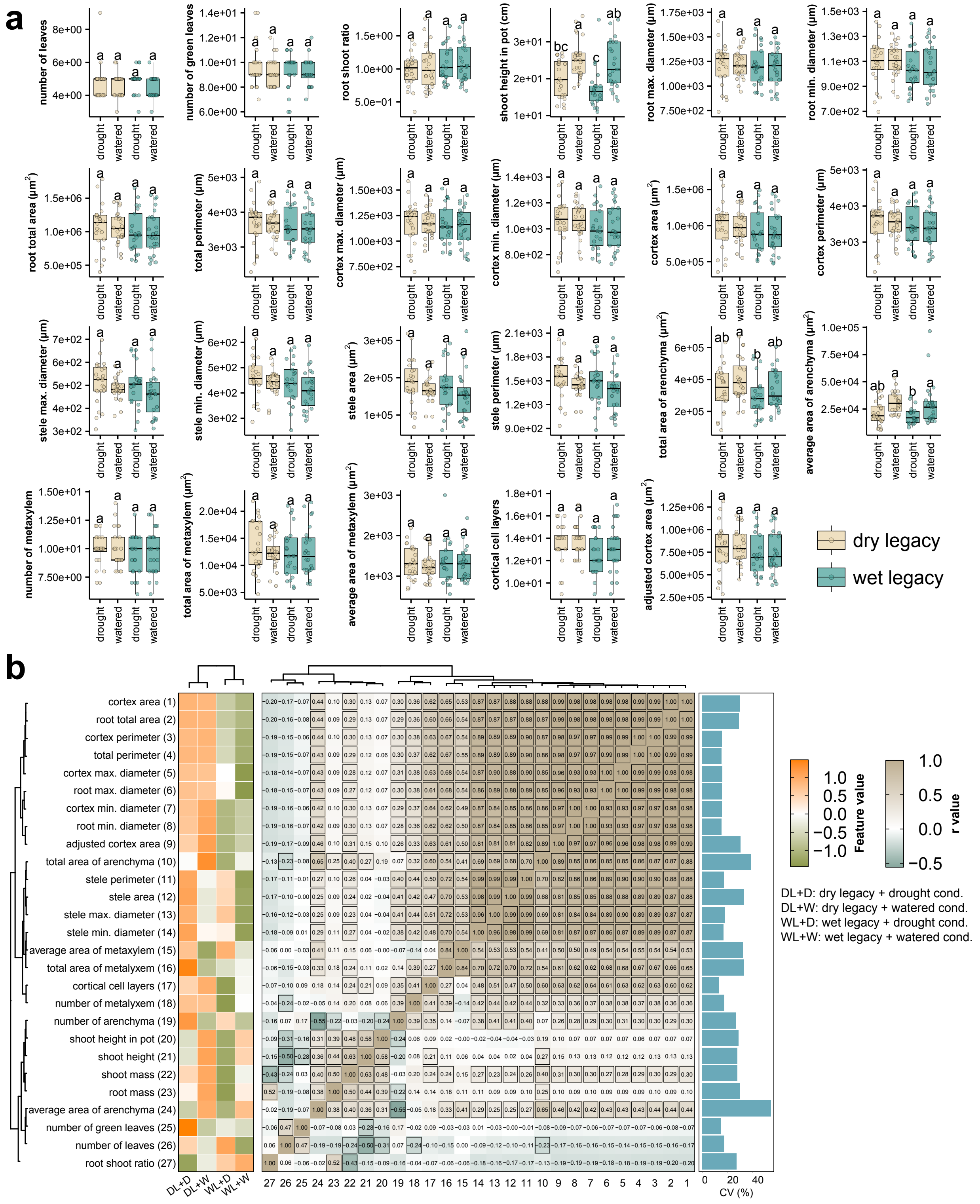


**Supplementary Fig. 1. Precipitation legacy effects in the soil microbiota are resilient to short-term water- and host-related perturbations. a**. Box plots showing how soil group (high-precipitation or low-precipitation legacy) and treatment (droughted or well-watered) affect phenotypic distributions of the *Tripsacum dactyloides* plants grown during the conditioning phase. ANOVA was used to partition variance among groups and estimated marginal means were compared using Tukey's post hoc test. **b**. Effects of soil group (high-precipitation or low-precipitation legacy) and treatment (droughted or well-watered) on phenotypic traits of *Tripsacum dactyloides* plants grown during the conditioning phase. The heatmap on the left depicts key plant phenotype features, coloured based on mean normalised values. The Pearson correlation coefficients (r) between these features are presented in the centre heatmap, with significant correlations ($p < 0.05$) highlighted by black squares. The bar plot on the right indicates the coefficient of variation of the feature values. The heatmaps are clustered based on Pearson correlation coefficient values.

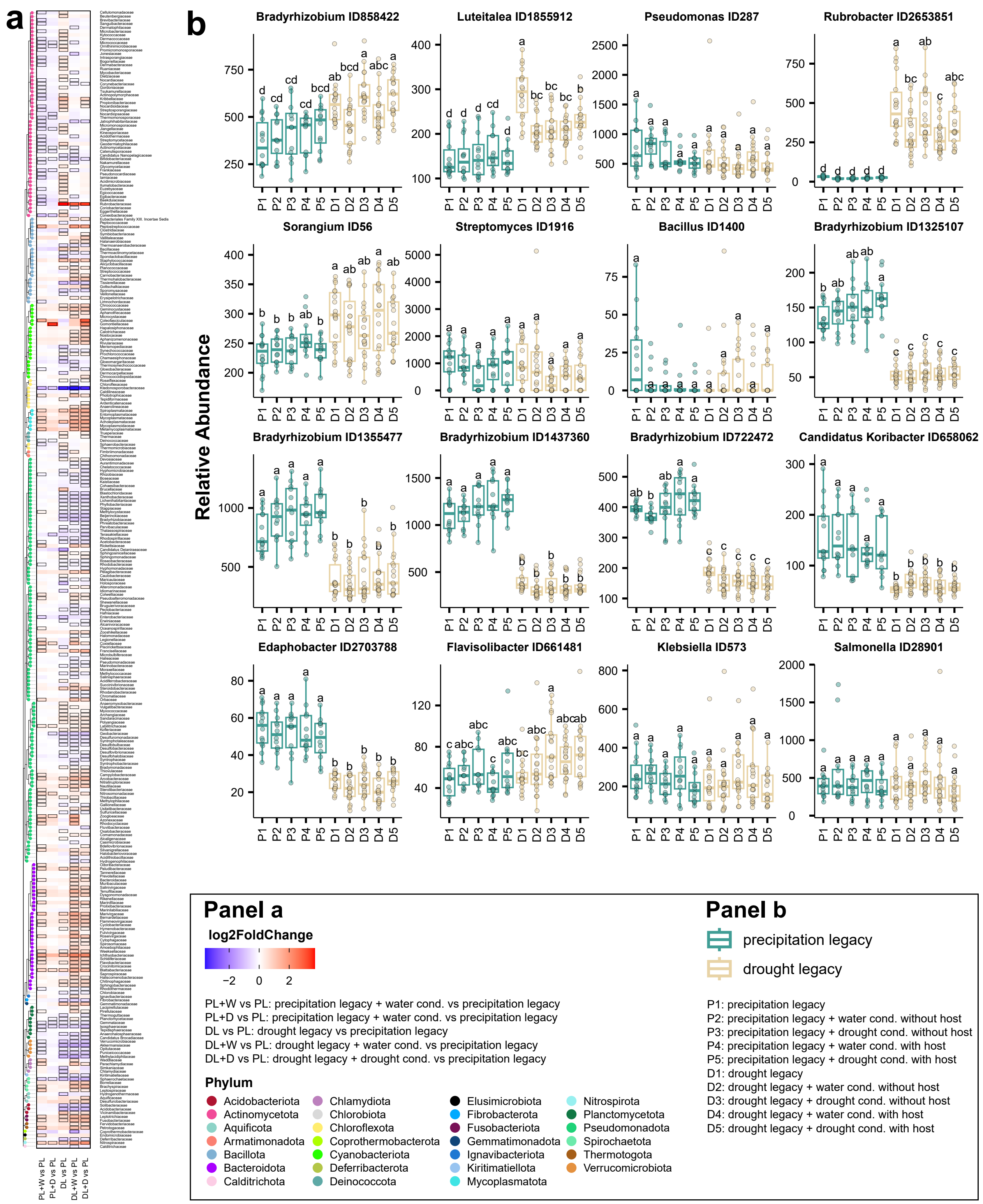


**Supplementary Fig. 2. Limited influence of drought and watering treatments on soil microbiota with established precipitation legacies. a**. The heatmap displays the enrichment of bacterial families across soils with high-precipitation (PL) and low-precipitation (DL) legacies, exposed to either drought (+D) or well-watered (+W) treatments, in comparison to the high-precipitation soil baseline (PL). DL indicates the baseline (pre-conditioning) low-precipitation-legacy soil. Colours in the heatmap represent $\log_2$ fold changes, calculated using a generalized linear model that contrasts the abundance of each bacterial family in the respective treatments with that of the high-precipitation soil baseline (PL). Tiles outlined in black denote statistically significant enrichment (red) or depletion (blue) ($q < 0.05$) with a $|\log_2 \text{fold change}| > 2$. The heatmap was clustered based on taxonomic classification (represented by the dendrogram on the left). **b**. Boxplots showing the relative abundance of bacterial markers of water legacy across soils with low-precipitation legacy (D1-5) or high-precipitation legacy (P1-5) either before (P1, D1) or after (P2-5, D2-5) exposure to conditioning phase treatments (drought or well-watered, with or without host). ANOVA was performed to detect significant differences among the groups, with Tukey's post hoc test used to compare the estimated marginal means. Notice that in most cases the conditioning treatments did not affect the relative abundance of bacterial markers compared to the baseline soils.

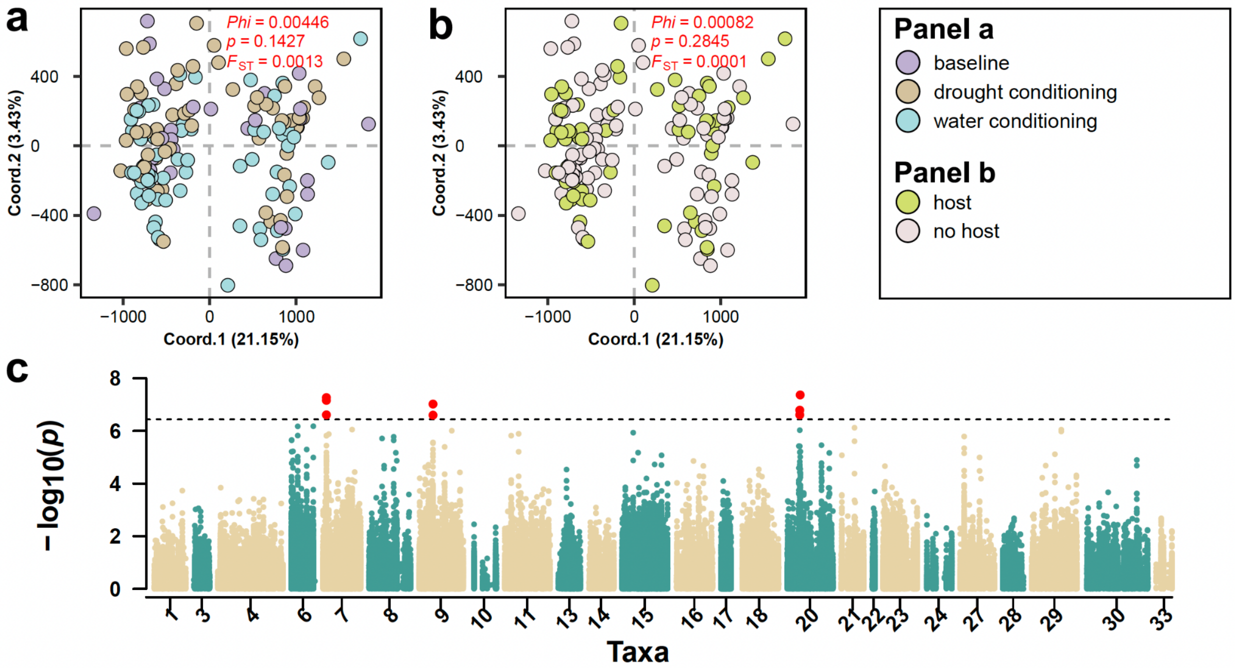


**Supplementary Fig. 3.** Genetic variation among bacterial lineages driven by long-term precipitation legacy remains stable despite acute drought/water perturbations and host presence/absence during the five-month-long conditioning phase. Filtered shotgun metagenomic reads were aligned to the reference genomes of 33 selected taxa, including 15 identified bacterial markers plus 18 additional abundant and prevalent taxa. Genetic distances between these bacterial lineages were calculated based on identified single nucleotide polymorphisms (SNPs). Principal Coordinate Analysis (PCoA) plots were generated and coloured to depict **a.** acute drought and well-watered treatments, and **b.** host and no-host treatments during the conditioning phase. Phi values, derived from Analysis of Molecular Variance (AMOVA), indicate the genetic variance between groups, while the p-value reflects the significance of the AMOVA results based on permutation testing. Fixation index ($F_{ST}$) values were used to measure the degree of genetic differentiation between groups. The variance explained by each axis is displayed on the PCoA plots. **c.** A Manhattan plot illustrated significant SNPs linked to the high-precipitation and low-precipitation soil legacies, derived from the genetic-environment association (GEA) analysis. This analysis was conducted using a general linear model and significant associations were identified using the permutation method. In the Manhattan plot, the x-axis represents the SNP positions along the genomes of the selected bacterial species, while the y-axis displays the $-\log_{10}$ p-values from the association model. A horizontal line represents the statistical significance threshold, determined by the permutation test. SNPs exceeding this threshold (highlighted in red) were significantly associated with soil legacy effects. Bacterial taxa with fewer than 1,000 high-quality biallelic SNPs after filtering, or with no significant SNPs detected from the GEA analysis, are excluded from the plot.

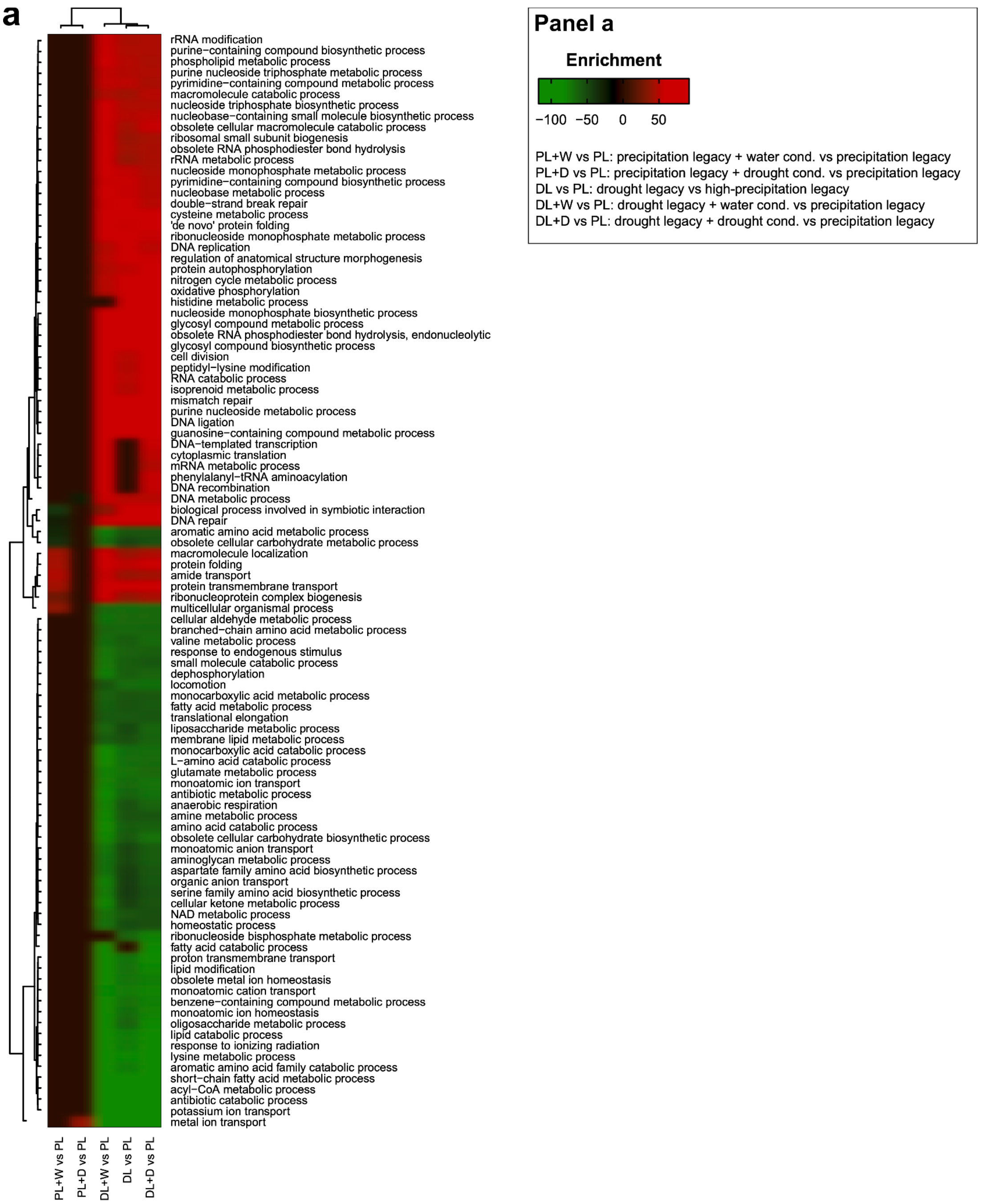


**Supplementary Fig. 4. Precipitation legacies shape soil microbial functional potential and remain stable under short-term water perturbations. a.** The heatmap illustrates enriched or depleted Gene Ontology (GO) categories in soils with high-precipitation legacies (PL) or low-precipitation legacies (DL), subjected to drought (+D) or well-watered (+W) treatments, in comparison to soils with the high-precipitation soil baseline (PL). DL indicates the baseline (pre-conditioning) low-precipitation-legacy soil. Gene enrichment analysis was conducted using a generalized linear model, followed by GO classification. Significantly enriched and depleted GO categories were identified using an adjusted p-value threshold of $q < 0.05$. Colours (red for enrichment, green for depletion) represent enrichment scores, calculated based on square root-transformed delta rank values of the GO categories. The heatmap is clustered by soil treatments and GO terms.

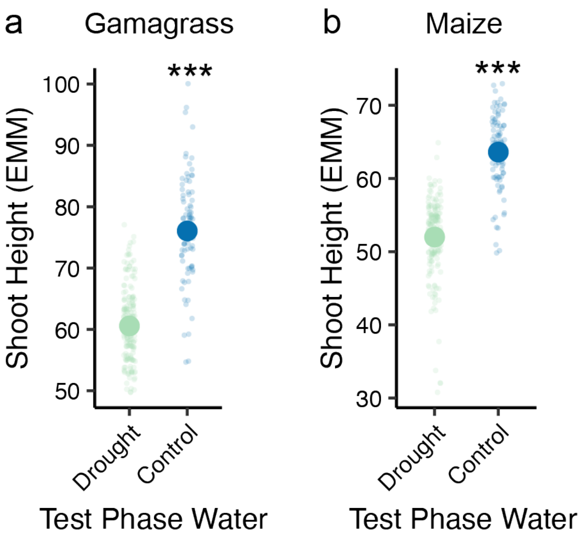


**Supplementary Fig. 5. Test phase gamagrass and maize drought treatments effectively reduce plant growth**. Test phase drought treatment significantly reduced shoot height (cm) in (**a**) gamagrass and (**b**) maize. Large points represent estimated marginal means, while small points show individual plant heights.

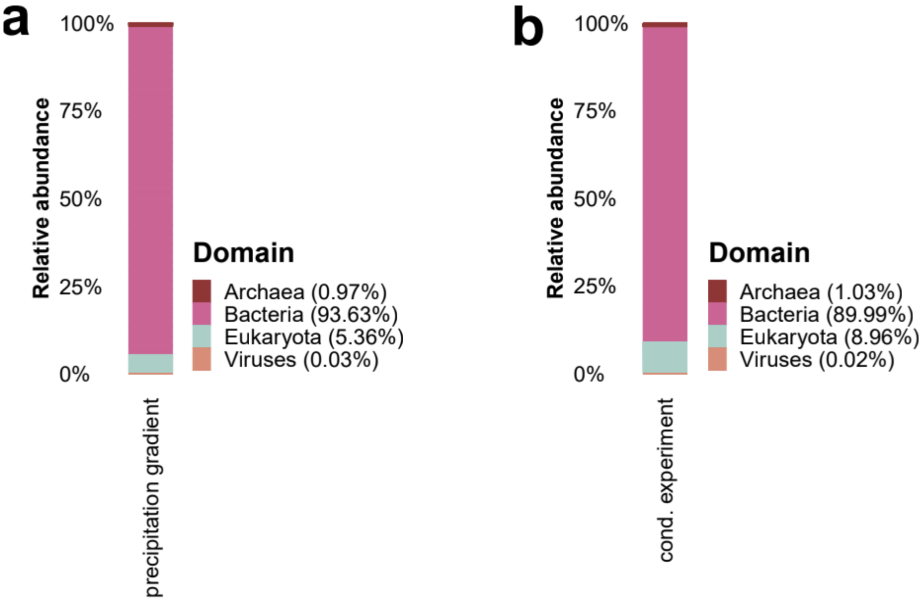


**Supplementary Fig. 6. Dominance of bacterial sequences in soil metagenomes.** Metagenomic profiles were dominated by bacterial sequences, which accounted for the vast majority of reads in both (**a**) the original soils sampled across a precipitation gradient and (**b**) the soils from the conditioning experiment. In contrast, archaeal, eukaryotic, and viral sequences constituted only minor fractions of the total reads.

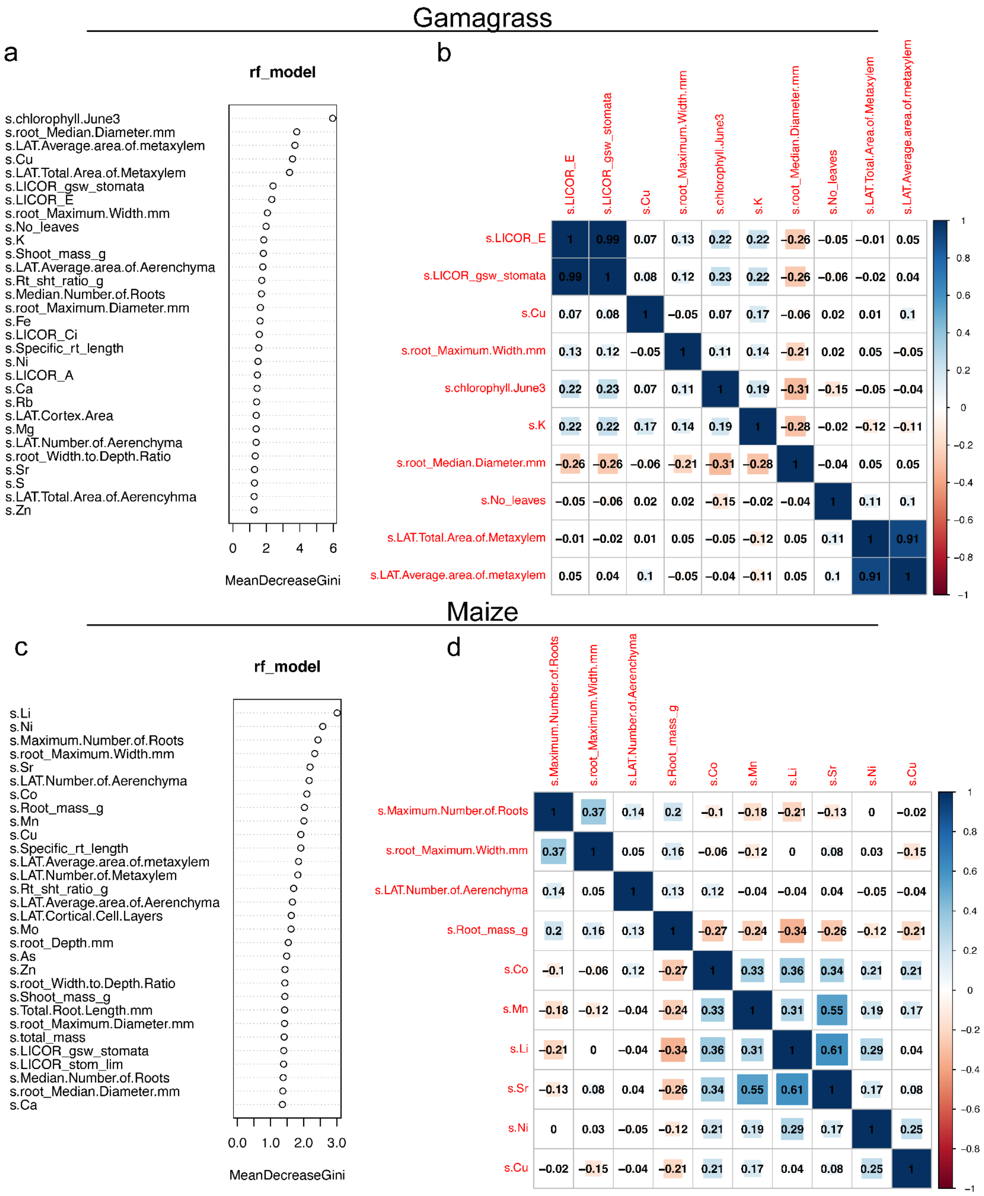


**Supplementary Fig. 7. Plant trait feature selection. a-b.** Gamagrass and **c-d.** maize traits important in explaining legacy effects were ranked using a random forest model. The top ten traits were tested for significant correlations (≥ 0.7).

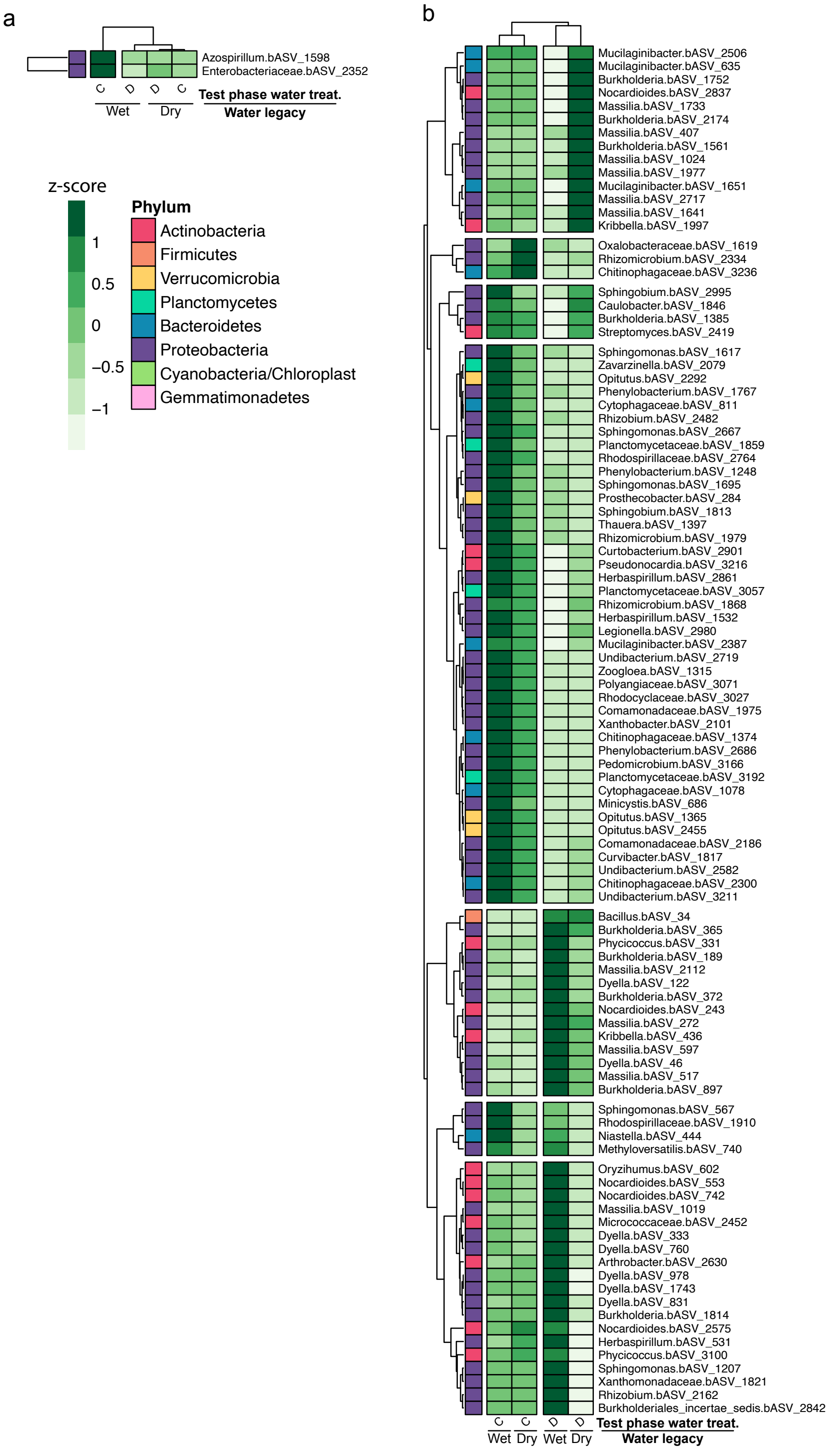


**Supplementary Fig. 8. Test phase water and legacy have a large impact on ASV abundances in maize, but not gamagrass. a**. Gamagrass root bacterial and **b.** maize root bacterial centered-log ratio transformed ASV counts were individually fit to a linear model to identify ASVs that were differentially abundant across test phase water treatments in the context of precipitation legacy. Dark green represents ASV enrichment and light green to white represents depletion. Multi-colored squares indicate phylum-level taxonomic assignments, which do not have an obvious phylogenetic pattern.

### Supplementary Table Legends

**Supplementary Table S1. List of soil collection sites.** The soils used in this work were collected from these regions. The table also shows the coordinates associated with these regions, the state, and country.

**Supplementary Table S2. Enriched KEGG reactions across the precipitation gradient (Low vs. High precipitation) in Kansas.** Positive enrichment scores (“Low_vs_high_precipitation”) indicate that the KEGG reaction was relatively more abundant in soil metagenomes from low-precipitation sites than in high-precipitation sites. Enrichment scores are the cumulative log2-fold changes of all differentially abundant genes within the same functional KEGG category, as determined using DESeq2.

**Supplementary Table S3. Significant SNPs linked to precipitation gradient, derived from the genetic-environment association (GEA) analysis.** Results of GEA based on metagenomic data from the original field-collected soils. This table specifies the genomic locations of SNPs, the reference and alternative alleles, GenBank accession numbers, and annotations when available. The ‘Effect’ column represents the estimated additive effect of the alternate allele (ALT) from the association model on the binary trait (0 = drought, 1 = precipitation). A positive effect indicates that the ALT allele is associated with an increased probability of precipitation conditions, while a negative effect indicates that the ALT allele is associated with an increased probability of drought conditions.

**Supplementary Table S4. List of bacterial markers of soil water legacy identified in this work using soils from Kansas.** In addition to the bacterial taxa ID (taxID), the table also shows taxonomical attributes (Family, Genus, label Family, and Label genus) of the bacterial markers identified. The column "DirectionEnrichment" shows the direction of the marker enrichment: “High to low” indicates that the marker taxon is enriched in the soils from high-precipitation sites relative to soils from low-precipitation sites, and vice versa.

**Supplementary Table S5. Significant SNPs linked to long-term water and drought legacies even after five months of experimental perturbation.** Results of GEA based on metagenomic data from soils that had undergone five months of experimental perturbation (acute drought or control conditions, with or without a host). This table specifies the genomic locations of SNPs, the reference and alternative alleles, GenBank accession numbers, and annotations when available. The ‘Effect’ column represents the estimated additive effect of the alternate allele (ALT) from the association model on the binary trait (0 = drought, 1 = precipitation). A positive effect indicates that the ALT allele is associated with an increased probability of precipitation conditions, while a negative effect indicates that the ALT allele is associated with an increased probability of drought conditions.

**Supplementary Table S6. Enriched GO categories across the precipitation gradient after drought and watering conditioning (metagenome).** At the end of the conditioning phase, 270 GO categories were significantly enriched or depleted in at least one treatment group or in the baseline low-precipitation-legacy soils, relative to the high-precipitation-legacy baseline soils. Enrichments and depletions were much more common in the low-precipitation-legacy soil group, regardless of whether the soils experienced the drought treatment (“+ drought cond.”) or the well-watered treatment (“+ water cond.”) during the conditioning phase. “NA” indicates that the GO category was neither enriched nor depleted.

**Supplementary Table S7. Overlapping enriched GO categories across the precipitation gradient and conditioning treatments.** By the end of the conditioning phase, most of the GO categories that distinguished the original field-collected soils from low-precipitation *vs.* high-precipitation sites (see Extended Data Fig. 2g) retained the same pattern of enrichment/depletion, regardless of whether the soils had experienced the drought treatment ("+ drought cond.") or the well-watered treatment ("+ water cond.") during the conditioning phase. "NA" indicates that the GO category was neither enriched nor depleted.

**Supplementary Table S8. Enriched GO categories across the precipitation gradient after drought and watering conditioning (metatranscriptome).** Metatranscriptomic analysis of pre-conditioning (baseline) and post-conditioning (5 months of drought ["+ drought cond."] or well-watered conditions ["+ water cond."]) soils revealed GO categories that were differentially abundant relative to the high-precipitation-legacy baseline soils. "NA" indicates that the GO category was neither enriched nor depleted.

**Supplementary Table S9. Enriched KEGG reactions across the precipitation gradient after drought and watering conditioning (metatranscriptome).** Metatranscriptomic analysis of pre-conditioning (baseline) and post-conditioning (5 months of drought ["+ drought cond."] or well-watered conditions ["+ water cond."]) soils revealed KEGG reactions that were differentially abundant relative to the high-precipitation-legacy baseline soils.

**Supplementary Table S10. Differentially expressed genes in the roots of gamagrass and maize inoculated with low-precipitation-legacy and high-precipitation-legacy microbiota during the test phase.** Tab one provides information for interpreting the column names and the four other tabs. Tab two lists gamagrass genes that are significantly up or down-regulated based on the main effects of microbial inoculum legacy. Tab three lists gamagrass genes that are significantly up or down-regulated based on the interaction of test phase water treatment and microbial inoculum legacy. Tab four lists maize genes that are significantly up or down-regulated based on the main effects of microbial inoculum legacy. Tab five lists maize genes that are significantly up or down-regulated based on the interaction of test phase water treatment and microbial inoculum legacy.

**Supplementary Table S11. Test phase plant phenotypic data and microbiome metadata.** Tab one provides information for interpreting the column names. Tab two provides the measurements for all plants used in the plant phenotypic analyses, including both Gamagrass and maize. Tab three provides the measurements for the plant samples used in the root bacterial microbiome data analyses, with sequencing data that passed all qualify checks and filtering.

**Supplementary Table S12. Differentially abundant test phase gamagrass and maize root microbiome ASVs.** Tab one provides information for interpreting the column names. Tab two lists the gamagrass root microbiome ASVs, including their full taxonomic assignment, that were differentially abundant based on inoculum precipitation legacy in the context of test phase water treatment. Tab three lists the maize root microbiome ASVs, including their full taxonomic assignment, that were differentially abundant based on inoculum precipitation legacy in the context of test phase water treatment.

### Supplementary Note 1

We identified four sets of orthologous genes whose transcription patterns were sensitive to microbiota legacy in both maize and *T. dactyloides*, but none showed congruent drought responses. In maize, a peroxidase encoding gene, Zm00001eb076200, reversed its drought response depending on inoculum, while its *T. dactyloides* ortholog, Td00002ba025285, was up-regulated 4-fold by drought only  when inoculated with wet-legacy microbiota. The maize gene Zm00001eb077640, (L-allo-threonine aldolase) also showed a reversed drought response, whereas its ortholog Td00002ba026366 was consistently up-regulated, especially in plants inoculated with dry-legacy biota. The pathogenesis-related protein-like gene Zm00001eb150050, which has been linked to both biotic and abiotic stress responses, was up-regulated 7-fold in maize under drought, but only in plants inoculated with wet-legacy microbiota; however, its *T. dactyloides* orthologs were down-regulated 8-fold in response to low-precipitation *vs.* high-precipitation legacy inoculum, independent of watering. Finally, the tryptophan synthase encoding gene Zm00001eb301540 was up-regulated 1.7-fold in maize inoculated with wet-legacy microbiota, while; its ortholog Td00002ba012570 was down-regulated under the same conditions.
